# CHROMATIX: computing the functional landscape of many-body chromatin interactions in transcriptionally active loci from deconvolved single-cells

**DOI:** 10.1101/854190

**Authors:** Alan Perez-Rathke, Qiu Sun, Boshen Wang, Valentina Boeva, Zhifeng Shao, Jie Liang

**Affiliations:** University of Illinois at Chicago; Shanghai Jiao Tong University; U1016 INSERM, Institut Cochin, CNRS UMR 8104 / Universite Paris Descartes UMR-S1016

## Abstract

Chromatin interactions are important for gene regulation and cellular specialization. Emerging evidence suggests many-body spatial interactions can play important roles in condensing super-enhancer regions into a cohesive transcriptional apparatus. Chromosome conformation studies using Hi-C are limited to pairwise, population-averaged interactions; therefore, not suitable for direct assessment of many-body interactions. We describe a computational model, CHROMATIX, that reconstructs structural ensembles based on Hi-C data and identifies significant many-body interactions. For a diverse set of highly-active transcriptional loci with at least 2 super-enhancers, we detail the many-body functional landscape and show DNase-accessibility, POLR2A binding, and decreased H3K27me3 are predictive of interaction-enriched regions.

## Background

Chromosome folding and nuclear organization play essential roles in fundamental processes such as regulation of gene expression (1; 2) and cellular specialization (3; 4). A wealth of information on chromatin organization has been gained through studies based on chromosome conformation capture techniques such as Hi-C (5; 6; 7; 8), which measure pairwise, proximity interactions between chromatin regions that are averaged over a population of cells (6; 9). There is now growing evidence that multi-valent interactions play important roles in formation of phase-separated and highly dense, functional chromatin assemblies in super-enhancers (SEs) (10; 11); however, it is difficult to detect and quantify many-body (≥ 3) interactions from pairwise and averaged Hi-C measurements.

Several experimental techniques have been developed to detect putative many-body chromatin interactions. These include single-cell Hi-C (12; 13; 14), Dip-C (15; 16), Tri-C (2), GAM (17), and SPRITE (18). However, there are limitations with these techniques. For example, while single-cell Hi-C permits detection of instances of many-body interactions in individual cells, it often has low genomic coverage (19); GAM and SPRITE do not readily distinguish direct from indirect many-body chromatin interactions due to ancillary coupling effects (17; 18). Overall, our current knowledge of many-body chromatin interactions and their functional roles in chromatin condensation is limited.

With the extensive availability of population-averaged Hi-C data for many biological systems, we ask whether it is possible to gain insight into functionally important many-body spatial interactions from these high-quality, high-resolution measurements. While no computational method is currently available, we hypothesize that 3-D polymer modeling can be used to overcome the limitations of population-averaged, pairwise Hi-C measurements. However, there are a number of significant technical challenges. These include: *i*) deconvolving the population-averaged and pairwise Hi-C contact frequencies into an underlying ensemble of single-cell 3-D chromatin folds, such that instances of many-body interactions in single cells are collectively consistent with the input Hi-C; and *ii*) distinguishing *specific* (*i*.*e*, highly non-random) many-body interactions from non-specific interactions which are largely due to effects of linear genomic proximity (20) and nuclear confinement (21; 22; 23).

Modeling of 3-D chromatin structure allows for detailed analysis of nuclear organization patterns and can detect spatially interacting regions (21; 22; 23; 24; 25; 26; 27; 28; 29; 30; 31; 32; 33; 34). There are many well-developed physical models for chromatin folding, including the Strings and Binders Switch (SBS) model (24), the Minimal Chromatin Model (MiChroM) (26; 28), and the n-Constrained Self-Avoiding Chromatin (nCSAC) model (21; 22). The nCSAC approach folds polymers under the influence of predicted specific pairwise interactions obtained after controlling for effects of nuclear confinement. The SBS and MiChroM models follow block copolymer approaches (29; 30), in which chromatin regions are assigned different affinities for each other based on their corresponding *types*. In SBS, chromatin types are defined by their affinity to Brownian binder particles which facilitate bridging of multiple chromatin sites up to a specified valency. In MiChroM, chromatin types and affinities are based on clustering of epigenetic markers, followed by maximum-entropy optimization of the resulting energy function. SBS and MiChroM can reproduce important physical phenomena such as the dynamics of chromatin condensation leading to phase separation; however, no methods for calling specific many-body chromatin interactions based on these models have been reported yet.

Several computational methods have been developed to detect specific pairwise chromatin interactions present within Hi-C datasets (20). These include the negative binomial model of Jin *et al*. (35), the non-parametric spline approach of Fit-Hi-C (36), the binomial model of GOTHiC (37), the local neighborhood loop-calling approach of HiCCUPS (9), and the hidden Markov random field model of Xu *et al*. (38). These methods rely on the empirical Hi-C for estimation of a background model that is then used to assess the significance of each pairwise chromatin contact; hence, these approaches may contain intrinsic bias as the observed Hi-C data is being used for construction of its own null hypothesis test. In addition, these methods lack a 3-D folding model and therefore cannot assess the significance of many-body (≥ 3) chromatin spatial interactions.

In this work, we describe CHROMATIX (CHROMatin mIXture), a new computational approach for detecting *specific* many-body interactions from population averaged Hi-C data. We focus on uncovering occurrences where 3-, 4-, or more genomic regions all spatially co-locate to within a defined Euclidean distance threshold. We further require that these occurrences do not arise from simple physical effects of monomer connectivity, excluded volume, and spatial confinement; we refer to these as *specific many-body interactions*.

We extend the nCSAC (21; 22) folding method which allows for nearly unbiased construction of random polymer chains to serve as a null model completely decoupled from the Hi-C data. By further integrating extensive polymer simulations under a Bayesian generative framework (39), we resolve complex dependencies among chromatin contacts and deconvolve population Hi-C data into the most likely single-cell contact states. These contact states are then folded to produce a 3-D structural ensemble consistent with the measured Hi-C. We achieve our results through a novel deep-sampling algorithm called *fractal Monte Carlo*, which can generate 3-D polymer ensembles with improved structural diversity and target distribution enrichment (see Supplementary Information).

To study highly non-random and direct higher-order interactions among super-enhancers, enhancers, and promoter regions, we apply our method to a diverse set of 39 highly transcriptionally-active loci in the GM12878 mammalian cell line; specifically, all TAD-bounded (40; 41) loci (<2 MB), each with at least 2 super-enhancers (1; 3; 4) showing evidence of possible super-enhancer condensation (see Supplementary Information, Table S1) (18). We detect specific many-body interactions in each of these loci, summarize the landscape of functional associations among participating regions, and report common biological factors predictive of interaction enrichment.

## Results

### Model for chromatin folding

We independently modeled the 39 genomic loci, ranging in size from 480 KB to 1.94 MB, each as a connected, self-avoiding polymer chain where monomer beads represent 5 KB of 11 nm chromatin fiber (42; 43). Loci lengths in base pairs are from the corresponding TAD (arrowhead) boundaries as reported in Rao *et al*. (9) (see Supplementary Information). Each locus was simulated under a confining sphere based on the GM12878 nuclear diameter reported in Sanborn *et al*. (44) and scaled to preserve a constant base pair density (bp*/*nm^3^).

#### Identifying specific interactions from Hi-C data

The CHROMATIX modeling pipeline is illustrated in Figure 2. Briefly, we first identify pairwise *specific contacts* from measured Hi-C interaction frequencies by following the general approach of Gürsoy *et al*. (21); namely, we identify chromatin interactions with Hi-C frequencies unlikely to be observed under a uniform random folding environment (45). We extend the approach of Gürsoy *et al*. by using the method of fractal Monte Carlo weight enrichment (see Supplementary Information) to uniform randomly sample an ensemble of ∼400, 000 3-D polymer conformations (see Figure 2a, and Figure S1 for examples of random polymers). These polymers are used as a null ensemble for identifying significant Hi-C interactions that are unlikely to be formed due to random chance (Figure 2b). The assumption of spherical confinement makes this null model more stringent in calling specific interactions as discussed in (22), although our tool supports other confinement models (e.g. ellipsoid). Details on *p*-value calculations can be found in Methods.

**Figure 1:**
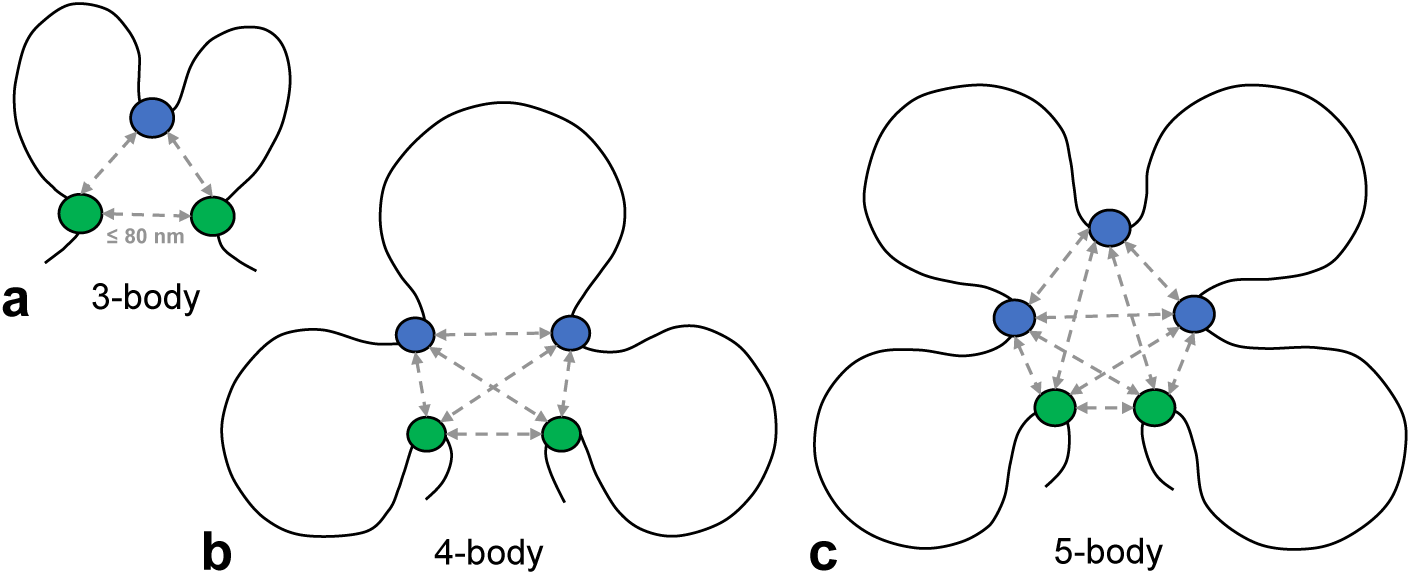
Diagrams of 3-, 4-, and 5-body chromatin interactions. **a, b, c** Diagrams illustrating 3-, 4-, and 5-body chromatin interactions respectively (green and blue dots). Grey arrows represent spatial Euclidean distances within 80 nm (46). The *principal loop* is the longest loop (in bp) among chromatin regions forming a many-body (≥ 3) interaction, and genomic regions serving as anchors of principal loops are represented by green dots.

**Figure 2:**
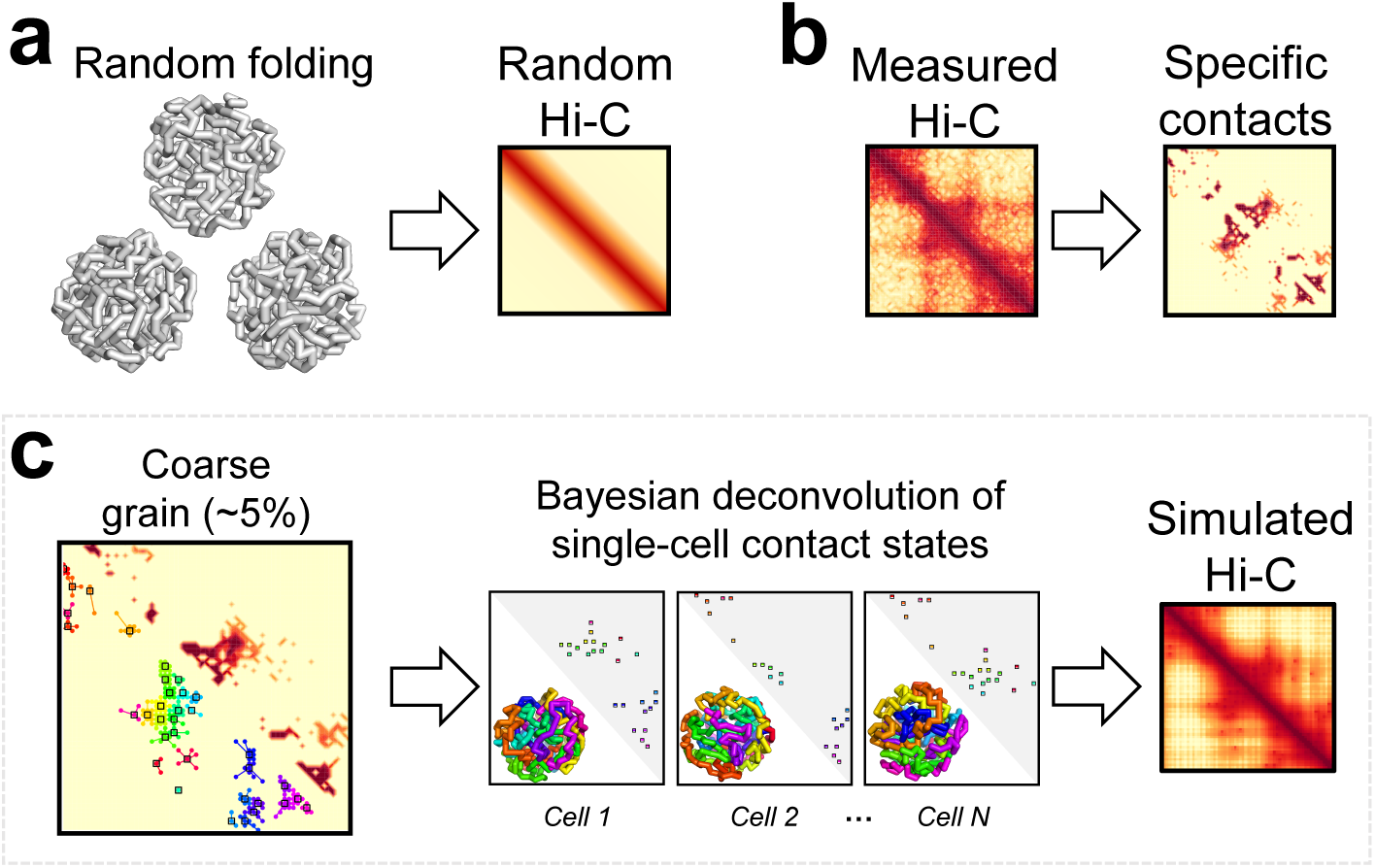
CHROMATIX modeling pipeline. **a** Random polymers are generated using fractal Monte Carlo sampling. **b** Specific contacts are identified from measured Hi-C using a random polymer ensemble as the null distribution (21). **c** Specific contacts are coarse-grained and single-cell contact states are deconvolved then folded to generate simulated Hi-C (see Supplementary Information).

#### Identifying a minimal set of sufficient interactions

We conjecture that not all specific interactions are required to produce the observed Hi-C chromatin folding patterns (22; 46). To identify a minimal set of interactions that are sufficient to drive chromatin polymers into a folded ensemble that exhibit the observed Hi-C frequencies, we retain roughly 5% of the identified specific contact interactions using clustering (47; 48) (see Supplementary Information for more details). We call this procedure *coarse-graining* of the specific contacts (Figure 2c); coarse-graining also regularizes our model to help prevent overfitting.

#### Single-cell contact state deconvolution

Many-body interactions occur probabilistically in individual cells. To reconstruct the 3-D chromatin polymer for each cell of a modeled population, we must predict which contacts among the set of minimally sufficient interactions are co-occurring within each individual cell. We call these co-occurring interactions the *single-cell contact states* (Figure 2c). Once a single-cell contact state is properly generated, we then construct a set of 3-D chromatin polymers that are all consistent with this single-cell contact state. By generating a large number of single-cell contact states, we can obtain an ensemble of 3-D chromatin polymers which accurately reproduce the observed population Hi-C measurements. Structural analysis of the ensemble of single-cell chromatin conformations can then reveal specific spatial many-body interactions.

The key to properly generating single-cell contact states is to account for dependencies among chromatin interactions; namely, how certain physical interactions may cooperatively induce formation of other interactions due to polymer folding. These dependencies are identified by *in silico* knock-in perturbation studies, where differential contact probabilities are assessed between two ensembles of chromatin polymers, one with and another without the target contact knocked-in. A large number of possible dependencies are identified through these extensive polymer knock-in simulations (see Methods and Supplementary Information). Such simulations also identify geometrically infeasible contact combinations.

To properly deconvolve population Hi-C interactions into single-cell contact states, we adopt a Bayesian generative approach. The dependencies and infeasible geometries among contacts are incorporated as a Bayesian prior. This physically-based prior along with the measured Hi-C data enables efficient Bayesian inference over the posterior distribution of single-cell contact states. Specifically, we use Gibbs sampling for this inference (see Supplementary Information). For efficiency, we first coarse-grain the called specific Hi-C interactions before carrying out knock-in simulations and Gibbs sampling. Only about 5% of the specific interactions are retained, which substantially reduces the computational cost, making this approach highly practical.

#### Reconstructing 3-D chromatin folds

For a given deconvolved single-cell state of chromatin contacts, we uniformly sample among the set of 3-D folds satisfying the spatial proximity interactions specified by the single-cell state. Specifically, we sample from the uniform distribution of chromatin chains conditioned on the deconvolved contact state of each cell, where two regions are spatially interacting if their Euclidean distance is ≤ 80 nm (46). This procedure is repeated for each sampled single-cell contact state (see Figure S2 for examples of sampled chromatin polymers).

Overall, we aggregate ∼50 folds per single-cell to generate an ensemble of 25,000 3-D chromatin polymers at each of the 39 modeled genomic loci. These sampled conformations form the reconstructed ensemble of intrinsic 3-D folds underlying the population-aggregated Hi-C.

### Simulated 3-D polymer ensembles strongly correlate with Hi-C measurements

We find the chromatin interaction frequencies from the computed 3-D polymer ensembles (called *simulated Hi-C*) to strongly correlate with measured Hi-C frequencies (Figure 3). The Pearson correlations between the simulated and measured Hi-C frequencies have approximate *mean* and *standard error of the mean* (SEM) of 0.970 ± 0.003 over the 39 modeled genomic loci (see details in Supplementary Information). Here, correlations were computed at 5 KB resolution after the measured Hi-C counts were quantile normalized according to the uniform randomly sampled polymer ensemble (Figure 2a). This approach is motivated by similar methods for comparing gene expression microarrays (49); it allows direct comparison between simulated ensemble frequencies and measured Hi-C counts. To exclude proximity effects owing to genomic distance, we further remove the first two diagonals from the Hi-C heatmaps; namely, all Hi-C frequencies within 10 KB are excluded. The simulated and measured Hi-C data again exhibit excellent Pearson correlations, with an approximate mean and SEM of 0.96 ± 0.003; more details on simulations of the 39 loci are shown in Figure S3. We also computed the *distance corrected* Pearson correlations (50) and obtained a mean and SEM of 0.64 ± 0.02 (more details in Table S1 and Figure S4). These results indicate that our 3-D ensembles are consistent with the measured Hi-C interaction patterns.

**Figure 3:**
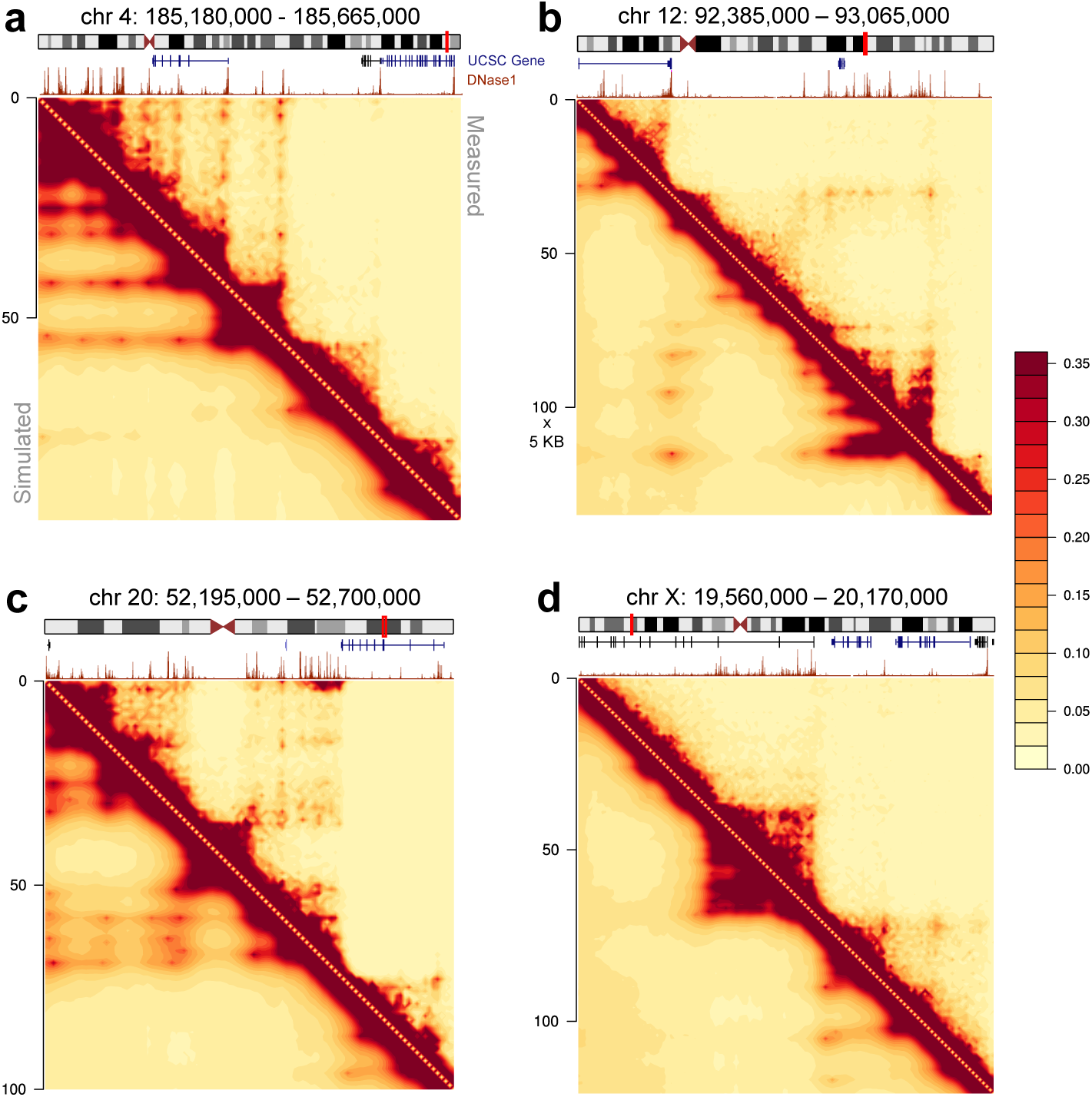
CHROMATIX Hi-C reconstruction. 4 representative genomic regions (**a, b, c, d**), with the measured Hi-C (9) on the upper triangle and the simulated Hi-C from aggregation of 3-D polymer folds on the lower triangle. The Pearson correlations between simulated and measured Hi-C for all 39 modeled genomic loci have approximate mean of 0:96 ± 0:003 SEM, after removal of the first 2 diagonals. DNase data are from ENCODE (52; 53) (ENCSR000EMT) with corresponding signal, gene, and chromosome diagrams from UCSC genome browser (74; 75). All heatmaps are in units of 5 KB.

### Reconstructed single-cell chromatin structures

We have compared our single-cell chromatin models with publicly-available single-cell Dip-C data for GM12878 (15). For each cell in the Dip-C ensemble, we identified the corresponding CHROMATIX cell with maximal overlap of contacts. Fig 4 shows the overall pattern of agreement and examples of individual single-cells. In general, CHROMATIX single-cell models contain more contacts (grey regions in Fig 4a-c) than that of Dip-C, but there is overall good agreement, with many long-range contacts appearing in both Dip-C and CHROMATIX single cells (Fig 4a-c). The median overlap coefficient is ∼ 65% for the *n* = 976 cell-loci.

**Figure 4:**
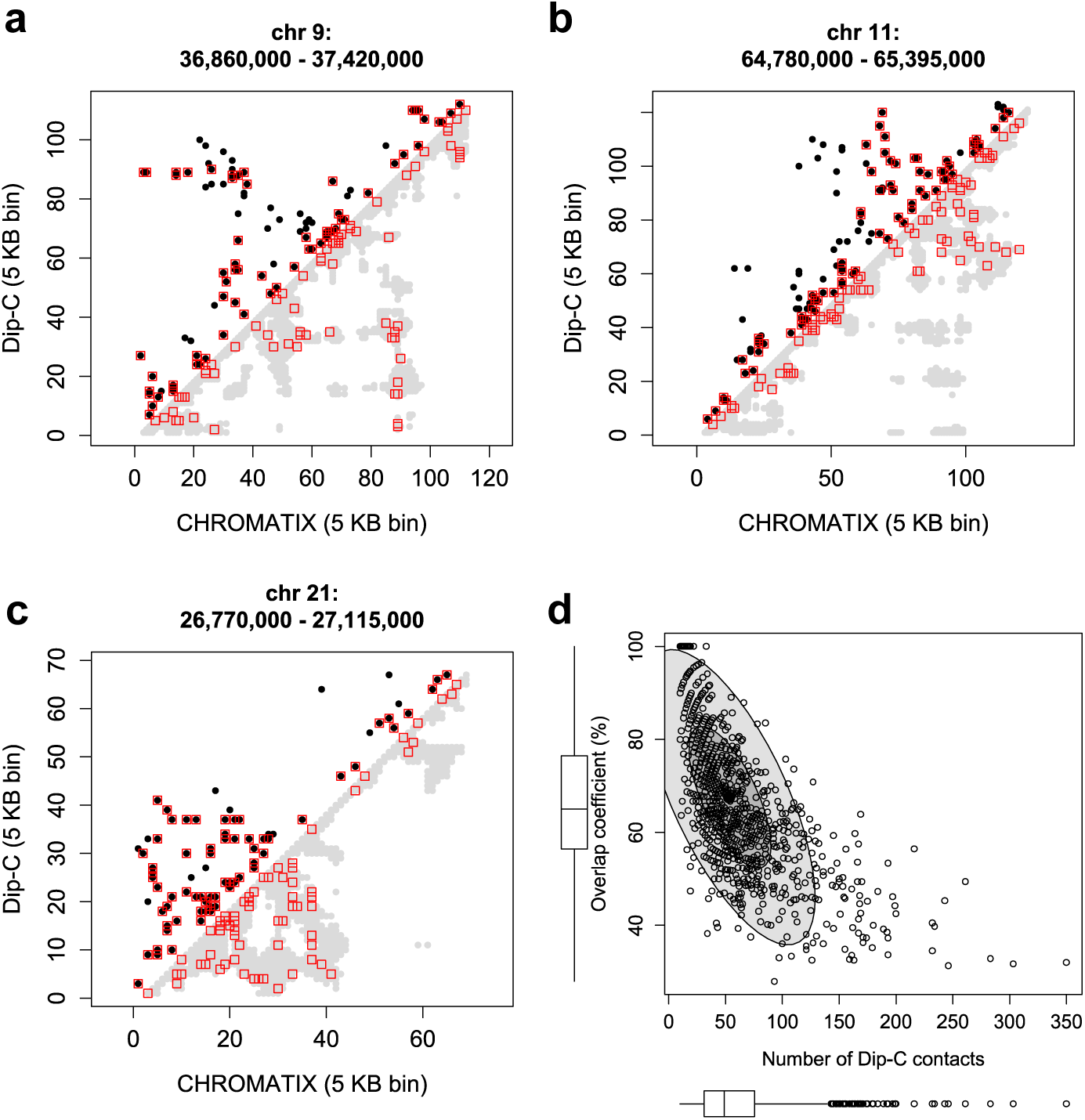
Comparison with Dip-C single-cell data (GSE117874) (15). **a, b, c** Plots of pairwise contacts between representative Dip-C cells (upper triangle, black dots) and the corresponding CHROMATIX cells (lower triangle, grey dots) of maximal overlap coefficient. Contacts present in both models are outlined in red. **d** Scatter plot of maximal overlap coefficient (*Y*-axis) versus number of contacts present within each Dip-C model (*X*-axis) of single-cell chromatin at different loci (*n* = 976). The horizontal boxplot shows the distribution of Dip-C contacts per cell (median ∼ 50). The vertical boxplot shows the distribution of maximal overlap coefficients between the Dip-C and CHROMATIX ensembles (median ∼ 65%). The inner and outer ellipses contain 5% and 95% of the single-cells, respectively. More details can be found in Supplementary Information.

### Analysis of single-cell chromatin domains

Motivated by single-cell optical imaging studies of Bintu *et al* (51), we examined the 3-D chromatin structures at locus chrX:19,560,000–20,170,000 to assess if single-cell domains are present (Fig 5). Our key findings are similar to that of (51), even though the cells we modeled are of different cell lineage. Specifically, diverse patterns of chromatin contacts are seen in reconstructed chromatin folds of single cells: domain-like patterns appear among single-cell distance plots (Fig 5c), which resemble the domains in the mean distance plots (Fig 5a). Similar to (51), there are many instances where the domain patterns are less clear. Furthermore, there is non-zero probability of forming domain boundaries at all locations of the locus, and the precise boundaries shift from cell to cell. However, we observe similarly consistent boundary strengths at similar genomic coordinates (Fig 5b and 5d).

**Figure 5:**
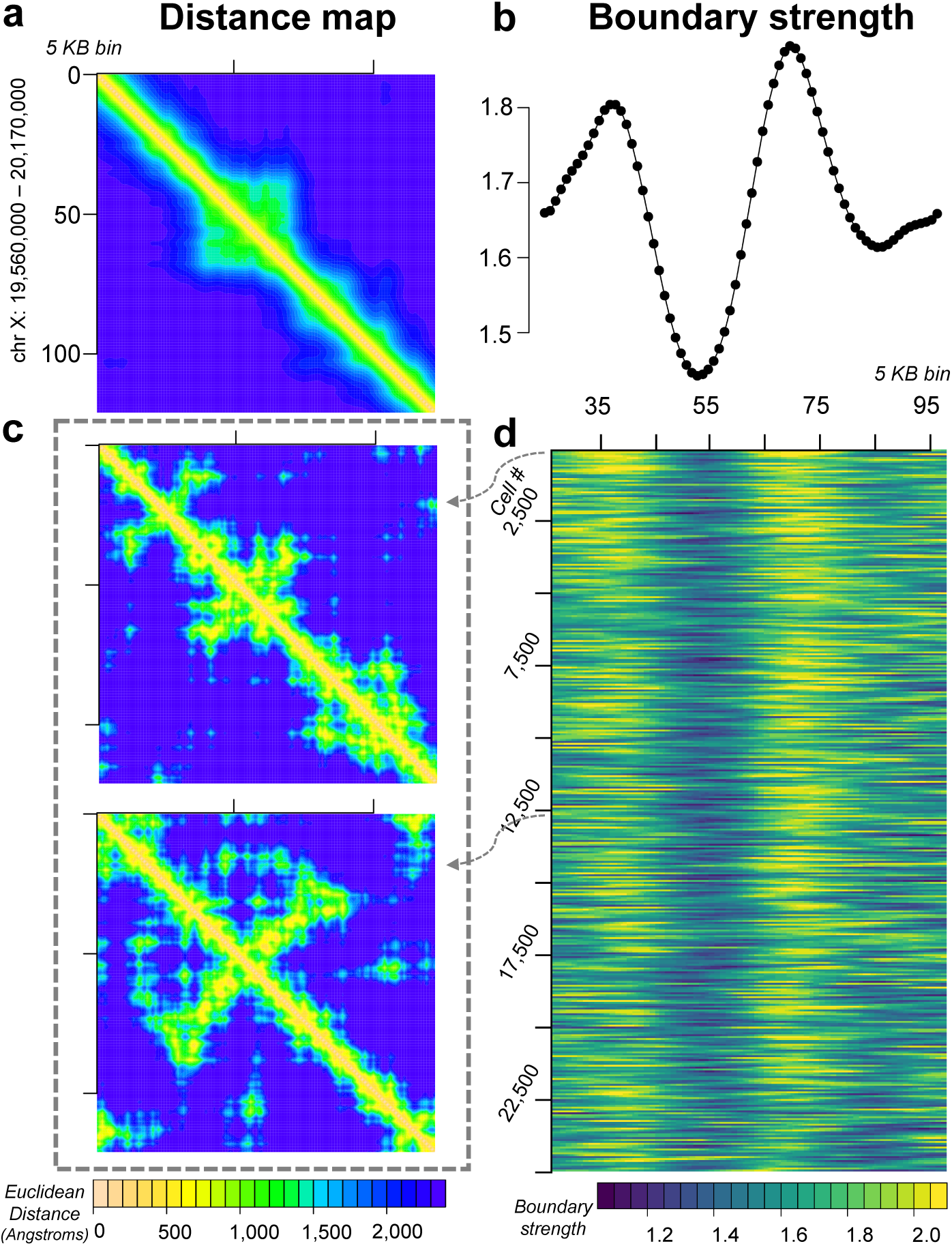
Reconstructed ensemble of 25,000 single-cell chromatin structures of the locus chr X: 19,560,000 - 20,170,000 at 5 KB resolution. **a** Heatmap of mean pairwise Euclidean distance in Å. Corresponding Hi-C heatmaps (experimental and simulated) can be seen in Figure 3d. **b** Boundary strength of mean pairwise distances computed following (51) at each 5 KB bin. **c** Single-cell pairwise distance heatmaps for two representative cells. **d** Heatmap of single-cell boundary strengths, each row is the boundary strength curve of an individual cell among the 25,000 cell ensemble.

### 3-body complexes, *maximal* many-body complexes, and principal loops

For each of the 39 loci, we are interested in fully-interacting *3-body complexes*, which are formed by three genomic regions where the Euclidean spatial distances among all pairs of regions are ≤ 80 nm (46). These 3-body complexes may be a component of a larger (*k* > 3) fully-interacting complex.

We are also interested in *maximal many-body complexes* which are formed by *k* ≥ 3 genomic regions, where all pairwise Euclidean distances are ≤ 80 nm, and cannot be extended to include additional regions while satisfying the distance requirement. We characterize a maximal 3-, 4-, 5-, or higher-order *k*-body complex by its *principal loop*, which is the longest genomic span in base pairs within each *k*-body complex (Figure 1).

Furthermore, we are interested in *specific* 3-body complexes and *specific* maximal many-body complexes, whose spatial interaction frequencies are unlikely to be observed under a uniform random folding environment (see Methods).

### SPRITE concordance

We compared our predicted *3-bodies* and *maximal many-body principal loops*, generated from population-averaged Hi-C, with publicly available SPRITE (split-pool recognition of interactions by tag extension) data for GM12878 cells (18). The SPRITE technique captures *clusters* of co-occurring chromatin interactions. However, SPRITE does not distinguish *direct* from *indirect* cross-linking among chromatin fragments (18) - *i*.*e*, some chromatin regions present within a SPRITE cluster may not have direct spatial interactions, but, rather, may have been co-captured through a sequence of cross-links among spatially proximal regions that could extend to distances beyond the cross-linking threshold. Nevertheless, a high proportion of our predicted many-body interactions were also observed to co-occur within a SPRITE cluster; we term this proportion the *found fraction*. Specifically, across all 39 modeled genomic loci, we saw fairly similar median found fractions for specific and non-specific 3-bodies (approximately 90% and 86% respectively) as well as for principal loops (both medians approximately 99%) at 5 KB resolution.

To adjust for bias due to genomic distance, we stratified principal loops of many-body complexes by base pair span and computed their respective SPRITE *coverage fractions, i*.*e*. proportion of SPRITE clusters containing the principal loop. Specifically, we computed the median SPRITE coverage fraction at each 5 KB genomic distance span for both *specific* and *non-specific* principal loops (Figure S5). We found the proportion of specific median coverage fractions exceeding the corresponding non-specific coverage was significantly elevated in 29 of 39 (∼74.4%) modeled genomic loci (FDR < 0.05, see Methods).

We performed a similar procedure for 3-body interactions, with stratification by both principal and minor (lowest bp span) loops. In this case, the proportion of specific median coverage fractions exceeding the corresponding non-specific coverage was significantly elevated in 25 of 39 (∼64.1%) modeled loci (FDR < 0.05, see Methods).

Overall, we find that after controlling for genomic distance, our many-body predictions are concordant with SPRITE clusters such that specific many-bodies generally exhibit elevated SPRITE coverage over the corresponding class of non-specific many bodies. More details can be found in the Supplementary Information.

### Specific 3-body complexes are enriched in direct interactions among functional genomic regions

Our 3-D chromatin ensembles contain rich structural information. Despite the strong effects of nuclear confinement and genomic connectivity that likely induce many bystander proximity ligations (Figure 2a) (21; 22), our model can identify *specific* many-body interactions. Figure 6 provides an overview of our findings for specific 3-body interactions across the 39 super-enhancer containing loci. While functional genomic regions (*i*.*e*., super-enhancers, enhancers, and promoters) participate in both specific and non-specific 3-body interactions, the proportion of interactions with no known functional associations is *markedly increased* for non-specific (33 ± 3% SEM, Figure 6a) compared to specific (19 ± 2% SEM, Figure 6c) 3-body interactions. Further, the medians of non-specific *vs*. specific 3-body interactions without functional associations (31% and 17% respectively) are significantly different (*p*-value = 4.5 × 10^−5^ by Mann-Whitney U test, Figure S6a).

**Figure 6:**
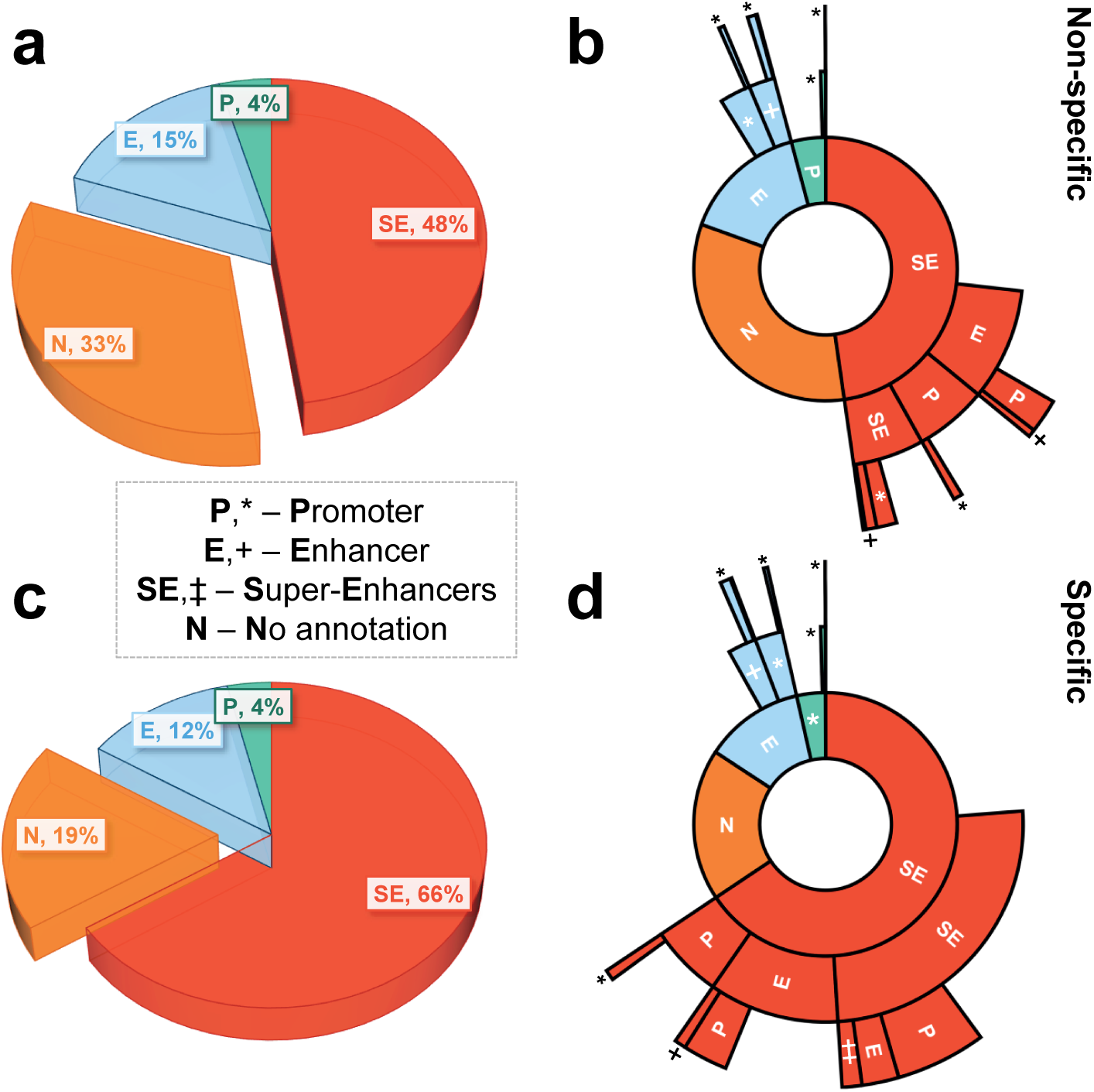
Functional landscape of 3-body chromatin interactions. Pie (**a,c**) and corresponding sunburst (**b,d**) charts for the proportion of specific (*bottom*) and non-specific (*top*) 3-body interactions involving the functional genomic regions of super-enhancer (SE), enhancer (E), and promoter (P). The innermost ring of the sunburst charts (**b,d**) are the same as the corresponding pie charts of (**a,c**), with outer rings representing the sub-fractions of interacting partners with SE, E, or P functional associations. Gaps in the sunburst charts represent the fractions of interacting partners with no known SE, E, or P annotation. Here, 3-body interactions are not required to be *maximal*, and can be part of a larger many-body complex where all regions are within 80 nm. Plots shown are the averages across all 39 modeled genomic loci.

### Functional landscape of specific 3-body complexes shows interactions among super-enhancers and promoters

The functional landscape of 3-body spatial interactions is shown in Figure 6b and 6d. We observe a higher proportion of specific 3-body interactions involving multiple (≥ 2) super-enhancers directly co-interacting with promoters, when compared to non-specific 3-body interactions (approximately 5.5 ± 0.6% SEM *vs*. 1.2 ± 0.3% SEM respectively, with *p*-value = 1 × 10^−8^ by Mann-Whitney U test on the corresponding medians of 4.5% and 0.8%, respectively, Figure S6b). Similarly, we observe a slightly higher proportion of specific 3-body interactions with at least 3 distinct super-enhancers relative to non-specific 3-body interactions (approximately 1.2±0.4% SEM *vs*. 0.2±0.1% SEM respectively at *p*-value = 8.4 × 10^−5^ by Mann-Whitney U test on the corresponding medians of 0.5% and 0.0% respectively, Figure S6c).

### Functional landscape of maximal 4- and 5-body complexes shows specific principal loops bridging super-enhancers

Our high-resolution 3-D chromatin ensembles also contain information on *maximal* higher-order many-body interactions. Figure 7 provides an overview of the functional landscape of maximal *k*-body complexes (*k* ≥ 3) among the 39 SE-associated loci. Here a maximal *k*-body complex is defined such that it cannot be extended to form a fully interacting *k* +1 or higher complex; this is unlike the 3-body complexes depicted in Figure 6, which may be part of still higher order (*k* ≥ 4) fully-interacting complexes. These maximal many-body complexes are grouped together by *principal loop*, namely, the longest genomic span in base pairs within each *k*-body interaction.

**Figure 7:**
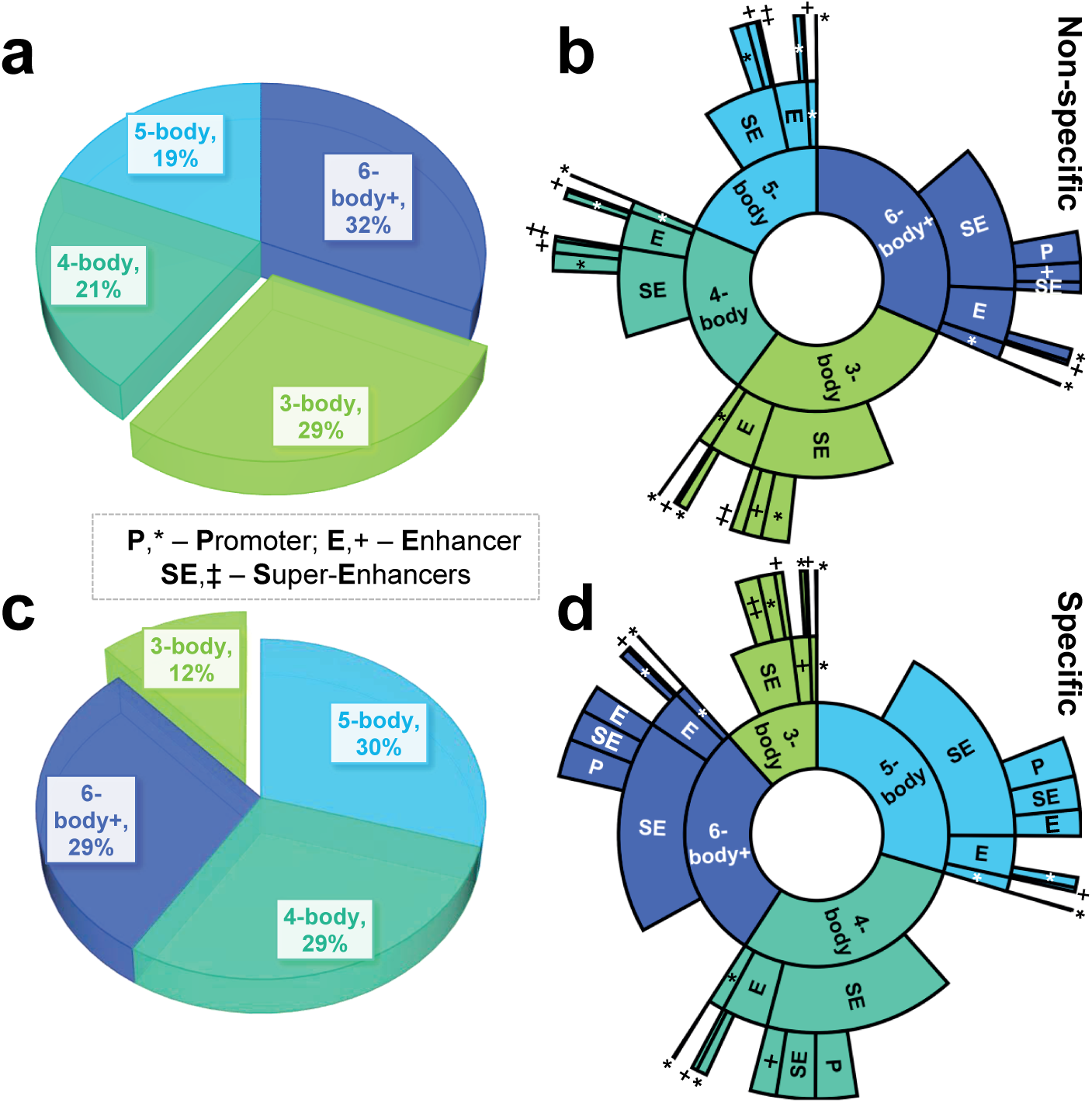
Functional landscape of principal loops in many-body chromatin interactions. A *principal loop* is the longest loop (in bp) among chromatin regions forming a many-body (≥ 3) interaction, where all pairs of bodies (i.e. chromatin regions) forming the interaction are within ≤ 80 nm Euclidean distance (46). The pie (**a,c**) and innermost ring of the sunburst (**b,d**) plots both show the proportion of specific (*bottom*) and non-specific (*top*) principal loops within maximal 3-, 4-, 5-, or ≥6-body interactions; the 2 outer rings(**b,d**) show the corresponding fraction of principal loops with functional annotations – super-enhancer (SE), enhancer (E), promoter (P) – where gaps represent the fractions of principal loop regions with no known SE, E, or P annotation. Only *maximal* many-body interactions are represented, *i*.*e*., no other chromatin region exits within the interaction distance such that all pairs are within 80 nm. Plots shown are the averages across all 39 modeled genomic loci.

Overall, we observe an increased proportion of specific maximal 4- and 5-body complexes relative to their non-specific counterparts (29 + 30 = 59 ± 0.9% SEM *vs*. 21 + 19 = 40 ± 0.5% SEM respectively, Figure 7a and 7c). Correspondingly, we observe a markedly decreased proportion of specific maximal 3-body complexes relative to non-specific maximal 3-body complexes (12±1% SEM and 29 ± 1% SEM respectively, Figure 7a and 7c). That is, maximal higher-order interactions beyond 3-body are preferred in the SE-associated loci.

Furthermore, we observe a higher proportion of specific principal loops bridging ≥ 2 super-enhancers when compared to non-specific complexes, at 7.6±1.4% SEM *vs*. 1.9±0.5 SEM respectively (Figure 7b and 7d), with a significant *p*-value of 6.1 × 10^−7^ (Mann-Whitney U test on the corresponding medians of 4.1% and 0.7% respectively, Figure S7a). In addition, we observe a higher proportion of specific principal loops bridging super-enhancers to promoters when compared to principal loops of non-specific complexes, at 8.2 ± 0.9% SEM *vs*. 5.6 ± 0.7% SEM respectively (Figure 7b and 7d), with a *p*-value of 0.026 (Mann-Whitney U test on the corresponding medians of 7.0% and 4.6% respectively, Figure S7b). Taken as a whole, these findings suggest that specific principal loops within higher order complexes serve the important role of bridging functional genomic regions to allow spatial coupling.

### Open and transcriptionally active chromatin is predictive of regions enriched in principal loops of many-body interactions

We then asked whether biological markers along the linear genome, such as epigenetic modifications, contained information on the specific higher-order physical interactions uncovered through our extensive 3-D modeling. While these loci with super-enhancers are enriched in active markers such as H3K27ac, we want to know if there are markers within the context of the enriched background that can differentiate regions of specific from non-specific many-body interactions. Notably, we asked whether biological markers could predict regions enriched in anchors of specific many-body principal loops.

To this end, we tested whether 5 KB intervals enriched in specific principal loop participation could be predicted using publicly available data, *e*.*g*., the ENCODE reference epigenome for GM12878 cells (ENCSR447YYN, Table S2) (52; 53). For this task, we built a machine learning classifier based on random forest (Figure 8, Methods) (54; 55).

**Figure 8:**
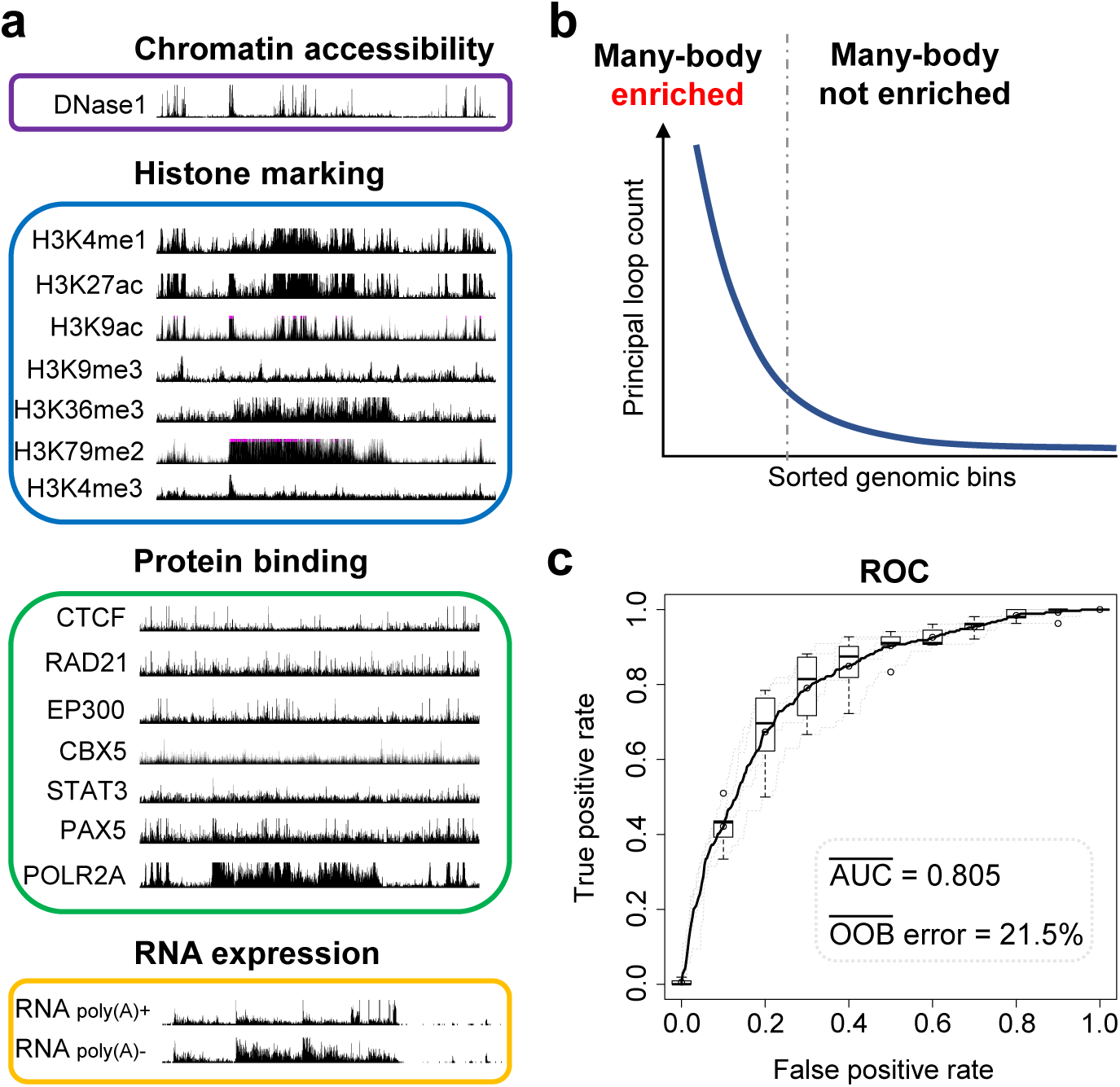
Predictive model for principal loop enrichment. **a** Publicly available biological datasets (Table S2), primarily from ENCODE reference epigenome for GM12878 (ENCSR447YYN) (52; 53), were used as predictive inputs to a random forest (54; 55) machine learning classifier. Illustrative signals shown are from the UCSC genome browser (74; 75) for locus chr 12: 11,690,000 – 12,210,000. **b** Cartoon illustration of *enriched* versus *not enriched* regions. Genomic regions, each corresponding to a non-overlapping 5 KB bin, were sorted based on principal loop participation; a subset of those occurring above the *elbow* inflection point were labeled as *enriched*; those occurring below the inflection point were labeled as *not enriched* (see Methods). **c** Receiver operating characteristic (ROC) curve (76) showing performance of our random forest classifier in discriminating principal loop *enriched* from *not enriched* genomic regions. Trained random forest model showed a mean area under the curve (AUC) of 0.805 on test set and a mean out-of-bag (OOB) error, an unbiased estimate of generalization error (54), of 21.5% over 5-fold cross-validation.

Our predictor achieved good performance, with a mean ROC AUC of 0.804 and an out-of-bag error of 21.5% over 5-fold cross-validation (Figure 8c). Our results indicate that genomic intervals enriched with specific principal loop anchors can be identified by biological markers.

Inspection of our model revealed biological markers most predictive of principal loop enrichment are consistent with *open chromatin* and *active transcription* – *i*.*e*., increased signal intensities for DNase accessibility, POLR2A binding, H3K4me1, and nuclear fraction RNA (Figure 9). Box plots of the corresponding *z*-score signal distributions revealed significant differences among principal loop *enriched* versus *non-enriched* regions (Figure 9b and 9c). The active chromatin marker H3K27ac was also significantly increased in principal loop enriched regions (*p*-value = 4.0 × 10^−23^); however, likely due to close correlations with both DNase accessibility and H3K4me1 (Pearson coefficients of 0.81 and 0.68 respectively), H3K27ac itself was not considered as informative according to the feature importance criteria of our classifier (Figure 9c).

**Figure 9:**
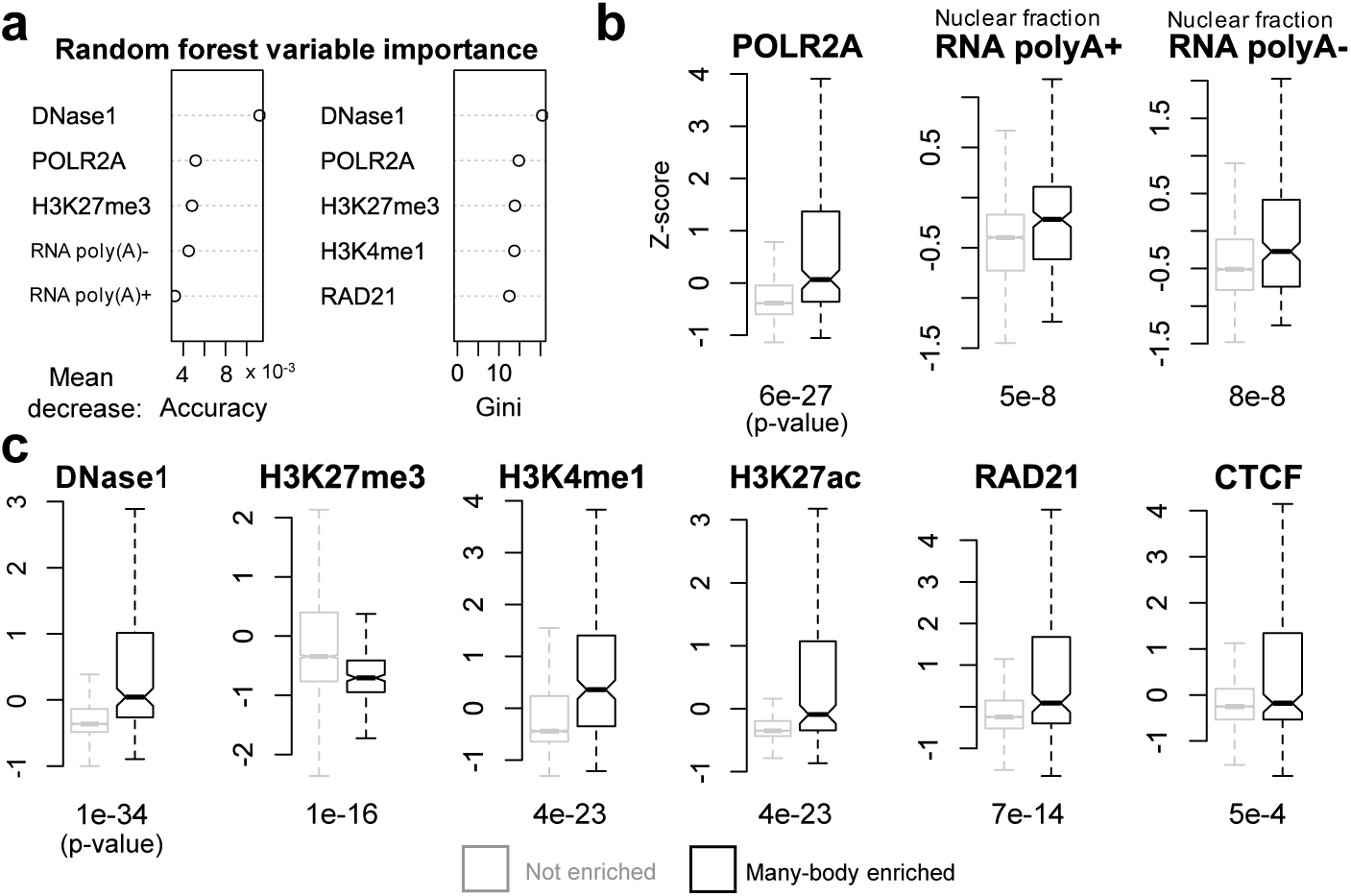
Predictive biological markers for principal loop enrichment. **a** Top 5 most important random forest predictors (*i*.*e*., variables or *features*) according to mean decrease in: accuracy (left) and gini coefficient (right) (54; 55). **b,c** Box plots of *z*-score distributions of predictive biological markers for principal loop *enriched* (black) and *not enriched* (grey) regions. *P*-values, according to Mann Whitney U testing for median difference among *enriched* versus *not enriched* regions, are listed below each box plot.

We also found that chromatin architectural protein CTCF and cohesin-subunit RAD21 exhibited significantly increased ChIP-seq signal intensities in principal loop enriched regions (*p*-value = 5.0 × 10^−4^ and 7.0 × 10^−14^ respectively); although RAD21 was found to be a more important predictor (Figure 9a and 9c).

Consistent with increased *active* markers, we found decreased ChIP-seq signal intensities for the *repressive* mark H3K27me3 to be predictive of principal loop enrichment (Figure 9a and 9c). Overall, we found that open and active chromatin markers, along with decreased repressive markers, to be strongly predictive of 5 KB intervals enriched for anchors of specific principal loops.

## Discussion

We have developed a computational model for identifying specific chromatin many-body interactions and for reconstructing their functional landscapes from population Hi-C contact frequencies. Our method exploits extensive biophysical folding simulations to infer dependencies among chromatin contacts. By incorporating the inferred dependencies into a Bayesian generative model (39), our method deconvolves the intrinsic single-cell chromatin contact states underlying the pairwise, population-averaged Hi-C data.

Our 3-D chromatin ensembles are highly realistic as they exhibit spatial interaction frequencies across many loci at Pearson correlations of 96–97% to the measured Hi-C. This close level of correlation is significant, as only basic biophysical assumptions are made (*e*.*g*., an 80 nm interaction distance threshold and nuclear volume confinement) with no adjustable parameters. This is in contrast to several prior studies where each domain or bead modeled requires a separate adjustable parameter (56; 57).

Furthermore, the reconstructed 3-D chromatin ensembles are generated from a very sparse set of interactions – just ∼5% of the predicted specific Hi-C interactions are sufficient to produce polymer ensembles with contact frequencies consistent with Hi-C measurements (Figure 3). Notably, our models indicate that only 15–32 interactions are sufficient to reconstruct loci of size 480 KB to 1.94 MB. Hence, these sparsely selected sets are likely enriched with interactions driving the chromatin fold (22; 46).

Our computed 3-D chromatin ensembles contain rich structural information, allowing prediction of *specific, i*.*e*., highly *non-random*, many-body (≥ 3) chromatin interactions. Our predictions are overall concordant with SPRITE, with a majority of modeled genomic loci exhibiting significantly elevated median coverages for specific *vs*. non-specific many-body interactions.

The landscape of many-body interactions emerging from our analysis of 39 active genomic loci showed super-enhancers (SE) as enriched in specific many-body principal loop participation compared to non-SE regions (*p*=2.24×10^−129^, Figure S8), with overall levels of SE-SE and SE-promoter interactions elevated in specific many-bodies (Figures 6 and 7). While the loci studied were *a priori* selected based on SPRITE clusters containing multiple super-enhancers, SPRITE measurements *per se* cannot distinguish direct from indirect cross-linking. Therefore, to our knowledge, this work is the first to provide computational evidence, with measurable Euclidean distances estimated from our modes, that super-enhancers are *directly* and *non-randomly* interacting spatially with other functional genomic regions in many-body complexes (18). These predictions can be tested experimentally.

Our principal loop heatmaps can reveal important insight into the higher-order spatial organization of chromatin. As an example, Figure 10 shows that at the SH3KBP1 locus, regions participating in many-body principal loops generally do not appear to be forming domains, with the exception of 3-body principal loops which appear to resemble the patterns of the original pairwise Hi-C (Figure 3d). Instead, as evidenced by the banding patterns of the 4-, 5-, and 6-body heatmaps (bottom row of Figure 10), principal loops may primarily be facilitating direct, long-range interactions among functional genomic regions such as super-enhancers, enhancers, and promoters. Such banding patterns at 5 KB are likely not due to A/B compartmentalization (100 KB - 1 MB scale), as our loci are mostly (> 90%, Table S1) in A compartments. This is consistent with our functional landscapes exhibiting decreased preference for maximal 3-body complexes and relatively increased functional associations among specific many-bodies (Figures 6 and 7).

**Figure 10:**
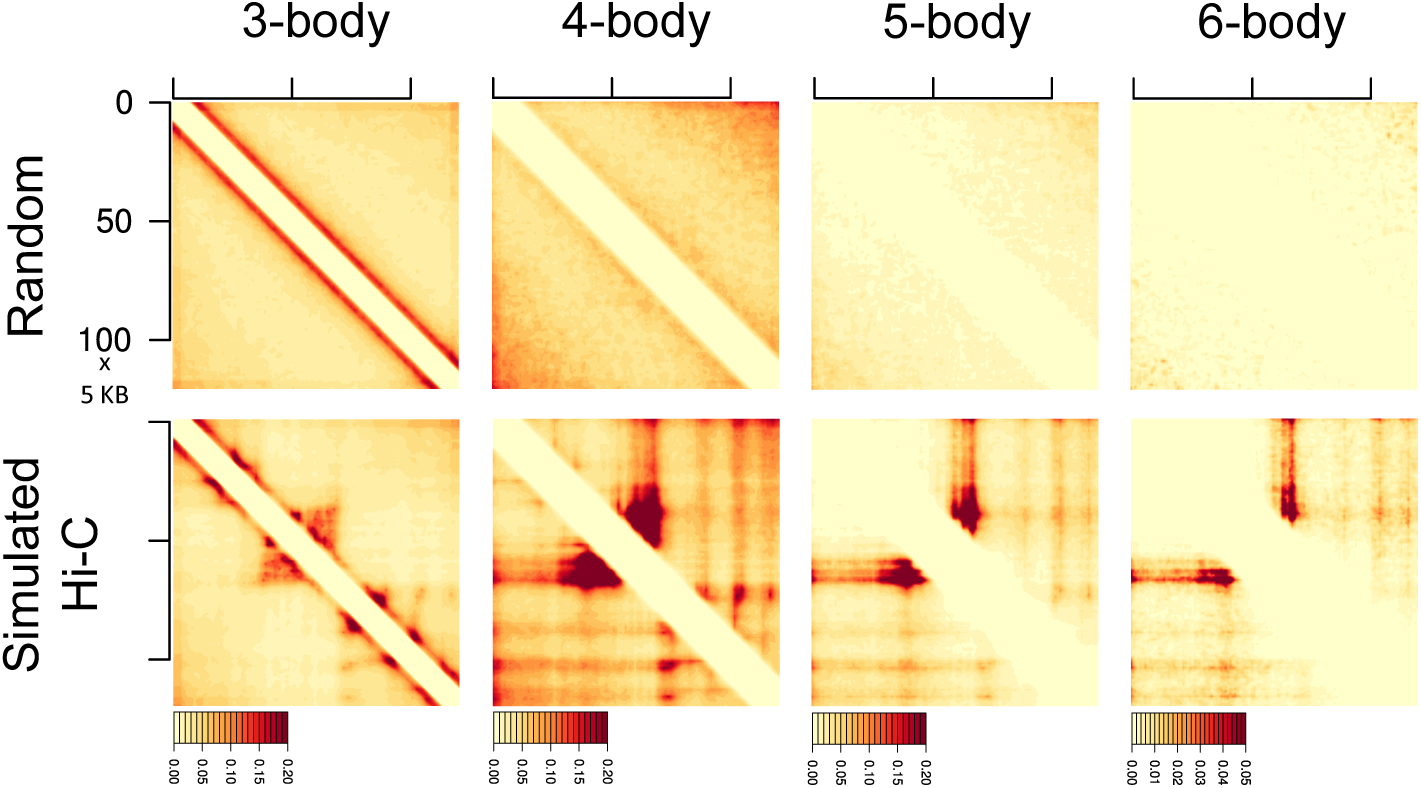
Principal loop heatmaps. Heatmaps are for the TAD (arrowhead) region containing the SH3KBP1 genomic locus (chr X: 19,560,000 – 20,170,000). For reference, the corresponding measured Hi-C is shown in Figure 3d. Columns, from left to right, are for principal loops within 3-, 4-, 5-, and 6-body chromatin interactions respectively. The rows show the principal loop interaction frequencies captured under random (*top*) and deconvolved, single-cell (*bottom*) folding after aggregation. Axes of all heatmaps are in units of 5 KB.

In contrast to other models which focus on heterochromatin condensation (29), we instead examine highly active chromatin regions. Our analysis showed that even in super-enhancer loci where active markers are enriched at baseline, open chromatin (DNase hypersensitivty) and the presence of active transcriptional marks such as POLR2A and nuclear fraction RNA are predictive of 5 KB regions enriched for anchors of specific many-body principal loops. Our findings are consistent with the opinion that nuclear RNAs may be important factors for nuclear organization through promotion of phase separation and ultimately enhancer-promoter looping (58; 59).

## Conclusions

We have developed CHROMATIX, a computational framework for predicting the intrinsic 3-D structural ensembles underlying population-averaged Hi-C data; our method is general and can be applied to other cell lines where pairwise chromatin contact information is available. We demonstrate our predicted 3-D structural ensembles have close correlation with the measured Hi-C data over 39 modeled genomic loci. Our CHROMATIX framework can also identify specific many-body chromatin interactions, and we show the predicted many-body interactions to be broadly concordant with SPRITE clusters.

We find our predicted specific many-body interactions to be significantly associated with functional genomic regions such as SEs and promoters; further, they preferentially form maximal 4- or higher-order interactions over 3-body interactions. These findings are consistent with specific principal loops likely playing the important role of bridging many genomically distant regions and allowing them to condense into functional assemblies through direct spatial contact. Overall, the many-body interactions uncovered in this study may serve as the 3-D manifestations of phase-separated, multi-valent assemblies among super-enhancer regions (10).

Further, we have shown that genomic regions enriched in anchors of principal loops are also enriched in open and active chromatin marks such as DNase accessibility, POLR2A, H3K4me1, H3K27ac, and nuclear fraction RNA, and depleted in the repressive mark H3K27me3. These biological markers are likely representative of factors needed to condense distant chromatin regions into ordered, spatial complexes necessary to regulate fundamental cellular processes such as gene transcription.

The CHROMATIX method has the promise of generating high-resolution 3-D ensembles of chromatin structures with detailed information of spatial many-body interactions using abundantly available population-averaged Hi-C data. As only about 5% of specific interactions are sufficient to reproduce measured Hi-C frequencies, CHROMATIX can provide higher resolution details beyond that of input Hi-C measurement.

Our method enables quantification of the extent of specific 3-, 4-, and higher-order many-body interactions at a large scale. It also elucidates the functional implications by providing details on how super-enhancers, enhancers, promoters, and other functional units probabilistically assemble into a spatial apparatus with measurable Euclidean distances. Our method can predict specific many-body interactions solely from markers along the linear genome and allows insight into the biological factors that drive the spatial coordination among genomic regions. Finally, our method can simulate multiple independent loci located on separate chromosomes within the same confining nuclear volume, and can be applied to identify specific inter-chromosomal many-body interactions.

## Methods

We now provide technical details on key components of the CHROMATIX method (Figure 2).

### Calculating *p*-values for calling specific Hi-C interactions

To assign statistical significance *p*-values to each Hi-C measured interaction, we use a scalable *Bag of Little Bootstraps* resampling procedure (60) over the uniform random 3-D polymer ensemble, with 10,000 outer replicates, to obtain a null distribution over random chromatin contacts. P-values are assigned to each Hi-C contact frequency based on the proportion of bootstrap replicate contact frequencies exceeding the measured Hi-C at the same genomic distance.

### Polymer simulation of structural perturbations

To predict which specific contacts are likely co-occurring within individual cells of the population, we carried out extensive structural *perturbation* simulations. These biophysical simulations were used to elucidate dependencies and infeasible geometries among chromatin contacts. We incorporated information from the perturbed simulations into a sparsity-inducing Bayesian prior distribution over hypothetical folding mechanisms among the specific contacts, where each mechanism is encoded in the form of a directed acyclic graph (DAG) (61; 62). A considered DAG, in which each edge represents a possible causal dependency between two contacts, is restricted according to computational *knock-in* perturbations supporting such a hypothesis; specifically, if knocking-in a contact is observed to significantly upregulate the frequency of another contact beyond random, a directed edge from the knocked-in contact to the upregulated contact is then available to be sampled when generating folding mechanisms. Given the observed population Hi-C data and the results of simulated biophysical perturbations, we infer the posterior distribution of single cell contact states through Gibbs sampling (see Supplementary Information for details on sampling procedures). We find that our models for 38 out of the 39 loci have higher posterior probabilities than the naive models of product of independent pairwise contacts. The naive models further suffer from the inability to recognize geometrically infeasible combinations of pairwise contacts.

### Functional annotation and loci selection

We used LILY (63) to detect functional genomic regions containing super-enhancers, enhancers, and promoters based on H3K27ac ChIP-seq data of GM12878 cells (64)(see Table S3). We used publicly available SPRITE data for GM12878 cells (18) to select clusters containing multiple (≥ 2) super-enhancers as a basis for investigating if many-body interactions may form among multiple super-enhancers. We then used publicly available Hi-C data for GM12878 at 5 KB resolution (9) to identify the median TAD (≤ 2 MB, arrowhead domain) boundaries for the considered SPRITE clusters. After discarding regions with greater than ∼25% overlap, we obtained 39 genomic loci (Table S1), 35 of which have no overlap, for further investigation of many-body interactions. Hi-C contact counts at each locus, normalized via Knight-Ruiz matrix balancing (65), were obtained using Juicer (66) also at 5 KB resolution.

### Cliques and maximal many-body interactions

We extend the nCSAC approach of Gürsoy *et al*. (21; 22) to identify *specific* many-body (≥ 3) chromatin interactions. We define a *many-body* interaction as a complex of 5 KB chromatin regions such that the Euclidean distances between all pairs of regions in the complex are within a cross-linking threshold of ≤ 80 nm (46). Using graph theory terminology, a many-body interaction is equivalent to a *clique* (67), *i*.*e*., a fully connected graph such that all pairs of vertices are connected by undirected edges. Further, a many-body complex, or clique, is *maximal* if no additional chromatin regions may be added such that all pairs remain within the cross-linking threshold. We use the highly optimized graph analysis library *igraph* to detect many-body interactions within a 3-D polymer (68).

### Calling specific many-body interactions

To generate a *null* distribution over many-body chromatin interactions, we first tally the frequency of each observed many-body interaction within a uniform randomly folded ensemble of 75,000 polymers. We repeat the tally procedure by bootstrap resampling over the full polymer ensemble for 1,000 total replicates; this produces a distribution over the many-body interaction frequencies under a *null* hypothesis of random folding. For 3-body interactions (Figure 6), we detect all cliques consisting of *exactly* 3 distinct chromatin regions and do not require them to be maximal; that is, these 3-bodies may be part of a larger fully-connected complex. For principal loop analysis, we detect cliques consisting of *at least* 3 distinct chromatin regions and require that each clique is maximal (Figure 7).

We then identify *specific* many-body interactions at a locus by first tallying the corresponding many-body frequencies within each sample of the CHROMATIX deconvolved Hi-C ensemble (*i*.*e*., *simulated Hi-C*) of 25,000 polymers. We stratify the many-body frequencies (random and simulated Hi-C) according to both genomic distance and clique size. Specifically, for 3-body interactions shown in Figure 6, we stratify all frequencies based on principal (*i*.*e*., longest) and minor (*i*.*e*., shortest) loop spans in base pairs. For maximal principal loop interactions shown in Figure 7, we stratify based on clique size and the base pair span of the principal loop. Stratification is necessary to control for genomic distance bias, *i*.*e*., the fact that genomic regions with short genomic separation tend to spatially co-locate (21), and that larger clique sizes tend to allow correspondingly longer genomic distances to interact spatially with increased frequency. We assign a *p*-value to each simulated Hi-C many-body frequency as the within-stratum proportion of random (bootstrap-replicated) many-body frequencies that exceed the simulated Hi-C many-body frequency. Finally, to control for multiple testing, a simulated Hi-C many-body interaction is called *specific* if the FDR-adjusted (69) *p*-value is < 0.05.

### Concordance with SPRITE

We compared our *3-body* and *maximal many-body principal loop* predictions with publicly available SPRITE data for GM12878 (18). To adjust for genomic distance bias, we stratified principal loops according to base pair span and computed the SPRITE *coverage fraction, i*.*e*., proportion of SPRITE clusters that contained each principal loop complex. Specifically, we computed the median SPRITE coverage fraction at each 5 KB genomic distance span for both *specific* and *non-specific* principal loops (Figure S5). At each of the 39 modeled loci, we assessed the significance of the proportion of specific medians exceeding the corresponding non-specific medians by permutation testing: we randomly permuted the *specific* and *non-specific* labels assigned to each principal loop and re-computed the proportion of specific medians exceeding non-specific medians for 1,000 total replicates. We then assigned a *p*-value to each locus by the fraction of permutation replicates exceeding the observed proportion. A similar procedure was performed for 3-body predictions, with stratification by both principal and minor loop. To control for multiple testing, *p*-values where called significant if < 0.05 after FDR-correction (69).

### Predictive model for principal loop enrichment

We built a random forest machine learning classifier (54) to identify biological markers predictive of regions enriched in the principal loop anchors of many-body complexes. We used publicly available biological datasets (Table S2), primarily from ENCODE reference epigenome for GM12878 (ENCSR447YYN) (52; 53), as our input features (Figure 8a). At each of the 39 modeled loci, genomic regions corresponding to non-overlapping 5 KB bins were sorted based on principal loop participation; a subset of those occurring above the “elbow” inflection point (Figure 8b) were labeled as *enriched*; those occurring below the inflection point were labeled as *not enriched*. To avoid ambiguous labels and to provide a more robust decision boundary among *enriched* versus *not enriched* regions, we retained the top 20% of the above-elbow fraction at each locus and discarded the remainder, while still retaining all samples below the elbow. Our final data set consisted of 231 regions enriched (*i*.*e. positive*) in many-body interactions and 5,800 regions not-enriched (*i*.*e. negative*). To control for potential class imbalance issues during training, we used the *randomForest* R package (55) with stratified resampling to present equal number of positive and negative samples to each decision tree (*n* = 500) in the random forest. Classifier performance results, mean ROC AUC of 0.805 and out-of-bag error of 21.5% (Figure 8c), were obtained on a held out test set (∼20% of labeled samples) over 5-fold cross-validation using the *caret* R package (70).

## Supporting information

Table of genomic regions modeled for this study.

Table of public biological datasets used.

Table of LILY annotations for super-enhancer, enhancer, and promoter regions.

## Availability of data and materials

All C++ source code for chromatin polymer folding as well as comprehensive tutorials demonstrating how to configure polymer folding simulations are publicly available via git repository (71). Similarly, C++ source code for our Bayesian Hi-C deconvolution Gibbs sampler and loci modeling scripts are also available via git (72; 73).

## Ethics approval and consent to participate

Not applicable.

## Author’s contributions

APR and JL conceived and designed the study. APR designed and implemented the algorithms. APR carried out computation and analysis, with assistance from QS. BH assisted in computing. VB provided tool and analysis of genome annotations. ZS provided additional analysis and contributed resources. JL supervised the overall study. APR and JL wrote the paper with input from QS, VB, and ZS, and assistance from BH. All authors read and approved the final manuscript.

## Acknowledgements

We thank Daniel M. Czajkowsky and Gamze Gürsoy for helpful discussions, Lin Du for help with A/B compartment calling, and Kimberly Beiting for her editing. This work is supported by grants NIH R21AI126308, R21AI137865, R01CA204962, and R35GM127084 (to JL); CAS-18H100000104 and NSFC-81627801 (to ZFS).

