## Supplementary methods and figures. for "CHROMATIX: computing the functional landscape of many-body chromatin interactions in transcriptionally active loci from deconvolved single-cells"

**2 Genomic Loci** **18**

**3 CHROMATIX 3-D Chromatin Reconstruction** **18**

**4 Comparison with single-cell studies** **20**

**5 Comparison with SPRITE data** **20**

**6 Supplementary Figures** **23**

### 1 Methods

#### 1.1 Background

We model chromatin polymers as self-avoiding walks (SAW) confined within the volume of the cell nucleus. Our goal is to sample chromatin polymers from a specified target distribution [9, 10, 11]. We use the sequential importance sampling (SIS) technique to achieve deep sampling of chromatin polymers from the desired target distribution of interest [16, 17]. Basics of SIS such as biased estimation, resampling, and rejection control can be found in [16, 17]. More advanced techniques developed for polymer modeling can be found in [9, 10, 11, 15].

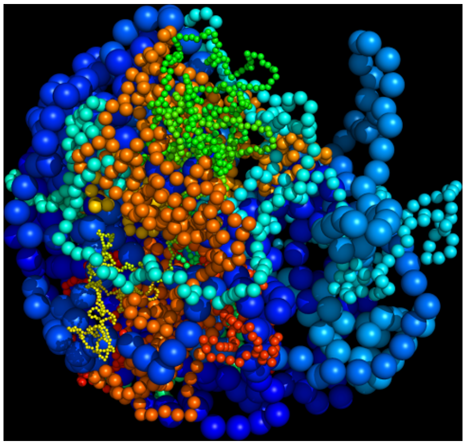

Figure 1: **3–D self-avoiding chromatin polymers generated by sequential importance sampling.** Our modeler can simulate multiple loci with heterogeneous chromatin fiber density within a confining nucleus.

##### 1.1.1 Null model

Our null model is that of the uniform distribution of all geometrically realizable SAW chromatin chains under a uniform energy model: all connected, self–avoiding, nuclear-confined chromatin folds have negligible difference in energy. That is, we assume a uniform *target distribution*:

$$P_t = \frac{1}{Z} \tag{1}$$

where $Z$ is the total number of connected folds of length $n$ that are self–avoiding and are within the bounding sphere. To sample from this target distribution, we use the SIS procedure with the intermediate *sampling distributions*:

$$P_s(X_m|x_1 \ldots x_{m-1}) = \frac{1}{q(x_1 \ldots x_{m-1})} \tag{2}$$

where $q(x_1 \ldots x_{m-1})$ is the number of available locations, given state $x_1 \ldots x_{m-1}$, such that the walk remains connected, self–avoiding, and within the confining nuclear sphere [9, 16]. Here, variables $x_1 \ldots x_{m-1}$ represent the 3-D spatial positions for the monomer indices $1 \ldots m-1$ along the polymer chain.

Since $Z$ is an unknown constant, we use the biased expectation estimator of Eqn (3):

$$\hat{Y}_{P_t} = \frac{\sum_{i=1}^{k} \dot{w}^{(i)} f(x_1^{(i)} \ldots x_n^{(i)})}{\sum_{i=1}^{k} \dot{w}^{(i)}}, \tag{3}$$

Where $f(\cdot)$ is the desired quantity (e.g. chromatin contact frequencies) to be estimated under the target distribution $P_t$. With uniform $P_t$, then $\dot{w}_m^{(i)} = \dot{w}_m^{(i-1)} q(x_1 \ldots x_{m-1})$ and $\dot{w}_m^{(0)} = 1$.

This null model of random physical chromatin chains controls effectively for genomic-distance dependent random interactions, regardless of the A/B compartmentalization of the loci.

The confinement of the null model can be either spherical or ellipsoidal, both supported by the CHROMATIX implementation. An example of a chromatin polymer folded under ellipsoidal confinement using our tool is shown below:

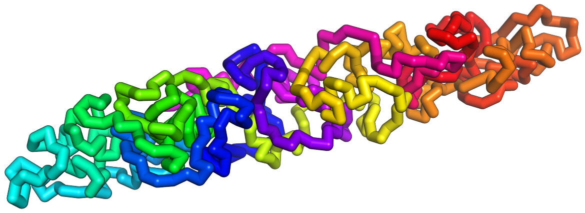

Figure 2: **An example of a chromatin chain in ellipsoidal confinement.**

#### 1.2 Fractal Monte Carlo

We describe the Fractal Monte Carlo (FMC) method used extensively in this study. FMC is a generalization of SIS, which enables multiple variance reduction techniques, including resampling, rejection control, and partial rejection control [16], to be unified under a single coherent framework. In brief, fractal Monte Carlo uses a recursive simulation strategy, where simulations at the same recursion depth must have homogeneous SIS configurations, but simulations may be heterogeneous from each other at different recursion depths. In parent simulations, polymer chains are grown to a desired *landmark* length by deferring to their child simulations for intermediate chain growth. The parents then select a single sample from each child ensemble upon completion of the child simulation, while correcting for sampling bias. Specifically, the weight of the selected child sample is divided by the child's selection probability, see 3. This process repeats until the target polymer size is reached. There is

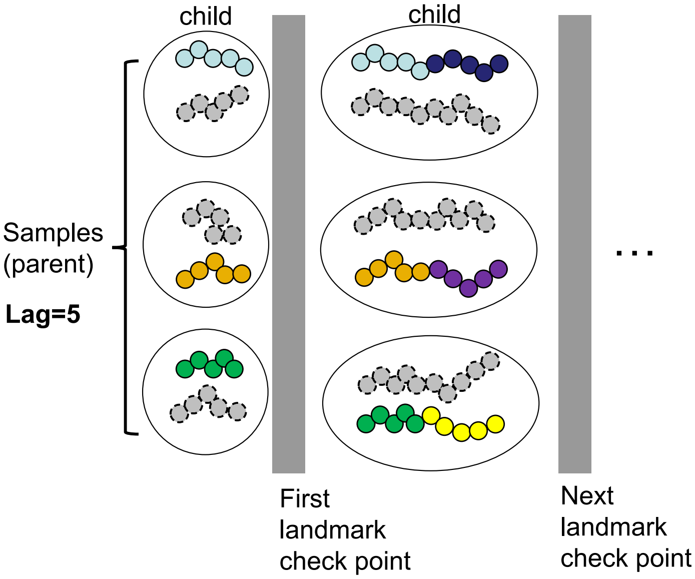

Figure 3: **An example of a 2-D fractal Monte Carlo simulation, with landmark checkpoints occurring after every 5th monomer bead is placed.** At each landmark checkpoint, which occurs after every 5th bead is placed, parents select from their respective child ensembles according to a selection probability, which is typically a monotonic function of the child's relative weight – for instance, the canonical resampling selection probability: $P_{rs}(\vec{x}_m^{(i)}) = \frac{w_m^{(i)q}}{\sum_{j=1}^{k} w_m^{(j)}}$. Here *lag* refers to a constant checkpoint interval, in this case the lag is 5. In this 2-D simulation, the child simulations are represented by the ensemble-containing circles and ellipses, the parent simulation is implicitly represent as containing all child simulations. For each child ensemble, the culled fragments which fail to pass landmark checkpoints are represented by grey dotted chains, whereas the retained fragments (i.e. those selected by the parent simulation) are represented by the colored (e.g. orange, blue, yellow, purple, etc) chains.

no conceptual limit on the recursive depth of the simulations; for instance the aforementioned child simulations may in turn defer to their own sub-simulations when generating their ensemble.

We refer to the *dimension* of an FMC simulation based on the recursion depth. For instance, a simulation consisting of a parent and a child, as in 3, is called a *2–D* fractal simulation. If the child simulations also deferred to their own sub-simulations, this then constitutes a *3–D* fractal simulation. Under the FMC framework, canonical resampling and rejection control SIS procedures can be considered as *1-D* fractal simulations.

FMC can help alleviate problems with canonical variance reduction techniques. For instance, resampling is generally associated with a reduction in sample diversity. A two-dimensional fractal simulation can address this issue by allowing the child simulation to use resampling, while the parent simulation only collects a single enriched sample from the child ensemble. This procedure would generate an ensemble with a degree of weight enrichment but retention of sample diversity.

Another usage is improved rejection control, which is known to have severe issues with scalability to higher dimensions. Since fractal Monte Carlo grows samples in stages, rejected samples within a child simulation are only regrown from the previous parent stage. We have observed dramatic performance increases when using rejection control in this fashion. Note, to allow pooling of importance weights from fractally grown samples with rejection control, we compute sample weights according to:

$$w_m^{(i)*} = \frac{Z_c \cdot w_m^{(i)}}{P_{rc}(\vec{x}_m^{(i)})} \tag{4}$$

where $P_{rc}$ is the canonical rejection control probability as defined in [18] and can be inferred from Eqn 5, and the term $Z_c$ is the partition constant:

$$Z_c = \int P_{rc} P_s(X_m = x_m) dx_m = \int \min\{1, \frac{w_m^{(i)}}{c}\} P_s(X_m = x_m) dx_m \tag{5}$$

An unbiased estimator for $Z_c$ is given by the following expression [16]:

$$Z_c = E_{P_s}[\min\{1, \frac{w_m^{(i)}}{c}\}] \approx \frac{1}{k} \sum_{j=1}^{k} \min\{1, \frac{w_m^{(i)}}{c}\}, \tag{6}$$

where $k$ is the ensemble size.

##### 1.2.1 Comparison of SIS weight enrichment methods

We compared a 3-D fractal Monte Carlo simulation against the following alternatives: no weight enrichment, canonical resampling, and rejection control. Specifically, the FMC simulation consisted of a top-level grandparent which selected its samples from a parent simulation with resampling; the parent simulation in turn selected its samples from a child simulation with rejection control. To allow comparison with canonical (1-D) rejection control, all simulations were limited to a polymer length of 150 monomer beads and an ensemble size of 50 polymer chains.

We compared the weight enrichment schemes using an $H_1$ score defined as the harmonic mean of the effective sample size (ESS) and the average proportion of unique (APU) beads across each monomer index within the resulting ensembles:

$$H_1 = \frac{2 \cdot ESS \cdot APU}{ESS + APU} \tag{7}$$

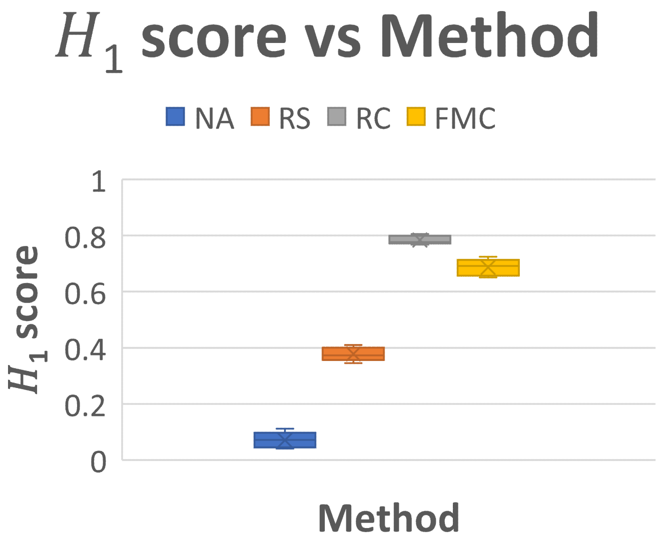

Figure 4: **Box plots of $H_1$ score distributions for several weight enrichment methods.** *NA* - no enrichment, *RS* - resampling, *RC* - rejection control, *FMC* - fractal Monte Carlo

The ESS is defined as [16]:

$$ESS = (1 + \frac{\sigma^2(w)}{\mu^2(w)})^{-1}, \tag{8}$$

where $\mu(w)$ and $\sigma^2(w)$ are the mean and variance of the ensemble's importance weights, respectively. ESS is essentially a *signal-to-noise ratio* of the importance weights, and is used as a *rule of thumb* measure for how efficiently the sampling distribution is capturing the target distribution [16].

The APU is computed by first determining the ensemble's proportion of unique beads at each monomer index, then averaging this proportion across the length of the polymer chain. It describes the diversity of the resulting polymer ensemble.

In Fig 4, we see, as expected, *no weight enrichment* has the worst $H_1$ score due to a low mean ESS of 0.04. *Resampling* has the second worst $H_1$ score because, though it has the highest mean ESS of 0.65, this is largely due to low ensemble diversity reflected in the lowest mean APU of 0.27. *Rejection control* and *FMC* received the top two $H_1$ scores with mean ESS of 0.64 and 0.52 respectively. Note, all methods, with the exception of resampling, exhibited an APU of 1.0.

In Fig 5, we compare the CPU run times of the weight enrichment methods. Collectively with Fig 4, we see that FMC produces high quality samples with comparable $H_1$ scores to rejection control at a far lower CPU cost.

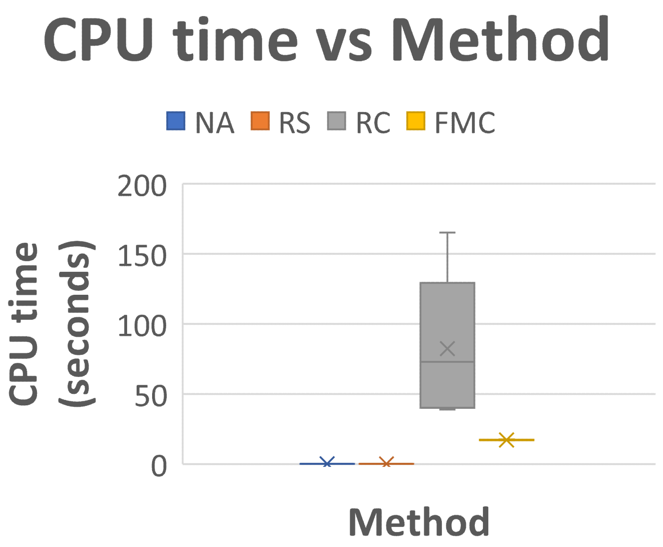

Figure 5: **Box plots of CPU times for several weight enrichment methods.** *NA* - no enrichment, *RS* - resampling, *RC* - rejection control, *FMC* - fractal Monte Carlo

A general issue for sequential importance sampling is that of outlier samples with exceedingly high weights. We find the outliers are usually more extended polymer chains. To make our random model more stringent, we remove these outliers using the interquartile rule (IQR, see [8]), where $IQR = Q_3 - Q_1$, $Q_1$ and $Q_3$ the 25-th and 75-th percentiles of the importance weights respectively. We retain samples with weights in the range of $Q_1 - 1.5 \cdot IQR$ and $Q_3 + 1.5 \cdot IQR$. This results in a more condensed random ensemble [16], hence calls of specific contacts against this condensed random ensemble are made more stringent; *i.e.*, specific many-body contacts identified will likely have more statistical significance than the $p$-value indicated by this random ensemble.

Overall, fractal Monte Carlo-based polymer folding provides a flexible framework with many configuration avenues for balancing CPU performance with sampling quality. The above example showed an FMC simulation with an $H_1$ score comparable to rejection control but with a CPU performance much closer to resampling.

##### 1.3 Hi-C deconvolution with CHROMATIX

*CHROMATIX* (CHROMATin mIXture) is a generative Bayesian decovolution method based on physical models of chromatin folding. It can provide estimates of contact probabilities among multiple ($\geq$3) interacting loci and can help to identify the cause-effect relationships among spatially interacting genomic regions. It can also help to identify driver interactions governing the folding state.

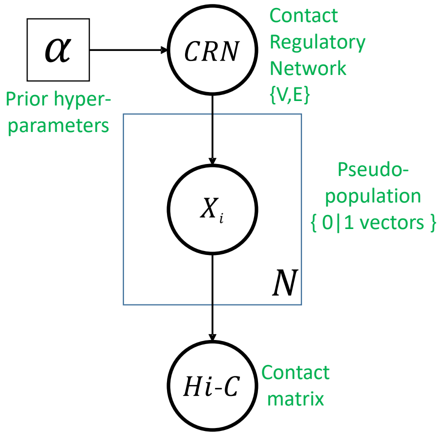

Figure 6: **Bayesian generative model for Hi–C.** Our model operates on a contact regulatory network ($CRN$), a latent population of single cell contact states (*i.e.*, pseudo-population), and the observed Hi–C contact matrix. The symbol $\alpha$ is used to represent specified model hyperparameters and prior geometric knowledge from 3-D biophysical folding simulations.

Fig 6 illustrates the graphical architecture of our Bayesian deconvolution model. It consists of the following random variables: a chromatin contact regulatory network ($CRN$), a latent population of single cell contact states ($\{X_i\}$), and the observed $Hi$-$C$ contact matrix. We provide a brief overview of each of these variables.

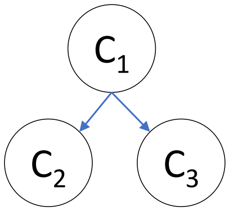

Figure 7: **Pedagogical contact regulatory network ($CRN$).**

**Contact regulatory network (CRN)** The $CRN$ represent a mechanistic folding *hypothesis* encoded as directed acyclic graph (DAG); this allows simple interpretation as well as efficient sampling of the latent population of contact states. Fig 7 demonstrates a small $CRN$ over three chromatin contacts $C_1$, $C_2$, and $C_3$. This instance of DAG may be interpreted as the following hypothesis "contact $C_1$ regulates contacts $C_2$ and $C_3$".

The symbol $\alpha$ in Fig 6 encodes the prior hyperparameters over the space of $CRN$s. Specifically, $\alpha$ encodes which directed edges are allowed to be represented, where an edge represents a folding dependency between the corresponding contacts. These edges are identified by simulated *perturbation* experiments, where a chromatin contact is knocked-in to assess its influence on the other modeled contacts. If knocking-in contact $C_i$ is observed to significantly upregulate the frequency of contact $C_j$ beyond random, then a directed edge from $C_i$ to $C_j$ is *allowed*, but not required, to be present in the $CRN$. Empirically, we have found contacts to have very localized effects, and therefore the perturbation edge restrictions greatly reduce the space of reachable $CRN$s.

An additional geometrical constraint is encoded in $\alpha$, which are the infeasible contact configurations. For example, the coexistence of contacts $C_i$, $C_j$, and $C_k$ may not be embeddable in 3-D Euclidean space. We run a heuristic search over the space of contact configurations to identify putative infeasible configurations; specifically, if our chromatin folder cannot satisfy the configuration proximity constraints in a threshold number of attempts (*e.g.* 50,000), then we label the configuration infeasible. $CRN$s are penalized based on the assigned probability mass to the infeasible corpus.

**Latent population of single-cell contact states.** The latent population $\{X_i\}$, called *pseudo-population* in Fig 6, represents the deconvolved single-cell contact states for each of the contacts modeled by the $CRN$. It is a matrix where each column $X_i$ is a (0, 1) vector, representing the knock-in, knock-out states respectively of each modeled contact within a single cell of the population.

**Simulated Hi–C contact matrix.** The simulated $Hi$–$C$ contact matrix contains the empirical frequencies of the modeled chromatin contacts. It can be interpreted as arising from noisy observation of the aggregated contact frequencies within the latent population $\{X_i\}$ of single-cell contact states.

##### 1.3.1 Gibbs sampling

Our generative Bayesian model of chromatin folding, illustrated in Fig 6, defines a probabilistic graphical model (PGM). It is a Bayesian network [14, 19, 20, 22] defining a joint probability distribution over the random variables $CRN$, $\{X_i\}$, and $Hi$-$C$:

$$P(CRN, \{X_i\}, Hi\text{-}C|\alpha) = P(CRN|\alpha) \cdot P(\{X_i\}|CRN) \cdot P(Hi\text{-}C|\{X_i\}) \quad (9)$$

Since $Hi$–$C$ data is observed, we can instead sample from the *posterior* distribution using Markov-chain Monte Carlo (MCMC):

$$\begin{aligned} &P(CRN,\, \{X_i\}|Hi\text{-}C = M_H,\, \alpha) \\ &\qquad \propto\, P(CRN|\alpha) \cdot P(\{X_i\}|CRN) \cdot P(Hi\text{-}C = M_H|\{X_i\}), \end{aligned} \quad (10)$$

where $M_H$ is the empirical Hi-C data.

MCMC, specifically Gibbs sampling [6, 7, 24], can be used to generate contact regulatory networks and latent populations of single-cell contact states (pseudo-populations) from the posterior distribution (Eqn 10). Gibbs sampling involves repeated draws from the family of conditional probability distributions $V_i' \sim P(V_i'|\{V\}_{-i})$, where $V_i$ is the random variable being updated conditional on the current states of all *other* $\{V\}_{-i}$ random variables in the system [6, 7, 24]. In the the case where direct sampling is not feasible, *e.g.*, due to non-conjugacy, we can instead perform *Metropolis-within–Gibbs*, *i.e.*, replace the Gibbs update with a single Metropolis–Hastings update [6].

We use a *reversible* Gibbs sampling approach which satisfies *detailed balance* [5]. It is a special case of Metropolis-Hastings sampling [6, 7]. To construct a reversible Gibbs sampler, we uniform randomly select a variable to update during each Markov step rather than deterministically iterate over the set of random variables as in canonical Gibbs sampling. Since this approach is equivalent to Metropolis-Hastings, namely, it produces a reversible Markov chain satisfying detailed balance, the *stationary distribution* of the Markov chain asymptotically converge to the target posterior distribution of Eqn 10 [5]. Furthermore, randomly sampling the variables rather than in a fixed order may help with stationary distribution convergence [21].

Ultimately, the Gibbs-sampled latent population of single-cell contact states (*i.e.*, pseudo-population) serves as input to the fractal Monte Carlo chromatin folder for generation of 3–D structures which are consistent with the measured *Hi-C* frequencies. We suggest retention of estimated *maximum a posteriori* (MAP) contact states for 3-D folding.

##### 1.3.2 Markov updating the CRN

We now describe the Markov updates for each of the latent random variables, specifically, the contact regulatory network $CRN$ and the latent pseudo-population $\{X_i\}$. The target Gibbs conditional distribution, $P_{tar}$, for the $CRN$ is given by Eqn (11). We use integration $\int P(\ldots, V)dV$ to universally denote marginalization of a random variable $V$ regardless of its domain. We have:

$$\begin{aligned} P_{tar}(CRN) &= P(CRN|\{X_i\}, Hi\text{-}C, \alpha) \\ &= \frac{P(CRN, \{X_i\}, Hi\text{-}C|\alpha)}{\int P(CRN, \{X_i\}, Hi\text{-}C|\alpha)dCRN} \\ &= \frac{P(CRN|\alpha)P(\{X_i\}|CRN)P(Hi\text{-}C|\{X_i\})}{\int P(CRN|\alpha)P(\{X_i\}|CRN)P(Hi\text{-}C|\{X_i\})dCRN} \\ &= \frac{P(CRN|\alpha)P(\{X_i\}|CRN)P(Hi\text{-}C|\{X_i\})}{P(Hi\text{-}C|\{X_i\})\int P(CRN|\alpha)P(\{X_i\}|CRN)dCRN} \\ &= \frac{P(CRN|\alpha)P(\{X_i\}|CRN)}{\int P(CRN|\alpha)P(\{X_i\}|CRN)dCRN} \\ &\propto P(CRN|\alpha)P(\{X_i\}|CRN) = Q_{tar}(CRN) \end{aligned} \tag{11}$$

We can perform a Metropolis-within-Gibbs update since the denominator

Eqn (12)

$$Z_{CRN} = \int P(CRN|\alpha)P(\{X_i\}|CRN)dCRN \tag{12}$$

is a normalizing constant and cancels out in the classic Metropolis-Hastings acceptance criterion given by Eqn (13):

$$\begin{aligned} P_{acc}(CRN') &= \min\{1, \frac{P_{tar}(CRN')P_{pro}(CRN|CRN')}{P_{tar}(CRN)P_{pro}(CRN'|CRN)}\} \\ &= \min\{1, \frac{Z_{CRN}^{-1}Q_{tar}(CRN')P_{pro}(CRN|CRN')}{Z_{CRN}^{-1}Q_{tar}(CRN)P_{pro}(CRN'|CRN)}\} \\ &= \min\{1, \frac{Q_{tar}(CRN')P_{pro}(CRN|CRN')}{Q_{tar}(CRN)P_{pro}(CRN'|CRN)}\} \end{aligned} \tag{13}$$

where $P_{pro}$ is the *CRN proposal* distribution to be discussed shortly; but first, we must define the *target* factor probabilities $P(\{X_i\}|CRN)$ and $P(CRN|\alpha)$ from Eqn (11).

**Factor distribution for the latent population of single-cell contact states.** We now define the factor distribution $P(\{X_i\}|CRN)$ for the latent population of single-cell contact states (*i.e.*, pseudo-population). Note that the $CRN$ is not just a DAG, but a probabilistic graphical model, specifically a Bayesian network. The nodes of the $CRN$ are conditional probability tables (CPTs) which define Bernoulli distributions conditioned on the parent contact states, where the parent nodes are according to the directed edge relations in the network. Fig 8 revisits the CRN from Fig 7, but with CPTs now at each of the nodes. The factor distribution $P(\{X_i\}|CRN)$ can be *forward sampled* [14] from the Bayesian network defined by the $CRN$. In forward sampling, the states of the parent nodes are sampled before the child nodes (according to any topological sort such that parents are listed prior to children); this is always possible since the network is a DAG.

Therefore, the pseudo-population factor distribution $P(\{X_i\}|CRN)$ is a product of binomial, specifically *Bernoulli*, distributions and defines the population $\{X_i\}$ such that each $X_i$ is set of binomial random variables that can be forward sampled according to any valid topological ordering given by the directed edges of the CRN.

**CRN factor distribution.** We now define the contact regulatory network (CRN) factor distribution $P(CRN|\alpha)$. A CRN Bayesian network consists of a set of edges $E$ and a set of conditional probability distributions $CPD$, where each element $c$ of $CPD$ defines a column within a node's conditional probability table according to the format specified by Fig 8. We define the target factor distribution as:

$$P(CRN|\alpha) \propto \kappa^{|E|} \cdot \exp\{-\beta \cdot \text{energy(infeas)}\} \cdot \prod_{\text{c}\in\text{CPD}} \text{Dir}(\theta) \tag{14}$$

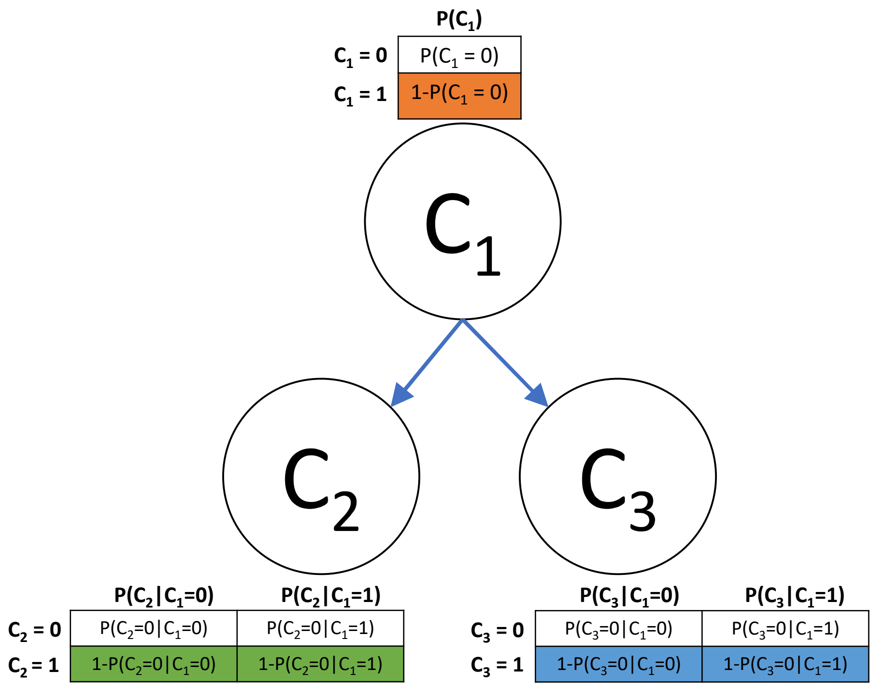

Figure 8: **Pedagogical contact regulatory network ($CRN$) as a Bayesian network.**

The expression $\kappa^{|E|}$, in which $\kappa$ is a hyperparameter $\kappa \in (0, 1]$ and $|E|$ is the cardinality of the edge set $E$, is an *edge-regularizing* prior introduced by (Heckerman et al, 1995) [12]. It induces a penalty on the number of edges and thereby promotes network sparsity, whereby the penalty increases as $\kappa$ approaches 0.

The expression $\exp\{-\beta \cdot \text{energy}(\text{infeas})\}$, where $\beta \in \mathbb{R}_{\geq 0}$ and energy(infeas) is a monotonic function of the probability mass assigned to the putative infeasible geometries captured from biophysical modeling. This penalizes CRN Bayesian networks based on how likely they are to generate a geometrically infeasible set of contacts.

Lastly, the expression $\prod_{c \in CPD} Dir(\theta)$ is a Dirichlet prior on $CPD$ values. Specifically, each conditional probability distribution has a $Beta(\theta)$ prior with a single pseudocount parameter $\theta$; this corresponds to the $K2$ prior as defined by (Cooper and Herskovits, 1992) [4]. Other related $CPD$ priors such as the $BDeu$ score [3] are also possible.

Overall, the term $\alpha$ implicitly denotes all hyperparameters such as the edge penalty $\kappa$, the infeasible geometry penalty $\beta$, and the $CPD$ pseudocount $\theta$, in addition to other data such as the set of allowed edges and the infeasible geometries as determined from biophysical modeling.

**CRN proposal.** We are now ready to define the CRN proposal distribution $P_{pro}(CRN)$ introduced in the Metropolis-Hastings criterion of Eqn (13).

First, given the set of legal edge moves $M_E$, we uniformly propose an edge move with probability $\frac{1}{|M_E|}$. Here an edge move is either removal or addition of a single edge, and the move is legal if and only if: *i.* it does not induce a cycle and *ii.* is allowed according to biophysical perturbations showing it is a possible causal relation.

Next, given the proposed network edge structure $E'$ from the sampled edge move, we propose a new $CPD'$ set from the distribution of Eqn (15).

$$P(CPD'|E', \alpha) = \prod_{c' \in CPD'} Dir(\theta + X[c']) \tag{15}$$

where $X[c']$ is the corresponding *pseudo-population* counts for the conditional probability distribution $c'$ in the set $CPD'$ and $\theta$ is the prior pseudocount as previously defined by Eqn (14).

Therefore, the CRN forward proposal distribution is given by Eqn (16).

$$P_{pro}(CRN'|CRN) = \frac{1}{|M_E|} \cdot \prod_{c' \in CPD'} Dir(\theta + X[c']) \tag{16}$$

The CRN reverse proposal distribution is given by Eqn (17).

$$P_{pro}(CRN|CRN') = \frac{1}{|M_E|'} \cdot \prod_{c \in CPD} Dir(\theta + X[c]) \tag{17}$$

**CRN acceptance.** We can now revisit the CRN Metropolis-Hastings acceptance criterion of Eqn (18):

$$P_{acc}(CRN') = \min\{1, \frac{Q_{tar}(CRN')P_{pro}(CRN|CRN')}{Q_{tar}(CRN)P_{pro}(CRN'|CRN)}\} \tag{18}$$

Recall that $Q_{tar}(CRN)$ is defined as the product of the factor distributions $P(\{X_i\}|CRN)$ and $P(CRN|\alpha)$. The factor $P(\{X_i\}|CRN)$, as discussed in page 12, is simply a product of binomial distributions. The factor $P(CRN|\alpha)$ is given by Eqn (14). The forward and reverse $P_{pro}$ distributions are defined by Eqn (16) and Eqn (17), respectively.

For ease of notation, we first collect the penalty priors into the term $\Upsilon(CRN)$ given by Eqn (19).

$$\Upsilon(CRN) \equiv \kappa^{|E|} \cdot \exp\{-\beta \cdot energy(infeas)\} \tag{19}$$

The exact CRN Metropolis-Hastings acceptance criterion is now given by Eqn (20):

$$
\begin{aligned}
P_{acc}(CRN') &= \min\{1, \frac{Q_{tar}(CRN')P_{pro}(CRN|CRN')}{Q_{tar}(CRN)P_{pro}(CRN'|CRN)}\} \\
&= \min\{1, \frac{\Upsilon(CRN') \cdot \prod_{c' \in CPD'} Dir(\theta) \cdot P(\{X_i\}|CRN') \cdot P_{pro}(CRN|CRN')}{\Upsilon(CRN) \cdot \prod_{c \in CPD} Dir(\theta) \cdot P(\{X_i\}|CRN) \cdot P_{pro}(CRN'|CRN)}\} \\
&= \min\{1, \frac{\Upsilon(CRN') \cdot \prod_{c' \in CPD'} Dir(\theta) \cdot P(\{X_i\}|CRN') \cdot |M_{E'}|^{-1} \cdot \prod_{c \in CPD} Dir(\theta + X[c])}{\Upsilon(CRN) \cdot \prod_{c \in CPD} Dir(\theta) \cdot P(\{X_i\}|CRN) \cdot |M_E|^{-1} \cdot \prod_{c' \in CPD'} Dir(\theta + X[c'])}\}
\end{aligned} \tag{20}
$$

Eqn (20) may be considered unwieldy and computationally cumbersome. However, if we examine $Q_{tar}(CRN)$, we have:

$$
\begin{aligned}
Q_{tar}(CRN) &\propto P(CRN|\alpha) \cdot P(\{X_i\}|CRN) \\
&\propto \Upsilon(CRN) \cdot \prod_{c \in CPD} Dir(\theta) \cdot P(\{X_i\}|CRN) \\
&\propto \Upsilon(CRN) \cdot \prod_{c \in CPD} Dir(\theta + X[c]),
\end{aligned} \tag{21}
$$

we note that the expression $\prod_{c \in CPD} Dir(\theta) \cdot P(\{X_i\}|CRN)$ in Eqn (21) has been replaced by $\prod_{c \in CPD} Dir(\theta + X[c])$. This follows from the fact that the product of Dirchlet distributions is a *conjugate prior* to the product of binomial distributions in $P(\{X_i\}|CRN)$ [14]; hence, we can replace the expression with the analytical posterior and the proportional relation will still hold. Therefore, in Eqn (22), we offer the following approximate Metropolis–Hastings acceptance criterion which we use in practice:

$$
\begin{aligned}
P_{acc}(CRN') &= \min\{1, \frac{Q_{tar}(CRN')P_{pro}(CRN|CRN')}{Q_{tar}(CRN)P_{pro}(CRN'|CRN)}\} \\
&\approx \min\{1, \frac{\Upsilon(CRN') \cdot \prod_{c' \in CPD'} Dir(\theta + X[c']) \cdot P_{pro}(CRN|CRN')}{\Upsilon(CRN) \cdot \prod_{c \in CPD} Dir(\theta + X[c]) \cdot P_{pro}(CRN'|CRN)}\} \\
&\approx \min\{1, \frac{\Upsilon(CRN') \cdot \prod_{c' \in CPD'} Dir(\theta + X[c']) \cdot |M_{E'}|^{-1} \cdot \prod_{c \in CPD} Dir(\theta + X[c])}{\Upsilon(CRN) \cdot \prod_{c \in CPD} Dir(\theta + X[c]) \cdot |M_E|^{-1} \cdot \prod_{c' \in CPD'} Dir(\theta + X[c'])}\} \\
&\approx \min\{1, \frac{\Upsilon(CRN') \cdot |M_{E'}|^{-1} \cdot \prod_{c' \in CPD'} Dir(\theta + X[c']) \cdot \prod_{c \in CPD} Dir(\theta + X[c])}{\Upsilon(CRN)|M_E|^{-1} \cdot \prod_{c' \in CPD'} Dir(\theta + X[c']) \cdot \prod_{c \in CPD} Dir(\theta + X[c])}\} \\
&\approx \min\{1, \frac{\Upsilon(CRN')|M_{E'}|^{-1}}{\Upsilon(CRN)|M_E|^{-1}}\},
\end{aligned} \tag{22}
$$

where the product of Dirichlet distributions conveniently cancel from the acceptance ratio. The approximation, as opposed to equality, results from the ratio of normalization constants that may not directly cancel; however, since the proposed edge move is a small, local change (only a single edge may be removed or deleted from a sparse edge list), we believe the ratio of constants is likely to be inconsequential; an area of future research is to analytically derive and efficiently compute the ratio of normalization constants to allow an exact criterion.

##### 1.3.3 Markov updating the latent population of single-cell contact states

The target Gibbs conditional distribution, $P_{tar}$, for the latent population of single cell contact states $\{X_i\}$ (*i.e.*, pseudo-population) is given by Eqn (23). As before, we use integration $\int P(\ldots, V)dV$ to universally denote marginalization of a random variable $V$ regardless of its domain:

$$\begin{aligned}
P_{tar}(\{X_i\}) &= P(\{X_i\}|CRN, Hi\text{-}C, \alpha) \\
&= \frac{P(CRN, \{X_i\}, Hi\text{-}C|\alpha)}{\int P(CRN, \{X_i\}, Hi\text{-}C|\alpha)d\{X_i\}} \\
&= \frac{P(CRN|\alpha)P(\{X_i\}|CRN)P(Hi\text{-}C|\{X_i\})}{\int P(CRN|\alpha)P(\{X_i\}|CRN)P(Hi\text{-}C|\{X_i\})d\{X_i\}} \\
&= \frac{P(CRN|\alpha)P(\{X_i\}|CRN)P(Hi\text{-}C|\{X_i\})}{P(CRN|\alpha)\int P(\{X_i\}|CRN)P(Hi\text{-}C|\{X_i\})d\{X_i\}} \\
&= \frac{P(\{X_i\}|CRN)P(Hi\text{-}C|\{X_i\})}{\int P(\{X_i\}|CRN)P(Hi\text{-}C|\{X_i\})d\{X_i\}} \\
&\propto P(\{X_i\}|CRN)P(Hi\text{-}C|\{X_i\}) = Q_{tar}(\{X_i\}),
\end{aligned} \tag{23}$$

where the normalization constant $Z_{\{X_i\}}$ is given by Eqn (24):

$$Z_{\{X_i\}} = \int P(\{X_i\}|CRN)P(Hi\text{-}C|\{X_i\})d\{X_i\}, \tag{24}$$

which will cancel from the *Metropolis* acceptance criterion.

Note, we assume the random variable $Hi\text{-}C$ has been experimentally measured (*i.e.*, has been *observed*) with value $M_H$. Therefore, the unnormalized (*i.e.*, improper) target Gibbs distribution, $Q_{tar}(\{X_i\}$, is more specifically given by Eqn (25):

$$Q_{tar}(\{X_i\}) = P(\{X_i\}|CRN)P(Hi\text{-}C = M_H|\{X_i\}) \tag{25}$$

**Hi–C factor distribution.** The factor distribution $P(\{X_i\}|CRN)$ for the latent population of single-cell contact states (*i.e.*, pseudo-population), as discussed in page (1.3.2), is simply a product of binomial distributions. We define the Hi–C factor distribution $P(Hi\text{-}C = M_H|\{X_i\}$ by Eqn.˜(26):

$$P(Hi\text{-}C = M_H|\{X_i\} \propto \exp\{\xi \cdot corr(M_H, \{X_i\})\} \tag{26}$$

where the hyper-parameter $\xi \in \mathbb{R}_{\geq 0}$ and $corr(M_H, \{X_i\})$ is a similarity score, *e.g.*, Pearson correlation, between the measured Hi–C value $M_H$ and the pseudo-population $\{X_i\}$.

**Proposal distribution for the latent population of single-cell contact states.** For the proposal distribution of the latent population of single-cell contact states (*i.e.* pseudo-population), we use a Polya Urn sampling scheme to simulate draws from a Beta-Bernoulli distribution[13]. Specifically, each row of the of the pseudo-population matrix $\{X_i\}$ is drawn independently from a Beta-Bernoulli distribution with expectation set to the corresponding row frequency in the measured Hi–C matrix $M_H$.

**Acceptance criterion for the latent population of single-cell contact states.** We use a simplified *Metropolis* acceptance criterion, since the pseudo-population proposal distribution is symmetric, *i.e.*, the forward and reverse proposal distributions are the same. The pseudo-population Metropolis acceptance criterion is given by Eqn (27):

$$\begin{aligned} P_{acc}(X_i{}') &= \min\{1, \frac{Q_{tar}(\{X_i\}')}{Q_{tar}(\{X_i\})}\} \\ &= \min\{1, \frac{P(\{X_i\}'|CRN)P(Hi\text{-}C = M_H|\{X_i\}')}{P(\{X_i\}|CRN)P(Hi\text{-}C = M_H|\{X_i\})}\}, \end{aligned} \tag{27}$$

where the factors $P(\{X_i\}|CRN)$ and $P(Hi\text{-}C|\{X_i\})$ are defined in page 1.3.2 and 26 respectively.

Empirically, we have found that proposing a full pseudo-population update will result in a low acceptance ratio and poor mixing. Therefore, it is prudent to only propose updating a small subset of the pseudo-population $\{X_k\} \subset \{X_i\}$. The subset $\{X_k\}$ are uniform randomly selected from $\{X_i\}$; therefore the proposal is still symmetric and the Metropolis criterion can still be used.

Furthermore, the proposal distribution uses a *rejection sampling* approach to avoid proposing samples with known infeasible geometry. That is, if a proposed $X_k$ is infeasible, then it is simply redrawn. Therefore, the proposal distribution is a *truncated* Beta-Bernoulli such that infeasible samples have zero probability mass.

Overall, the CHROMATIX framework incorporates both a fractal Monte Carlo physical folding algorithm and a Bayesian generative MCMC approach for deconvolving population-based Hi-C data into single-cell contact states. The Bayesian generative model utilizes biophysical simulation data to build a sparsity-inducing prior on the allowable folding mechanisms and co-occurring contact states; it can then perform posterior inference on the set of single-cell contact states given the measured population Hi-C. Results of applying CHROMATIX on the 39 loci studied here showed that models generated for 38 loci have higher posterior probabilities than a simple naive model of product of independent pairwise contacts. However, there may exist genomic regions where pairwise contacts are largely independent. For example, we find that the 1.7 MB genomic region (chr5:88,020,000–89,700,000) has few dependencies in the final folding model: about 80% of the coarse-grained specific contacts are likely acting independently.

#### 2 Genomic Loci

All TAD-bounded loci (< 2 MB) with $\geq 2$ super-enhancers with SPRITE evidence for possible SE condensation in the GM12878 mammalian cell line are included in this study. The size of the loci range from 480 KB to 1.84 MB. These loci are modeled at the resolution of 5 KB.

Details of these loci can be found in Table S1.

#### 3 CHROMATIX 3-D Chromatin Reconstruction

At each of the 39 genomic loci, the total number of unique experimentally measured $(i, j)$ Hi-C contacts ranges from 2,414 to 62,785; among these contacts, the number identified as specific ranges from 301 to 2,112. The number of specific $(i, j)$ interactions used to generate folded 3-D chromatin ensembles of the loci range from 15 to 35. The fraction of specific Hi-C contacts among all measured Hi-C contacts ranges from 1.8% to 12.5%. The fraction of retained specific Hi-C interactions for 3-D ensemble folding among all identified specific Hi-C interactions ranges from 0.7% to 10.6%. The corresponding fraction of retained specific Hi-C interactions for 3-D ensemble folding among all measured Hi-C interactions ranges from 0.023% to 1.33%.

The average specific contact retention fraction of 5% results from clustering with a fixed cap of at most 35 retained specific contacts per loci. This cap was chosen arbitrarily based on an initial trial run with a single locus, in which high quality reconstruction was obtained. No further attempt was made to optimize this fraction, as it affords the necessary computational cost required for large scale perturbation simulations.

Figure 9 shows the correlation between the simulated and experimental Hi-C data sets vs. the percentage of retained specific contacts among the 39 loci.

The Pearson correlation (orange) appears to be fairly stable, with only minimal decrease as the fraction of retained specific contacts decreases. In contrast, distance-corrected Pearson correlation (blue) appears to be more sensitive to a reduced specific contact retention fraction, with a steeper slope of the trend line (blue dash) compared to the original Pearson correlation trend line (orange dash).

For the 39 loci, the mean number of likely infeasible contact combinations is 255 combinations per locus among the coarse-grained specific contacts.

**Confining volume.** The random ensembles for a 2 MB region are generated using 400 monomers at 5 KB resolution. To determine the confining nuclear volume, we scale the experimentally measured nuclear diameter for the GM12878 cell line [23] to preserve constant base pair density ($\frac{bp}{nm^3}$). Specifically, from the following relation:

$$\frac{TotalGenomeLengthInBasePairs}{NuclearVolumeInNm^3} = \frac{LociLenthInBasePairs}{ScaledNuclearVolumeInNm^3} \quad (28)$$

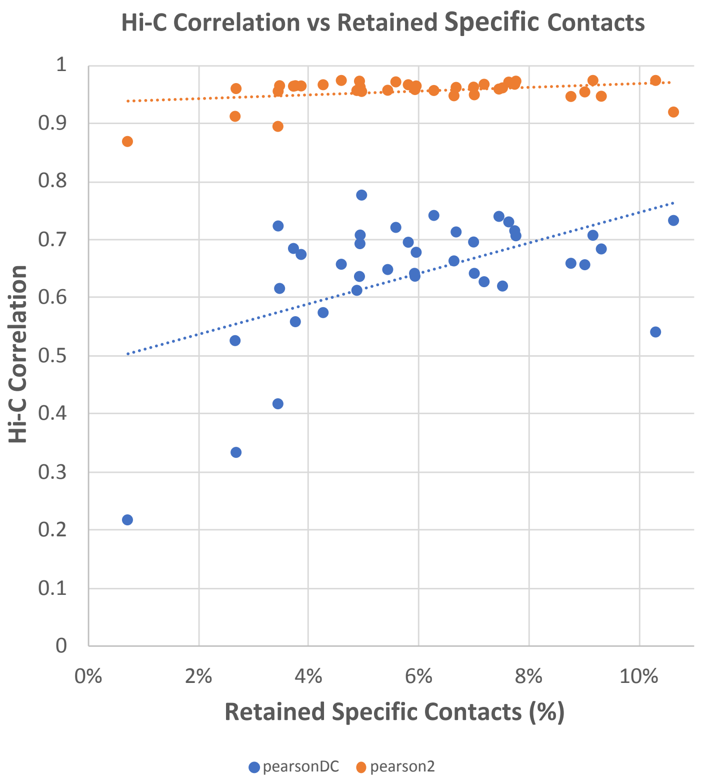

Figure 9: **Correlation between the simulated and experimental Hi-C data sets vs. the % of retained specific contacts** among the 39 loci. The orange dots (pearson2) are the Pearson correlations after removal of the first two diagonals from the Hi-C matrix. The blue dots (pearsonDC) are the corresponding distance-corrected Pearson correlation [1]. This figure was generated from the 'kf', 'pearson2', and 'pearsonDC' columns provided in supplementary Table S1.

we solve for $ScaledNuclearVolumeInNm^3$. Given the scaled nuclear volume, we can model the confining space as a sphere with the corresponding diameter $D$ given by relation:

$$ScaledNuclearVolumeInNm^3 = \frac{\pi D^3}{6} \tag{29}$$

For loci $< 2$ MB, we still simulate confinement and volume exclusion for a 2 MB region, but use the first $x$ beads, where $x$ is the size of the locus in units of 5 KB, as the relevant 3-D conformation of the modeled locus.

Examples of 3-D self-avoiding random chromatin polymers confined in nuclear volume are shown in Figure S1. Examples of reconstructed 3-D chromatin polymers from inferred single-cell states for locus chr 12: 11690000 – 12210000 are shown in Figure S2.

### 4 Comparison with single-cell studies

We have obtained publicly available single-cell Dip-C data of GM12878 cells from the GEO database (GSE117874) [25], and compared to our CHROMATIX cells. There are 17 Dip-C cells reported, two are damaged and thus removed. Each remaining Dip-C cell was then split into maternal and paternal homologs, so the Dip-C ensemble was effectively doubled to $2 \times 15$. We then examined the 39 loci in each of the 30 Dip-C cells. After removal of cells with $< 10$ contacts, we obtain a set of $n = 976$ Dip-C single-cell models for these 39 loci.

We calculate the overlap coefficient as [26]:

$$Overlap = \frac{|X \cap Y|}{\min(|X|,\, |Y|)}$$

where $X$ is the set of pairwise contacts within the Dip-C cell and $Y$ is the set of pairwise contacts within the CHROMATIX cell.

### 5 Comparison with SPRITE data

The SPRITE data is sparse, as it was designed for genome-wide study, which accounts for the relatively low coverage percentages in Figure S5. Furthermore, analysis of SPRITE data is typically at a coarser resolution of ∼1 MB. In contrast, our analysis of Super-Enhancer, Enhancer, and Promoter higher-order structures requires much finer resolution, at minimum 5 KB as in this study. However, as the SPRITE technique can capture higher-order interactions and SPRITE data for the GM12878 cell line is publicly available, it is therefore useful for us to compare our results with the SPRITE data.

The percentages shown in Figure S5 are over all SPRITE clusters located within the corresponding 500 KB to 2 MB TAD locus. This averages to ∼3,800 SPRITE clusters per locus of varying sizes ( *e.g.* from 3, 4, 5, $\cdots$ ). For a 500 KB TAD at 5KB resolution, there are $100 \times 100$ possible loop anchor locations.

In addition, the many-body sizes range from 3, 4, 5, 6, and beyond. When the 3,800 SPRITE clusters are distributed over $100\times$ 100 anchor positions and at a specific many-body size, the curse-of-dimensionality leads to overall sparse SPRITE data for analysis.

Although the SPRITE coverage itself is low due to this sparsity, we are able to detect statistically significant trends among specific vs. non-specific principal loops. That is, these trends, which we call *SPRITE concordance measures*, are quite robust in capturing the significant pattern of enrichment of specific principal loops relative to non-specific principal loops in the majority of loci modeled.

**Control for genomic distance.** Our model *controls for genomic distance* by building a separate random contact model at each $|j-i|$ genomic distance bin. We identify an $(i, j)$ contact as specific based on the distribution of *random contacts with the same* $|j-i|$ *genomic separation.*

Therefore, an $(i, j)$ contact that is called specific will be more common in our CHROMATIX ensemble than non-specific contacts with the same $|j-i|$ genomic distance, because we are assigning $p$-values based on the upper-tail of the corresponding $|j-i|$ random contact distribution. This holds true regardless of whether or not $|j-i|$ is a short interval.

Since there are relatively more specific principal loops by our model at any given $|j-i|$, we ask if the subset of our principal loops that show up in SPRITE data also exhibit this trend. We therefore examine the coverage of principal loops of many-bodies in our model by SPRITE data. It is reasonable to expect that more specific principal loops from our model are present in SPRITE data compared to corresponding non-specific principal loops at each $|j-i|$ bin.

**Concordance with SPRITE.** In Fig 10, the leftmost plot illustrates perfect SPRITE concordance. Here all specific principal loops (blue dots) have higher SPRITE coverage than non-specific principal loops (pink dots) at the same $|j-i|$ genomic distance.

The middle plot illustrates perfect discordance with SPRITE. Here the relationship is reversed and all specific principal loops have lower SPRITE coverage than corresponding non-specific principal loops.

The rightmost plot shows when the SPRITE coverage is mixed across principal loop classes. This is more reflective of real experimental data.

To quantify the SPRITE concordance of each of these plots, we compute the median specific SPRITE coverages at each genomic distance bin and compare to the corresponding median non-specific SPRITE coverages.

We then ask how likely we are to observe a given fraction of specific median coverages exceeding the corresponding non-specific median coverages, if we assume a null hypothesis where specific and non-specific labels do not matter. In other words, *our null hypothesis assumes there is no relationship between our*

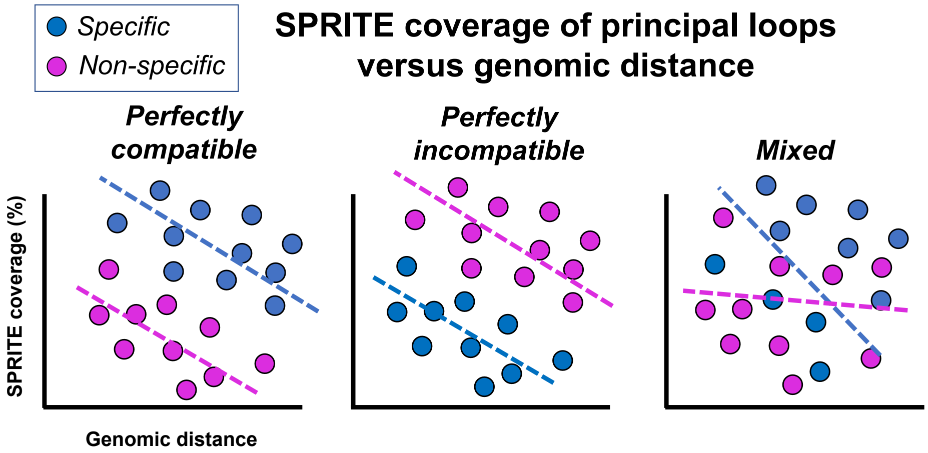

Figure 10: **Illustration of SPRITE concordance.** The trends of perfectly compatible *vs* incompatible *vs* mixed SPRITE coverage scenarios are shown. Here each dot is an $(i, j)$ principal loop with: 1) $x$-coordinate $= |j - i|$ 2). $y$-coordinate = corresponding SPRITE coverage for principal loop $(i, j)$. This is defined as percentage of compatible SPRITE clusters containing principal loop $(i, j)$, where compatible clusters are those with sizes $\geq k$ of the $k$-body containing the principal loop. The color distinguishes specific vs. non-specific principal loops according to our random polymer folding model.

*results and SPRITE data.*

To test the null hypothesis, we randomly permute the non-specific and specific labels and recompute the fraction of specific median coverages exceeding the corresponding non-specific median coverages using the permuted labels. We repeat the permutation procedure for 1,000 times, and then we assign an upper tail $p$-value to the observed fraction based on this null distribution. If the observed p-value is significant under the null distribution, then we reject the null hypothesis and simply state that the data is concordant with SPRITE.

### 6 Supplementary Figures

#### 6.1 Figure S1.

Examples of random, self-avoiding folded polymers under spherical confinement. Approximately 400,000 random polymers were sampled using fractal Monte Carlo to generate a null distribution over random pairwise contacts.

**FIGURE S1**

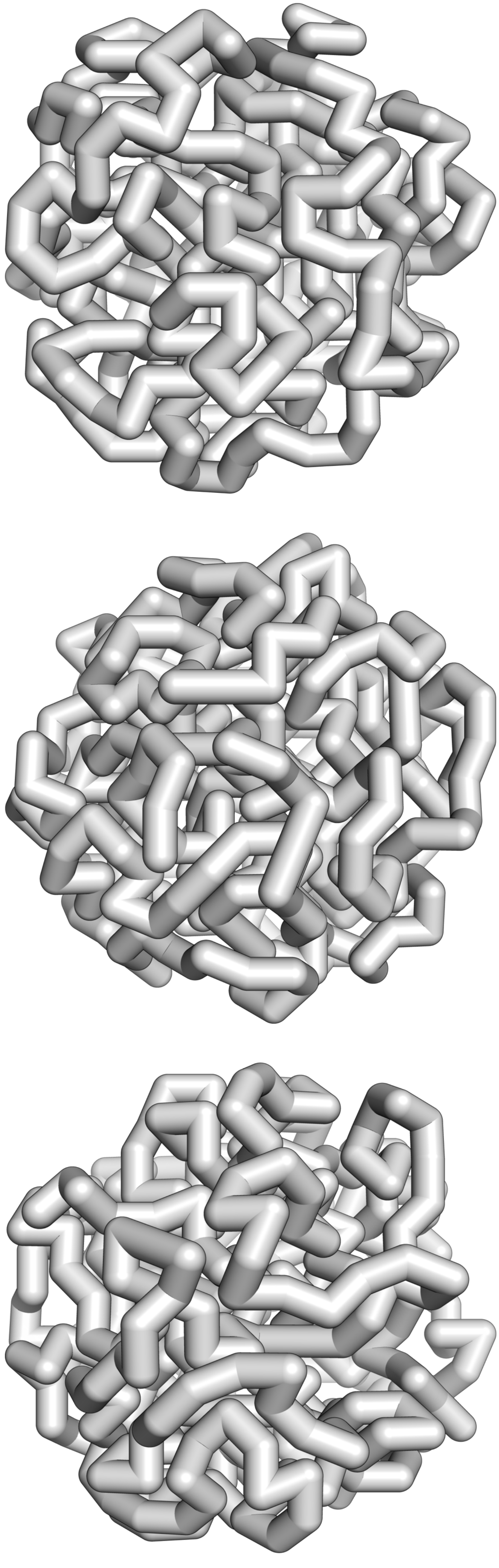

#### 6.2 Figure S2.

Examples of constrained, self-avoiding polymers under spherical confinement with coloring corresponding to the relative base pair position along the genome. Approximately 25,000 unique 3-D polymers were sampled at each of the 39 modeled loci in order to reconstruct the simulated Hi-C ensembles.

**FIGURE S2**

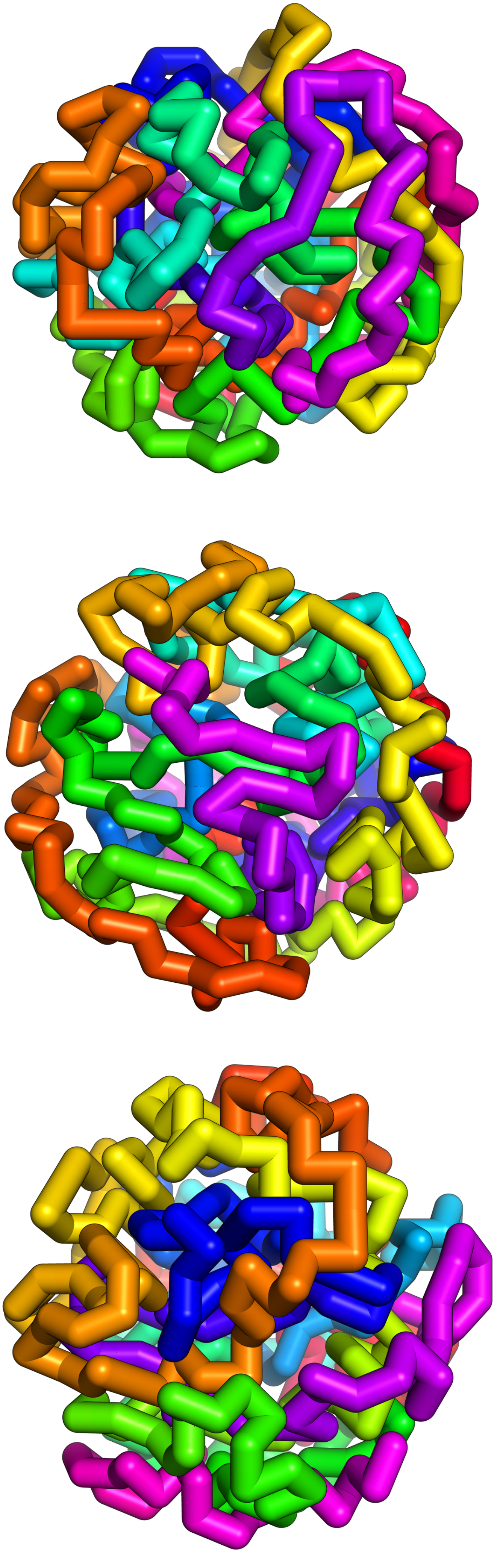

#### 6.3 Figure S3.

Details of the reconstructed ensembles of all loci are shown in the heatmaps of Figure S3. In each plot, the upper triangle shows measured Hi-C frequencies of the locus, after KR-normalization and then quantile normalization against the random ensemble as described in main text.

The lower triangle shows the simulated Hi-C frequencies from aggregation of 3-D polymer folds (see main text). The axes are in units of 5 KB intervals. The location of the locus, the total number of measured Hi-C contacts (*ntot*) of the locus, the total number of called specific Hi-C contacts (*nspec*), the number of specific contacts retained after coarse-graining that are used in constructing the 3D ensembles ($k$), and the Pearson correlation without diagonal removal (*pearson0*) and with 3 diagonals removed (*pearson3*) are all provided for each locus.

Note that for loci: chr 1: 234635000–235415000, chr 9: 4755000–5730000, chr 12: 11690000–12210000, chr 14: 68285000–69330000, the Pearson correlations shown are averages of two replicated simulations, one of which is depicted here.

**FIGURE S3**

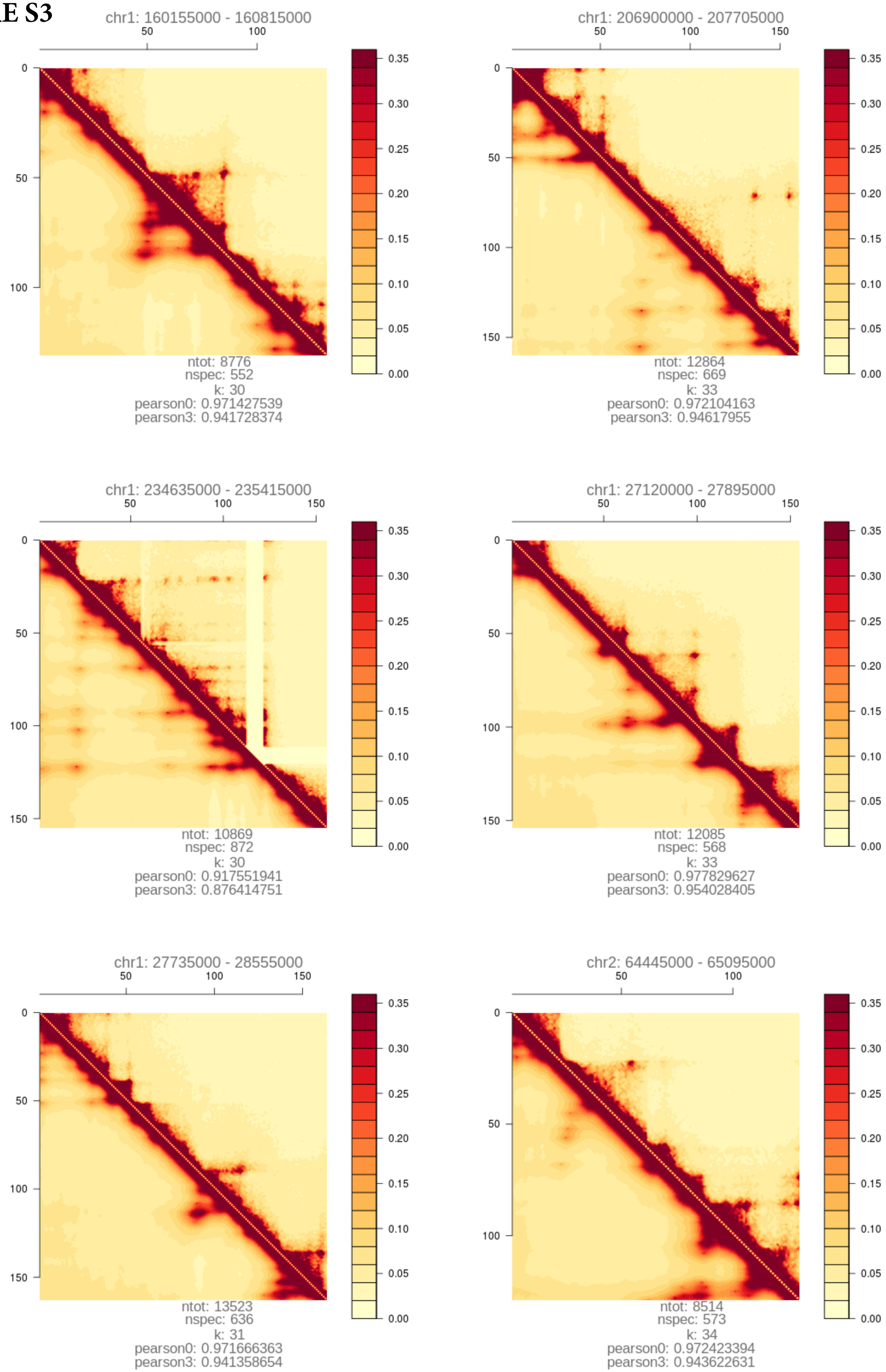
chr1: 160155000 - 160815000
ntot: 8776
nspec: 552
k: 30
pearson0: 0.971427539
pearson3: 0.941728374
chr1: 206900000 - 207705000
ntot: 12864
nspec: 669
k: 33
pearson0: 0.972104163
pearson3: 0.94617955
chr1: 234635000 - 235415000
ntot: 10869
nspec: 872
k: 30
pearson0: 0.917551941
pearson3: 0.876414751
chr1: 27120000 - 27895000
ntot: 12085
nspec: 568
k: 33
pearson0: 0.977829627
pearson3: 0.954028405
chr1: 27735000 - 28555000
ntot: 13523
nspec: 636
k: 31
pearson0: 0.971666363
pearson3: 0.941358654
chr2: 64445000 - 65095000
ntot: 8514
nspec: 573
k: 34
pearson0: 0.972423394
pearson3: 0.943622631
0.35
0.30
0.25
0.20
0.15
0.10
0.05
0.00

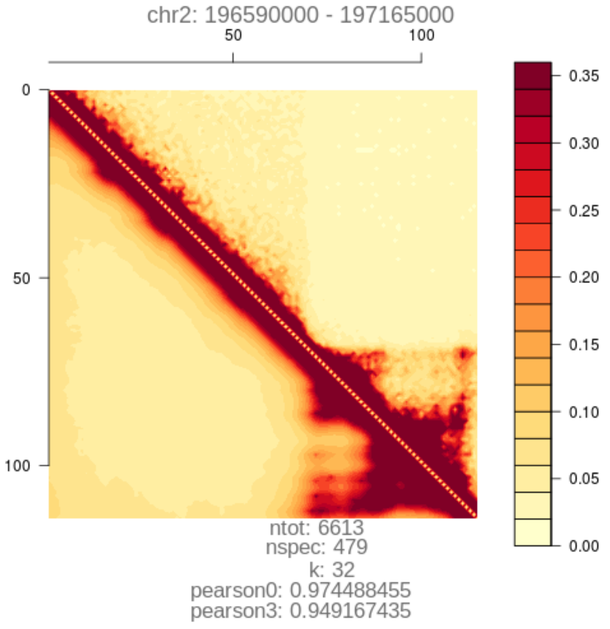
chr2: 196590000 - 197165000
ntot: 6613
nspec: 479
k: 32
pearson0: 0.974488455
pearson3: 0.949167435

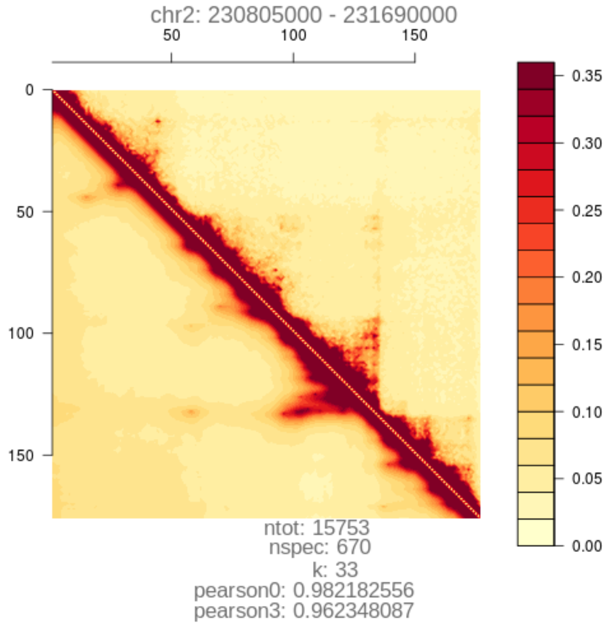
chr2: 230805000 - 231690000
ntot: 15753
nspec: 670
k: 33
pearson0: 0.982182556
pearson3: 0.962348087

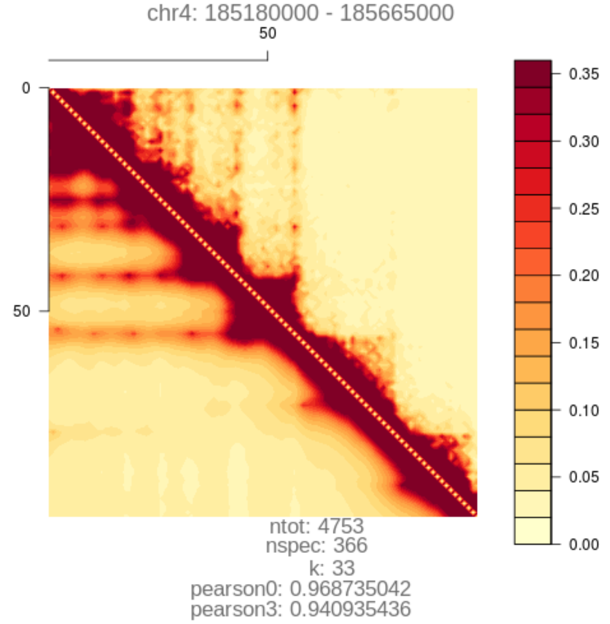
chr4: 185180000 - 185665000
ntot: 4753
nspec: 366
k: 33
pearson0: 0.968735042
pearson3: 0.940935436

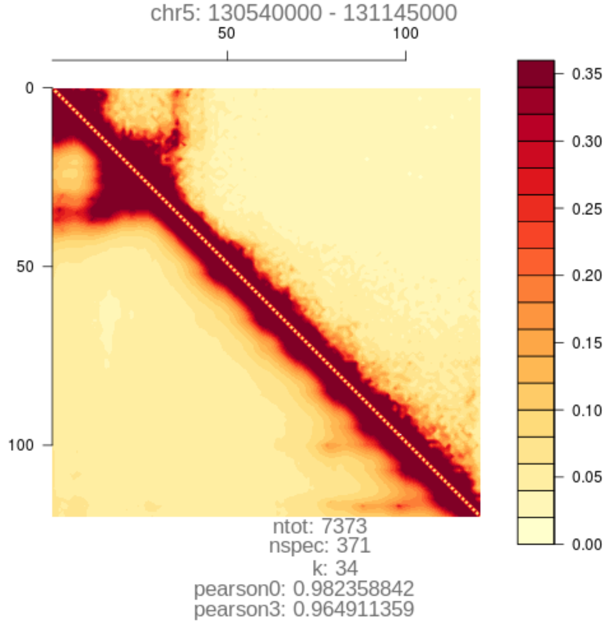
chr5: 130540000 - 131145000
ntot: 7373
nspec: 371
k: 34
pearson0: 0.982358842
pearson3: 0.964911359

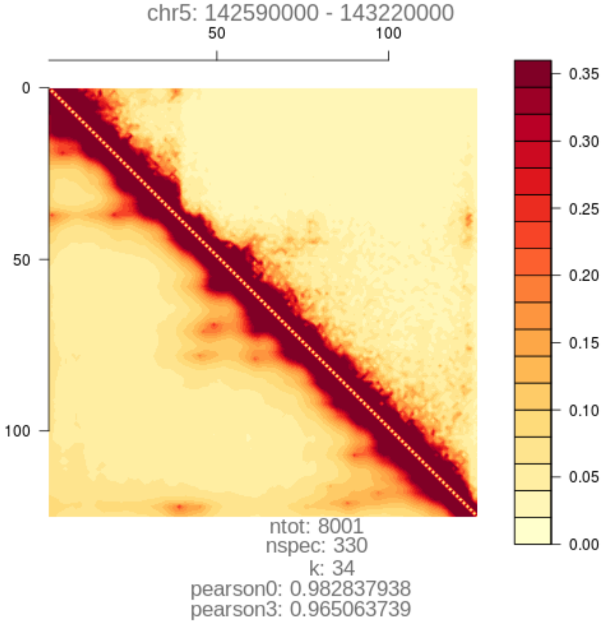
chr5: 142590000 - 143220000
ntot: 8001
nspec: 330
k: 34
pearson0: 0.982837938
pearson3: 0.965063739

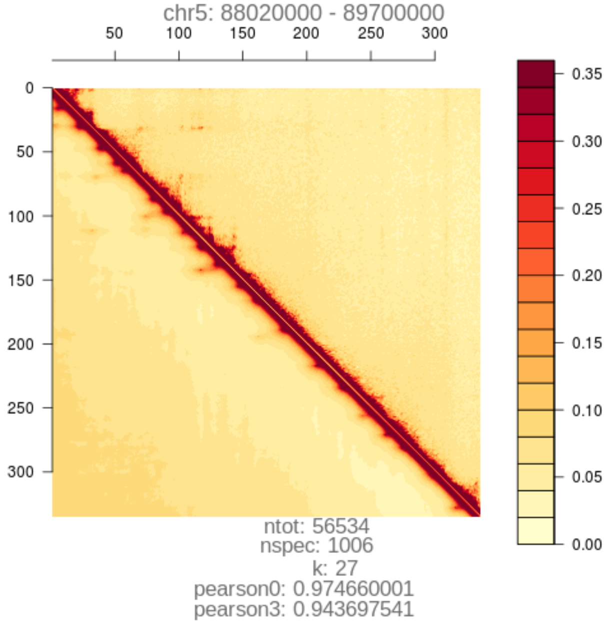
chr5: 88020000 - 89700000
ntot: 56534
nspec: 1006
k: 27
pearson0: 0.974660001
pearson3: 0.943697541

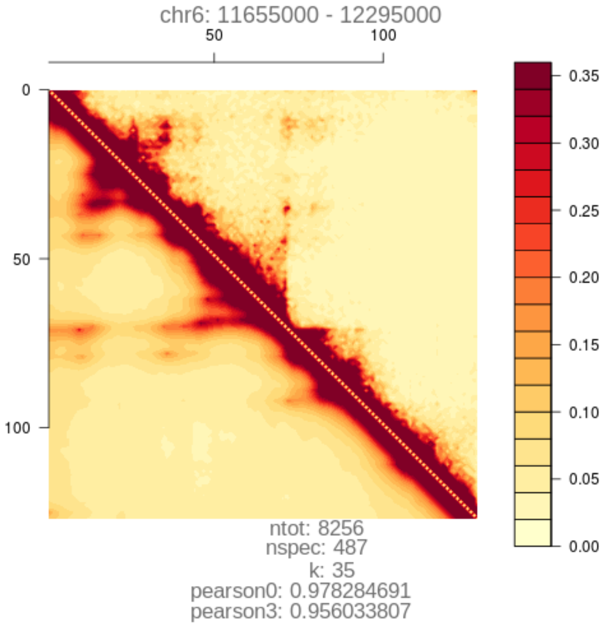
chr6: 11655000 - 12295000
ntot: 8256
nspec: 487
k: 35
pearson0: 0.978284691
pearson3: 0.956033807

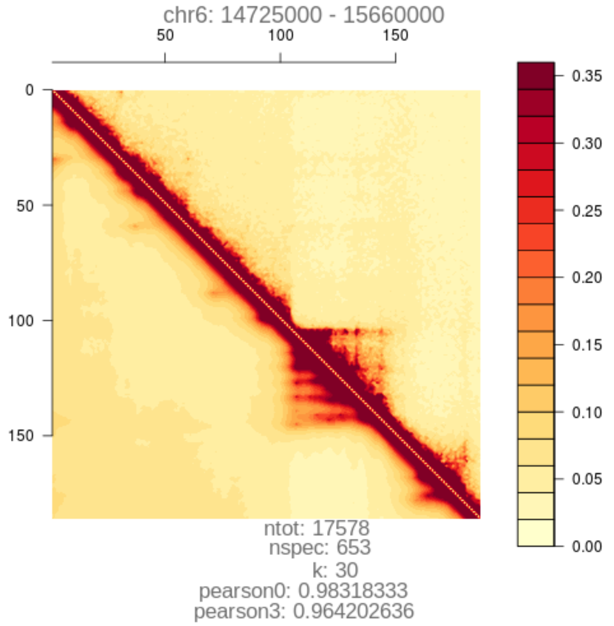
chr6: 14725000 - 15660000
ntot: 17578
nspec: 653
k: 30
pearson0: 0.98318333
pearson3: 0.964202636

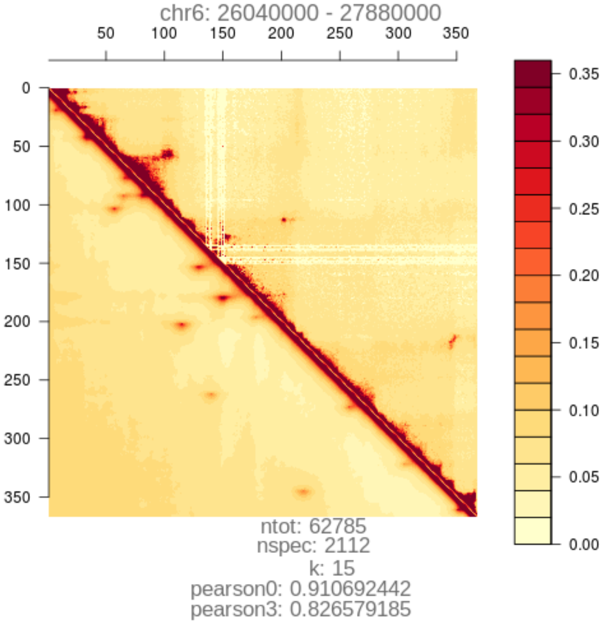
chr6: 26040000 - 27880000
ntot: 62785
nspec: 2112
k: 15
pearson0: 0.910692442
pearson3: 0.826579185

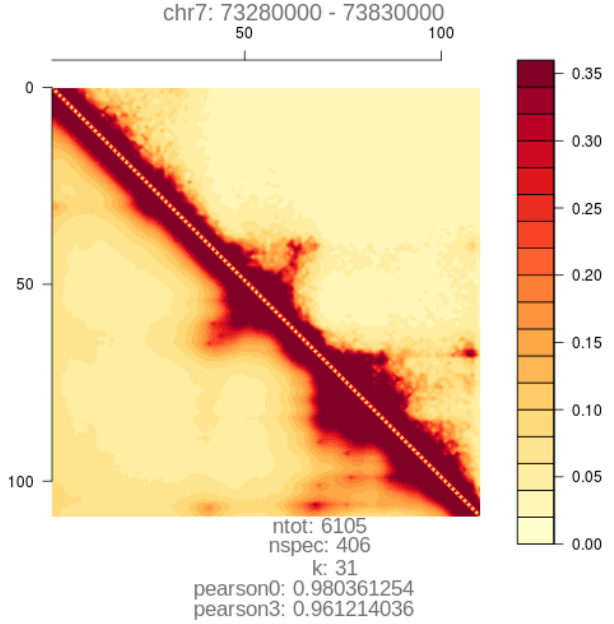
chr7: 73280000 - 73830000
ntot: 6105
nspec: 406
k: 31
pearson0: 0.980361254
pearson3: 0.961214036

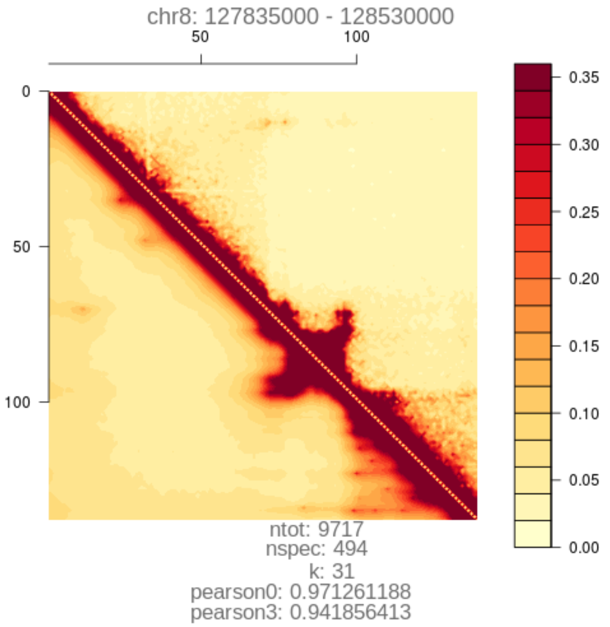
chr8: 127835000 - 128530000
ntot: 9717
nspec: 494
k: 31
pearson0: 0.971261188
pearson3: 0.941856413

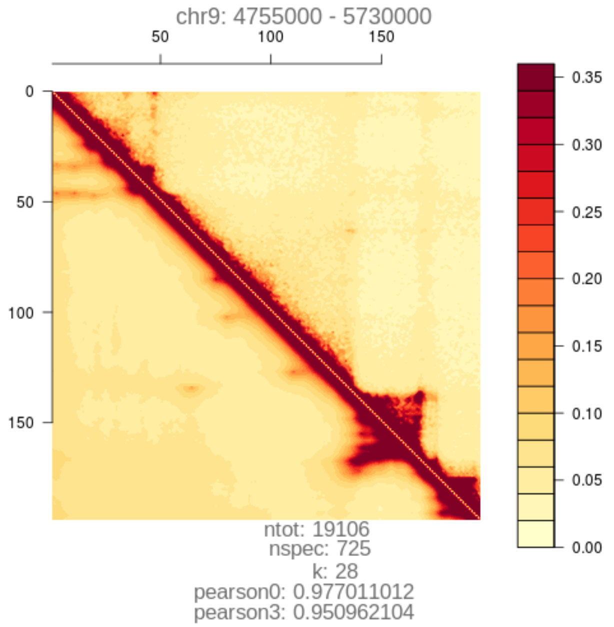
chr9: 4755000 - 5730000
ntot: 19106
nspec: 725
k: 28
pearson0: 0.977011012
pearson3: 0.950962104

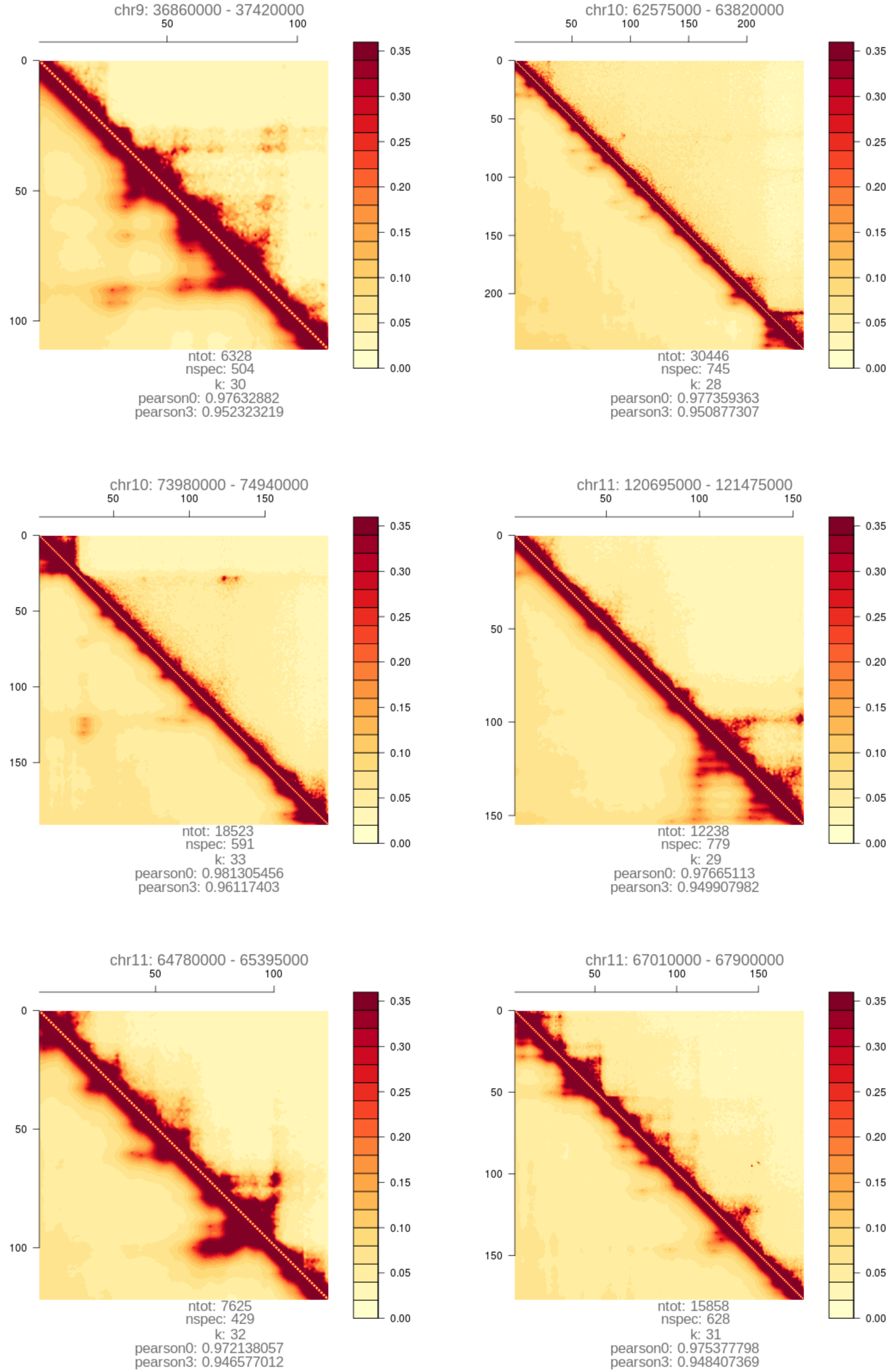
chr9: 36860000 - 37420000
ntot: 6328
nspec: 504
k: 30
pearson0: 0.97632882
pearson3: 0.952323219
chr10: 62575000 - 63820000
ntot: 30446
nspec: 745
k: 28
pearson0: 0.977359363
pearson3: 0.950877307
chr10: 73980000 - 74940000
ntot: 18523
nspec: 591
k: 33
pearson0: 0.981305456
pearson3: 0.96117403
chr11: 120695000 - 121475000
ntot: 12238
nspec: 779
k: 29
pearson0: 0.97665113
pearson3: 0.949907982
chr11: 64780000 - 65395000
ntot: 7625
nspec: 429
k: 32
pearson0: 0.972138057
pearson3: 0.946577012
chr11: 67010000 - 67900000
ntot: 15858
nspec: 628
k: 31
pearson0: 0.975377798
pearson3: 0.948407369

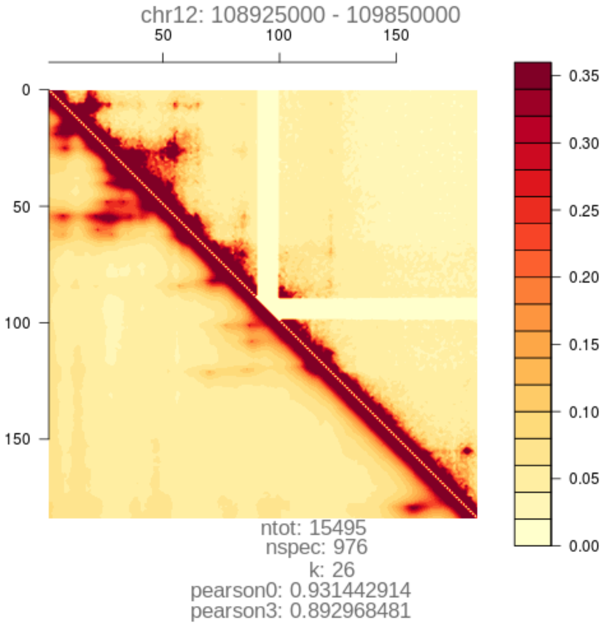
chr12: 108925000 - 109850000
ntot: 15495
nspec: 976
k: 26
pearson0: 0.931442914
pearson3: 0.892968481

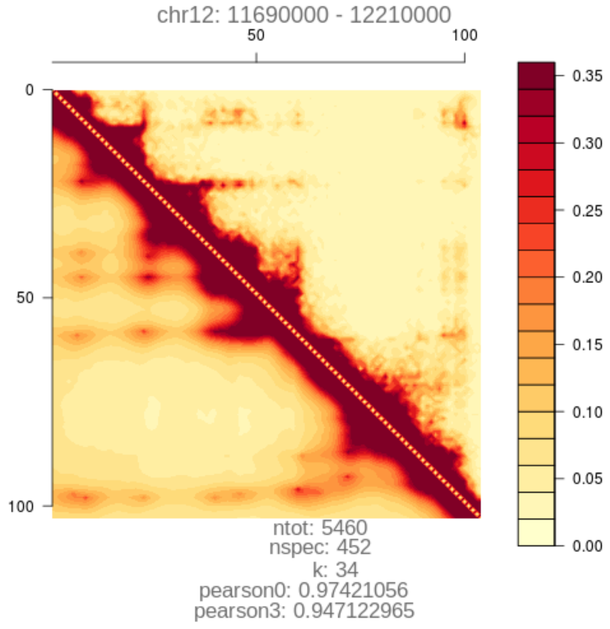
chr12: 11690000 - 12210000
ntot: 5460
nspec: 452
k: 34
pearson0: 0.97421056
pearson3: 0.947122965

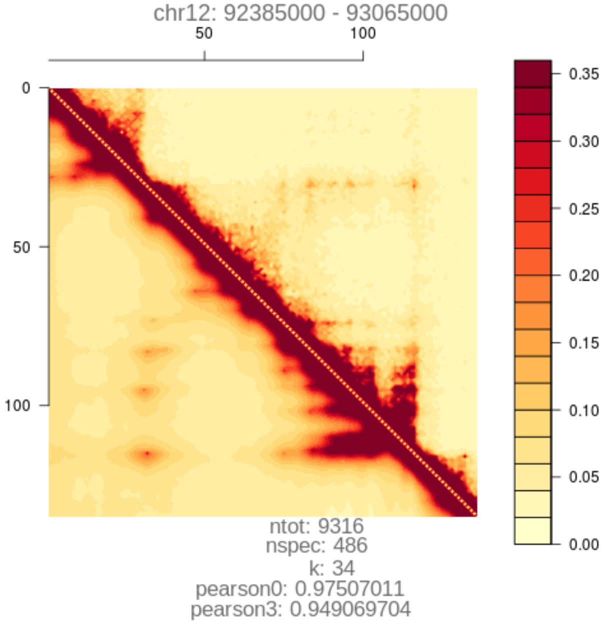
chr12: 92385000 - 93065000
ntot: 9316
nspec: 486
k: 34
pearson0: 0.97507011
pearson3: 0.949069704

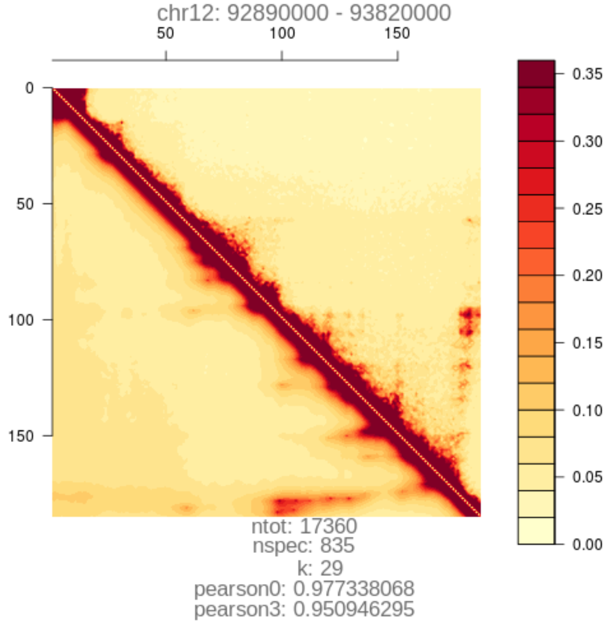
chr12: 92890000 - 93820000
ntot: 17360
nspec: 835
k: 29
pearson0: 0.977338068
pearson3: 0.950946295

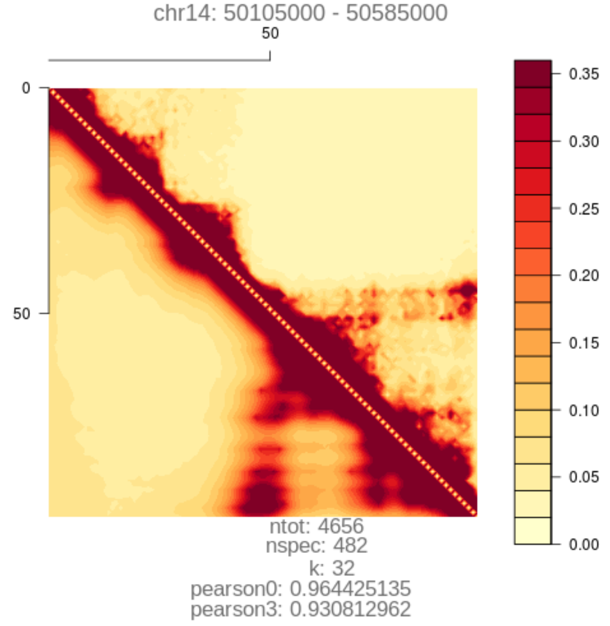
chr14: 50105000 - 50585000
ntot: 4656
nspec: 482
k: 32
pearson0: 0.964425135
pearson3: 0.930812962

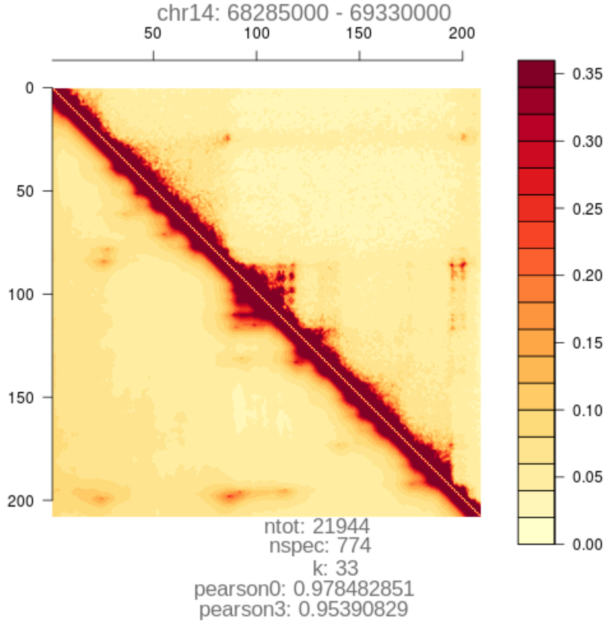
chr14: 68285000 - 69330000
ntot: 21944
nspec: 774
k: 33
pearson0: 0.978482851
pearson3: 0.95390829

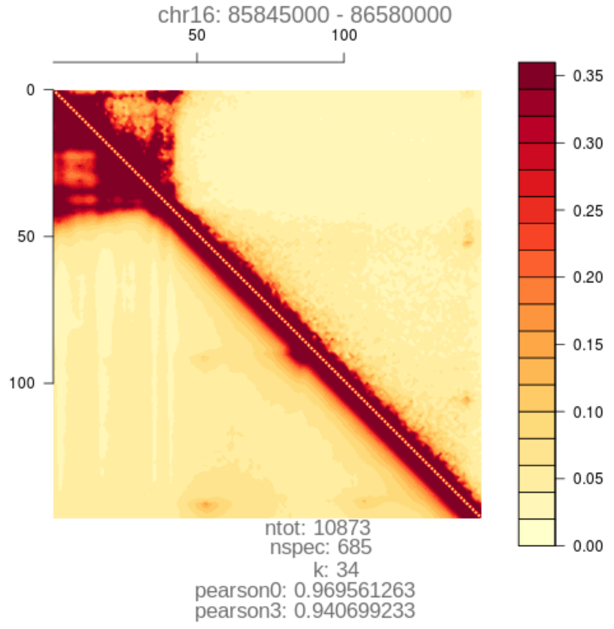
chr16: 85845000 - 86580000
ntot: 10873
nspec: 685
k: 34
pearson0: 0.969561263
pearson3: 0.940699233

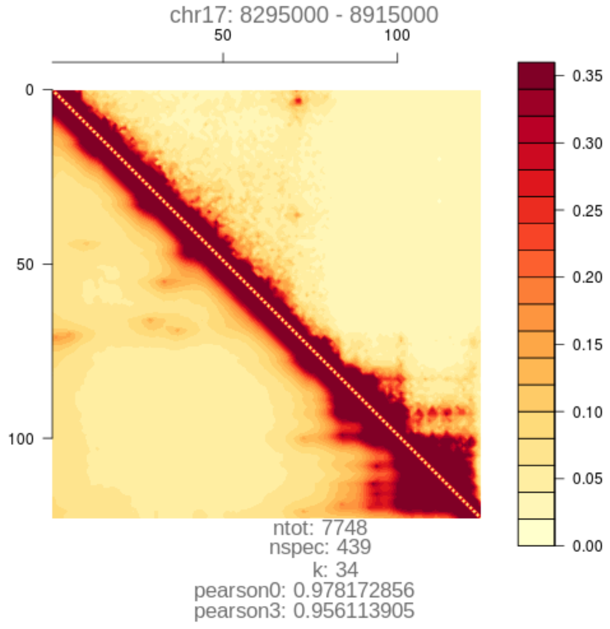
chr17: 8295000 - 8915000
ntot: 7748
nspec: 439
k: 34
pearson0: 0.978172856
pearson3: 0.956113905

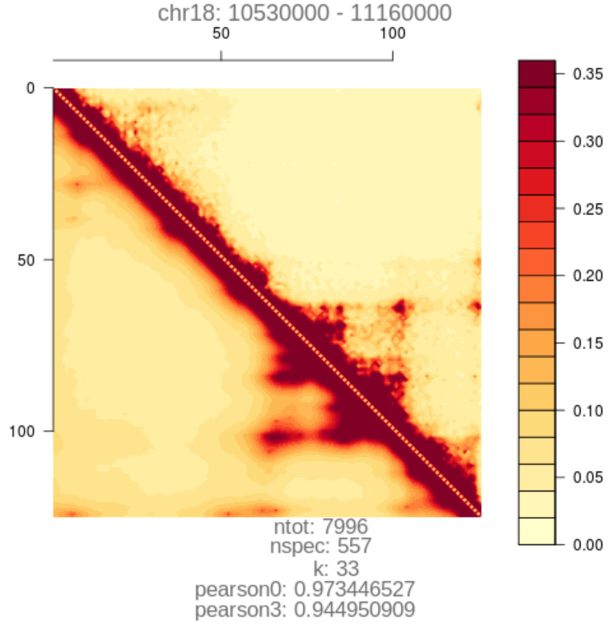
chr18: 10530000 - 11160000
ntot: 7996
nspec: 557
k: 33
pearson0: 0.973446527
pearson3: 0.944950909

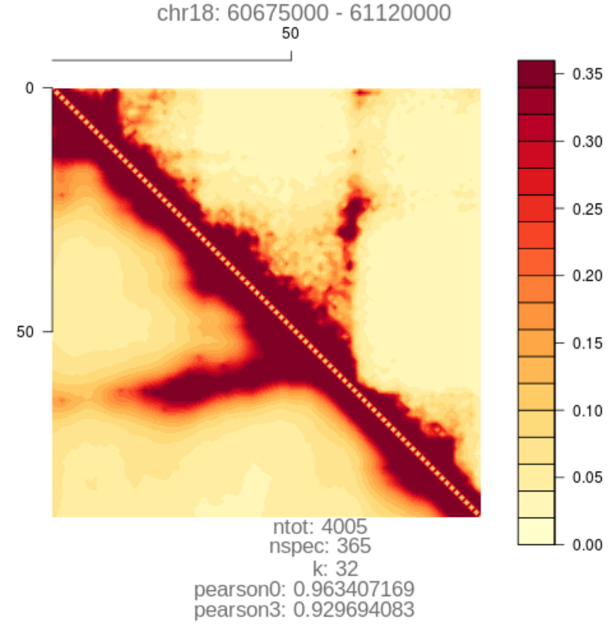
chr18: 60675000 - 61120000
ntot: 4005
nspec: 365
k: 32
pearson0: 0.963407169
pearson3: 0.929694083

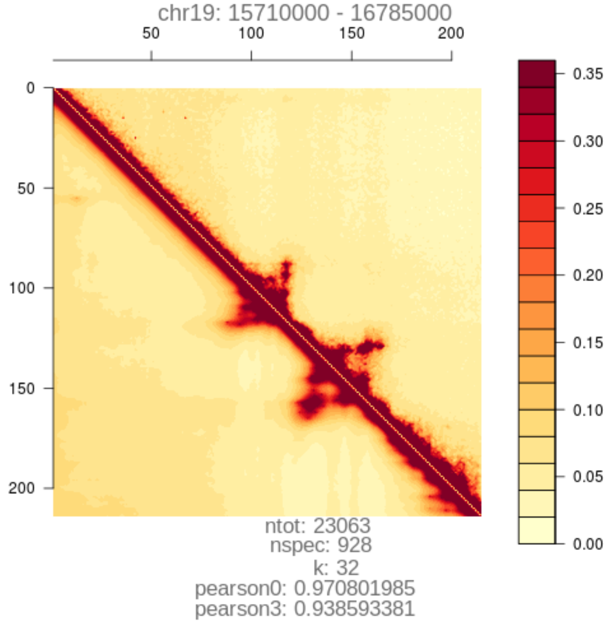
chr19: 15710000 - 16785000
ntot: 23063
nspec: 928
k: 32
pearson0: 0.970801985
pearson3: 0.938593381

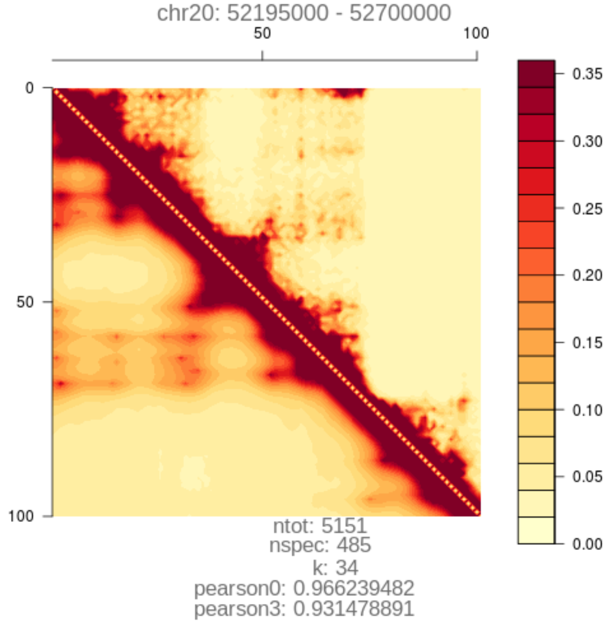
chr20: 52195000 - 52700000
ntot: 5151
nspec: 485
k: 34
pearson0: 0.966239482
pearson3: 0.931478891

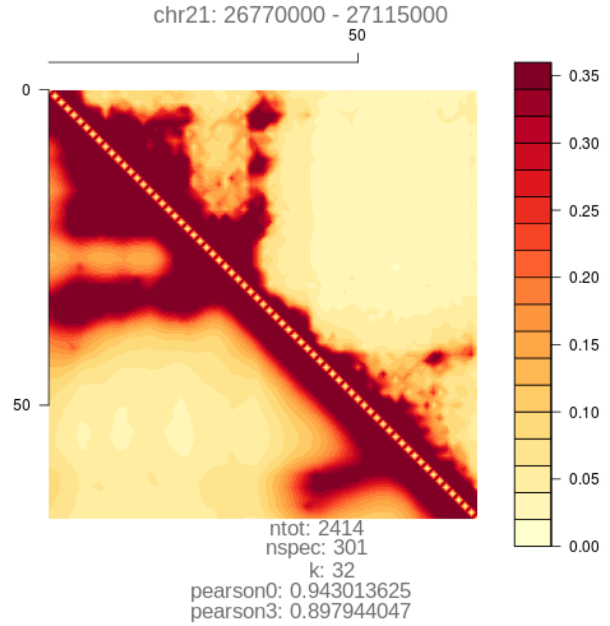
chr21: 26770000 - 27115000
ntot: 2414
nspec: 301
k: 32
pearson0: 0.943013625
pearson3: 0.897944047

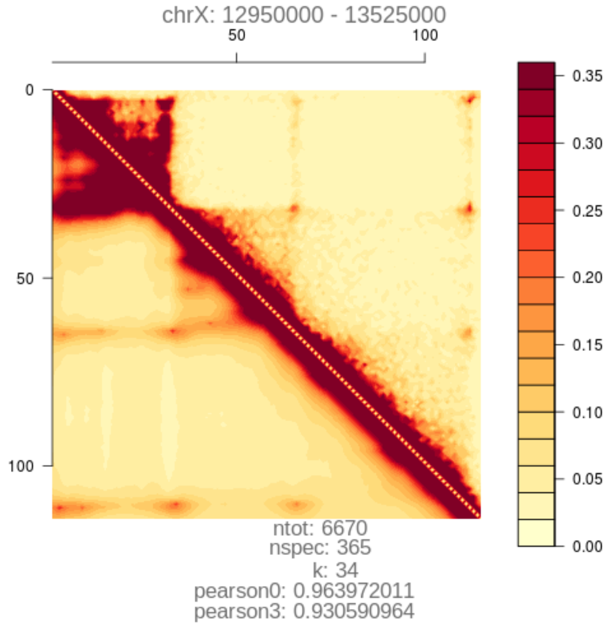
chrX: 12950000 - 13525000
ntot: 6670
nspec: 365
k: 34
pearson0: 0.963972011
pearson3: 0.930590964

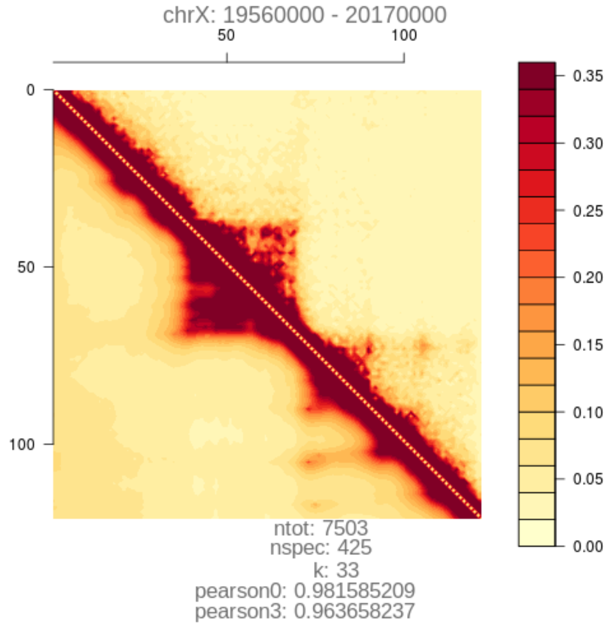
chrX: 19560000 - 20170000
ntot: 7503
nspec: 425
k: 33
pearson0: 0.981585209
pearson3: 0.963658237

#### 6.4 Figure S4.

Details of Pearson correlation between reconstructed ensembles and experimentally measured Hi-C of all loci, when a progressively increasing number of bands around the diagonal of the Hi-C heatmap are removed. For each locus, the $x$-axis shows the number of bands at intervals of 5 KB units around the diagonal that are removed, the $y$-axis shows the re-calculated Pearson correlation coefficient after removal.

**FIGURE S4**

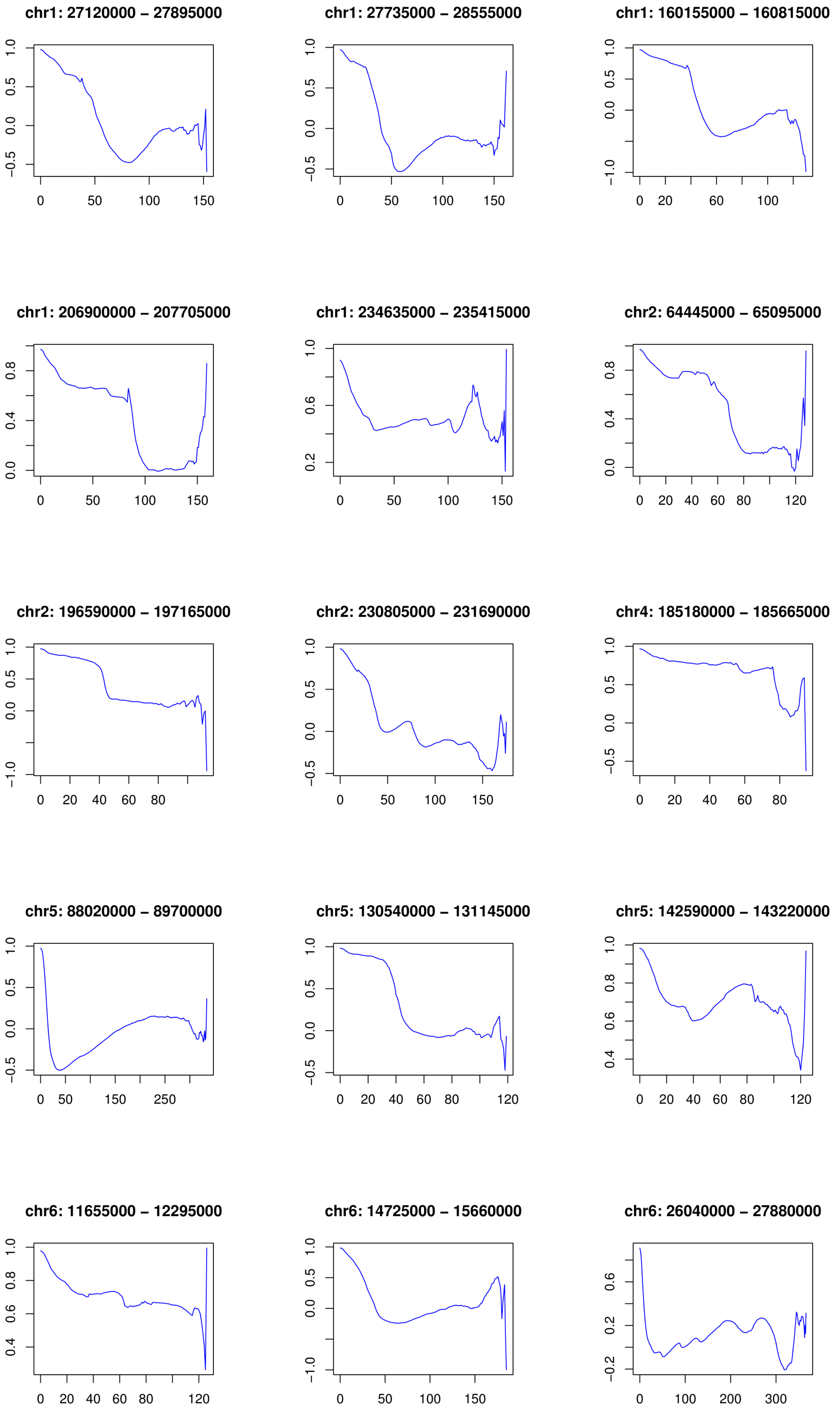
chr1: 27120000 – 27895000
chr1: 27735000 – 28555000
chr1: 160155000 – 160815000
chr1: 206900000 – 207705000
chr1: 234635000 – 235415000
chr2: 64445000 – 65095000
chr2: 196590000 – 197165000
chr2: 230805000 – 231690000
chr4: 185180000 – 185665000
chr5: 88020000 – 89700000
chr5: 130540000 – 131145000
chr5: 142590000 – 143220000
chr6: 11655000 – 12295000
chr6: 14725000 – 15660000
chr6: 26040000 – 27880000

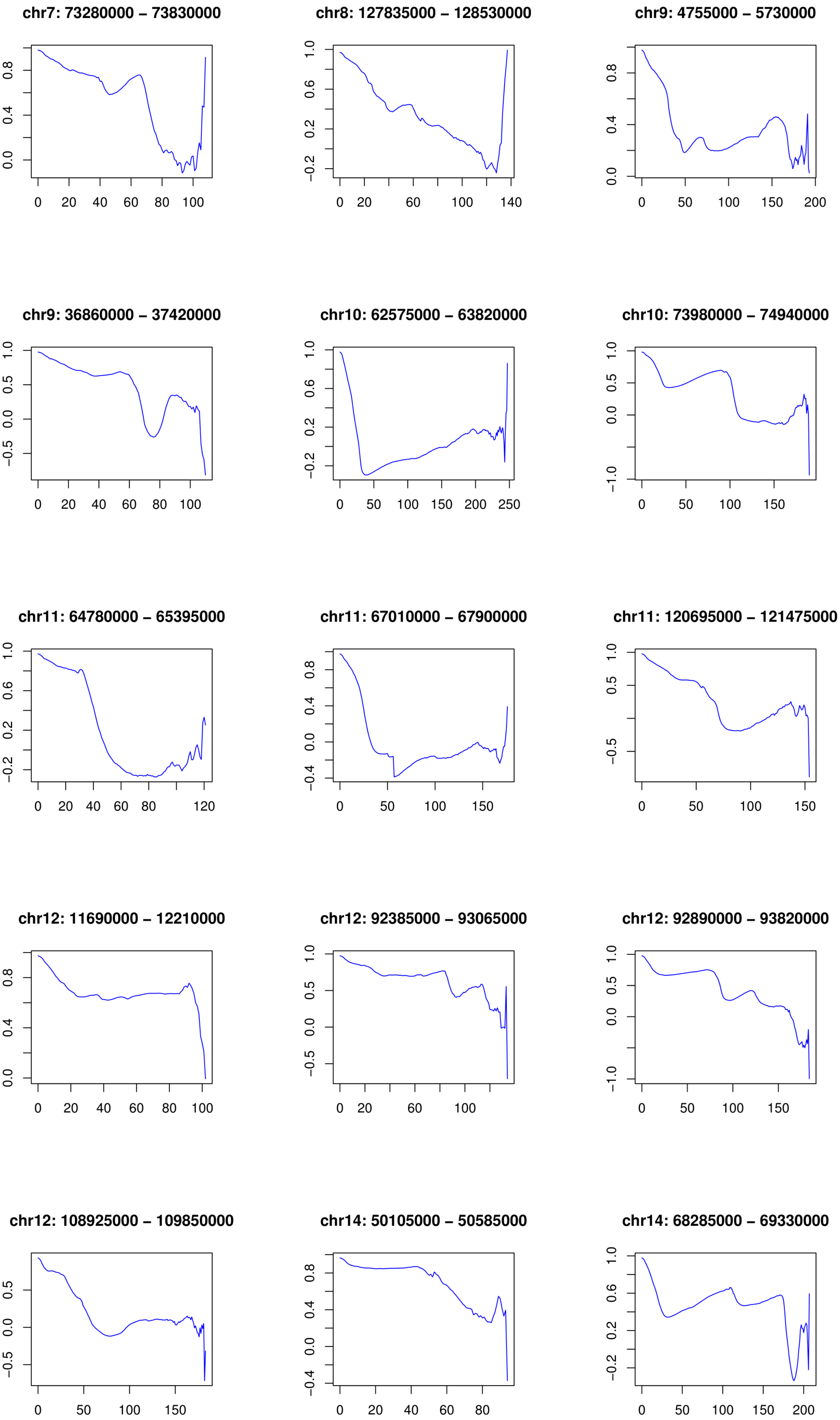

chr7: 73280000 – 73830000
0.0 0.4 0.8
0 20 40 60 80 100
chr8: 127835000 – 128530000
−0.2 0.2 0.6 1.0
0 20 60 100 140
chr9: 4755000 – 5730000
0.0 0.4 0.8
0 50 100 150 200
chr9: 36860000 – 37420000
−0.5 0.0 0.5 1.0
0 20 40 60 80 100
chr10: 62575000 – 63820000
−0.2 0.2 0.6 1.0
0 50 100 150 200 250
chr10: 73980000 – 74940000
−1.0 0.0 0.5 1.0
0 50 100 150
chr11: 64780000 – 65395000
−0.2 0.2 0.6 1.0
0 20 40 60 80 120
chr11: 67010000 – 67900000
−0.4 0.0 0.4 0.8
0 50 100 150
chr11: 120695000 – 121475000
−0.5 0.5 1.0
0 50 100 150
chr12: 11690000 – 12210000
0.0 0.4 0.8
0 20 40 60 80 100
chr12: 92385000 – 93065000
−0.5 0.0 0.5 1.0
0 20 60 100
chr12: 92890000 – 93820000
−1.0 0.0 0.5 1.0
0 50 100 150
chr12: 108925000 – 109850000
−0.5 0.0 0.5
0 50 100 150
chr14: 50105000 – 50585000
−0.4 0.0 0.4 0.8
0 20 40 60 80
chr14: 68285000 – 69330000
−0.2 0.2 0.6 1.0
0 50 100 150 200

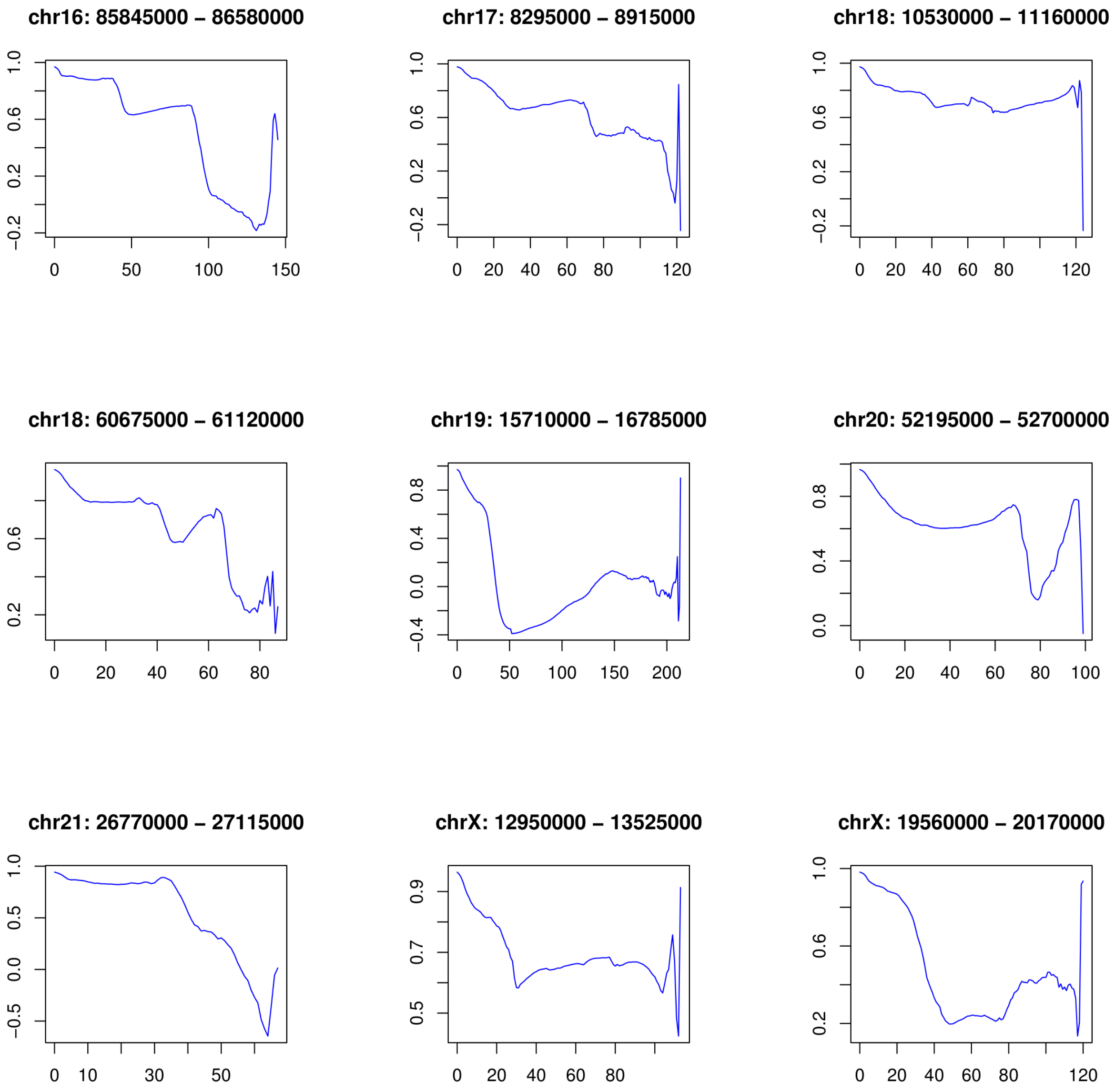
chr16: 85845000 – 86580000
chr17: 8295000 – 8915000
chr18: 10530000 – 11160000
chr18: 60675000 – 61120000
chr19: 15710000 – 16785000
chr20: 52195000 – 52700000
chr21: 26770000 – 27115000
chrX: 12950000 – 13525000
chrX: 19560000 – 20170000

#### 6.5 Figure S5.

**SPRITE coverage fractions for principal loops stratified by genomic distance span** for loci **(a)** chr14: 68,285,000 – 69,330,000 and **(b)** chr12: 11,690,000 – 12,210,000. Each dot is a specific (blue) or non-specific (pink) principal loop with corresponding LOESS smoothed trend lines.

#### Figure S5

**a**

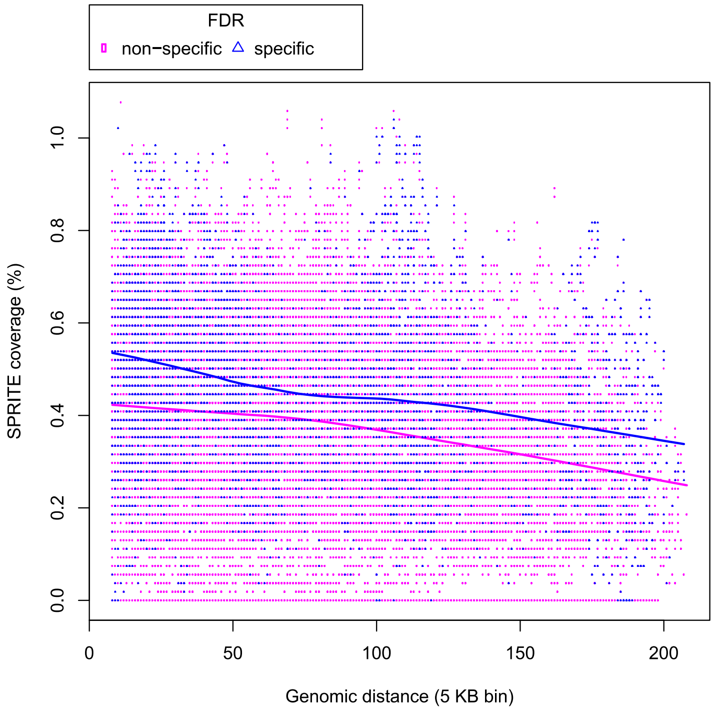
FDR
non−specific
specific
SPRITE coverage (%)
0.0
0.2
0.4
0.6
0.8
1.0
0
50
100
150
200
Genomic distance (5 KB bin)

**b**

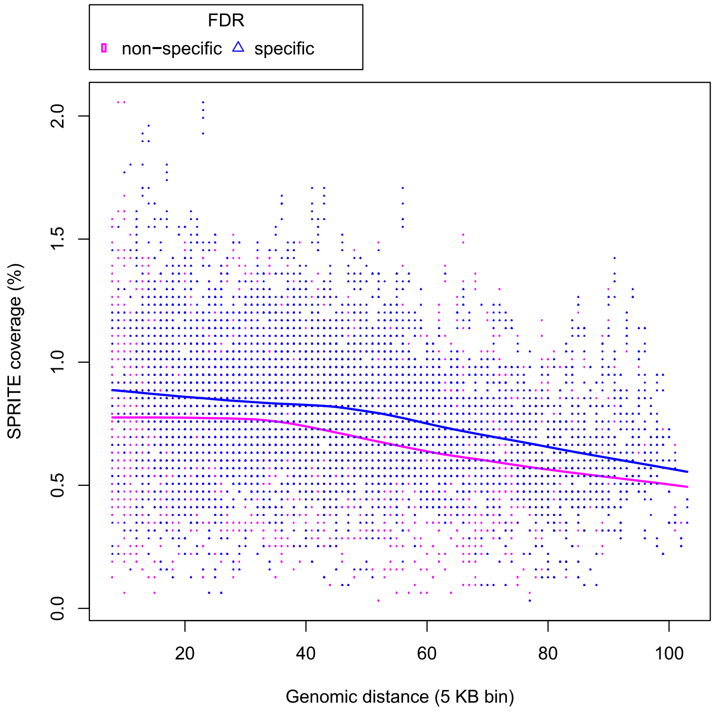
FDR
non−specific
specific
SPRITE coverage (%)
0.0
0.5
1.0
1.5
2.0
20
40
60
80
100
Genomic distance (5 KB bin)

#### 6.6 Figure S6.

**Box plots comparing proportions of functional associations for non-specific versus specific 3-body chromatin interactions across 39 modeled loci.** Comparative box plots for proportions of non-specific (grey, left) versus specific (black, right) for 3-bodies with **(a)** no functional association, **(b)** multiple super-enhancers ($\geq 2$) spatially interacting with a promoter, and **(c)** multiple super-enhancers ($\geq 3$) spatially co-interacting.

**Figure S6**

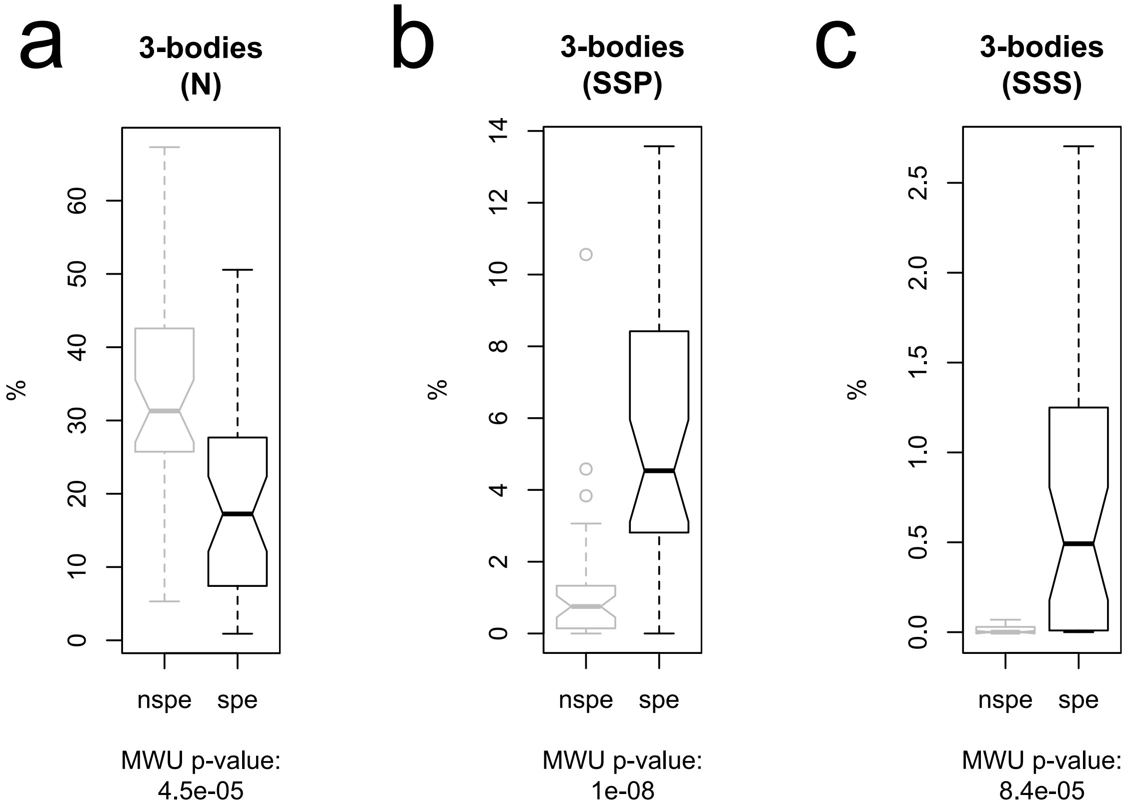

**Key**

- N – 3-body with no functional association
- SSP – 3-body among ≥ 2 Super-enhancers and a Promoter
- SSS – 3-body among ≥ 3 Super-enhancers
- nspe – non-specific
- spe – specific
- MWU – Mann-Whitney U

#### 6.7 Figure S7.

**Box plots comparing proportions of functional associations for non-specific versus specific principal loop chromatin interactions across 39 modeled loci.** Comparative box plots for proportions of non-specific (grey, left) versus specific (black, right) for principal loops with **(a)** multiple super-enhancers ($\geq 2$) spatially interacting, **(b)** a super-enhancer spatially interacting with a promoter.

**Figure S7**

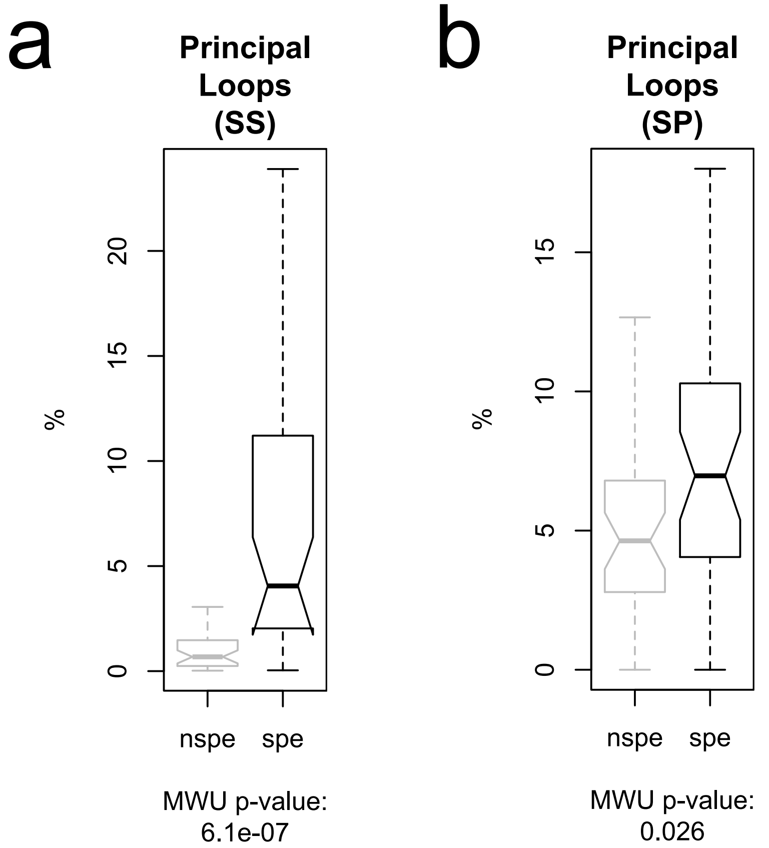

**Key**

- SS – Principal loop bridging ≥ 2 Super-enhancers
- SP – Principal loop bridging Super-enhancer to Promoter
- nspe – non-specific
- spe – specific
- MWU – Mann-Whitney U

#### 6.8 Figure S8.

**Comparison of principal loop participation (as z-score) for super-enhancer (SE) vs non-SE regions** across 39 modeled loci with **(a)** box plots with corresponding p-value for Mann-Whitney U test of median difference and **(b)** empirical cumulative distribution plots with corresponding p-value for Kolmogorov–Smirnov (KS) test. Note, yellow vertical line is the KS-statistic, defined as the maximum difference between the two distributions, and is used to determine the KS p-value.

**Figure S8**

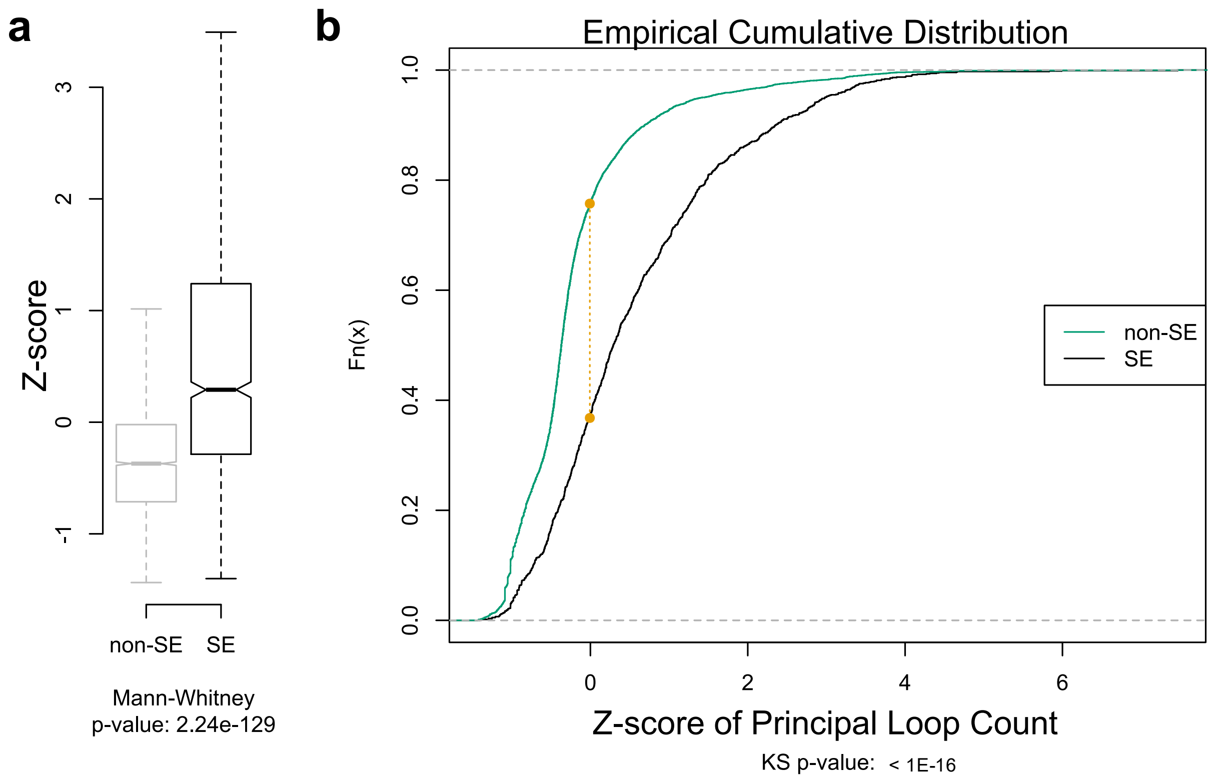
a
b
Z-score
3
2
1
0
-1
non-SE
SE
Mann-Whitney
p-value: 2.24e-129
Empirical Cumulative Distribution
Fn(x)
1.0
0.8
0.6
0.4
0.2
0.0
non-SE
SE
0
2
4
6
Z-score of Principal Loop Count
KS p-value: < 1E-16

### 7 Functional landscape category allocation

In the main text, the functional landscapes for 3-bodies and maximal many-bodies are depicted as sunburst plots. Since genomic regions are in units of 5 KB, a single 5 KB interval may contain multiple super-enhancer (SE), enhancer (E), and/or promoter (P) annotations; therefore, a many-body interaction featuring that 5 KB interval may qualify for multiple categories in the sunburst plots. To avoid double counting (*i.e.*, to maintain disjoint categories), each many-body interaction is allocated to the *single* category with the highest ranking. For 3-body interactions, the ranking rule is according to: SE-SE-P > SE-SE-SE > SE-SE-E > SE-E-P > SE-E-E > SE-P-P > E-E-P > E-E-E > E-P-P > P-P-P > SE-P-N > E-P-N > P-P-N > SE-SE-N > SE-E-N > E-E-N > SE-N-N > E-N-N > P-N-N > N-N-N; where N stands for no known SE, E, or P annotation. For principal loops of maximal many bodies, the ranking rule is according to: SE-P > E-P > P-P > SE-SE > SE-E > E-E > SE-N > E-N > P-N > N-N; where N is defined as before. These rules are applied separately to both specific and non-specific interactions. Note, the pie charts are simply taken as the innermost ring of the corresponding sunburst charts, and therefore also feature disjoint allocations of the many-body interactions.

### 8 Tables

#### 8.1 Table S1.

Details of the genomic loci, measured Hi-C, and simulated Hi-C. These include genomic locations of loci, annotated genes within the loci, statistics of Hi-C measurements, number of their specific calls, and sizes of coarse-grained specific contacts used for simulation. Canonical and distance corrected Pearson correlations [1] between simulated and measured Hi-C contacts are also reported. For loci: chr 1: 234635000–235415000, chr 9: 4755000–5730000, chr 12: 11690000–12210000, chr 14: 68285000–69330000, Pearson correlations are averages of two replicated simulations.

The columns are:

```
name - Region name in format <chr>.<bp_start>.<bp_end>
chr - Chromosome
bp_start - Start base pair
bp_end - End base pair
bp_span - Genomic span of locus in base pairs
refgene - Annotated genes within locus
mon_diam - Diameter of monomer bead in Angstroms
mon_num - Number of monomer beads used
ntot - Total number of observed (measured) Hi-C contacts
nspc - Total number of called specific Hi-C contacts
k - Number of retained specific contacts used in modeling
kf - Fraction of specific contacts retained (k/nspc)
fdr - FDR threshold for calling specific
pearsonDC - Distance corrected Pearson correlation between
            measured and simulated Hi-C
pearson0 - Pearson correlation between measured and
           simulated Hi-C with all diagonals retained
pearson1 - Pearson correlation but with 0-diagonal removed
pearson2 - Pearson correlation but with 0- and 1- diagonals removed
pearson3 - Pearson correlation but with 0-, 1-, and 2- diagonals removed
pearsonFHC - Pearson correlation at Fit-Hi-C significant contacts
nFHC - Number of Fit-Hi-C significant contacts at q-value < 1%
A - Fraction of locus designated as A compartment (100 KB bin)
B - Fraction of locus designated as B compartment (100 KB bin)
```

Note, some loci have missing Hi-C regions and therefore: $A + B \leq 1.0$

Table S1 also contains information on the correlation between simulated and experimental Hi-C data at contact locations identified as statistically significant by the Fit-HiC method ($q < 1\%$). After excluding 2 loci which have no significant contacts by Fit-Hi-C, the mean number of significant contacts per locus is $\sim 112 \pm 42$. The mean Pearson correlation is $\sim 0.97 \pm 0.03$. Similar results are obtained with $q < 5\%$ and $q < 10\%$.

#### 8.2 Table S2.

Details of biological markers used for the predictive models of enrichment of anchors of principal loops of specific many-body interactions.

The columns are:

```
Genome Build - Human genome assembly version
Technique - Experimental technique used for capturing biomarker
Target - Biomarker targeted by experimental technique
Replicate - Experimental replicate number
ENCODE - ENCODE accession number.
```

#### 8.3 Table S3.

LILY [2] super-enhancer (SE), enhancer, and promoter functional annotations.
The columns are:

```
chrom - Chromosome
chromStart - Start base pair
chromEnd - End base pair
label - Annotation, one of: enhancer, promoter, or SE
score - LILY score based on H3K27ac ChIP-seq signal
strand - Always '+' for plus strand
```

[5] Bhaskar DasGupta and Jie Liang. *Models and Algorithms for Biomolecules and Molecular Networks*. John Wiley & Sons, 2016.

[6] Andrew Gelman, Hal S Stern, John B Carlin, David B Dunson, Aki Vehtari, and Donald B Rubin. *Bayesian data analysis*. Chapman and Hall/CRC, 2013.

[7] Charles J Geyer. Markov chain monte carlo lecture notes. *Course notes, Spring Quarter*, 1998.

[8] Upton Graham and Ian Cook. *Understanding statistics*. Oxford University Press, 1997.

[9] Gamze Gürsoy, Yun Xu, Amy L Kenter, and Jie Liang. Spatial confinement is a major determinant of the folding landscape of human chromosomes. *Nucleic acids research*, 42(13):8223–8230, 2014.

[10] Gamze Gürsoy, Yun Xu, Amy L Kenter, and Jie Liang. Computational construction of 3d chromatin ensembles and prediction of functional interactions of alpha-globin locus from 5c data. *Nucleic acids research*, 45(20):11547–11558, 2017.

[11] Gamze Gürsoy, Yun Xu, and Jie Liang. Spatial organization of the budding yeast genome in the cell nucleus and identification of specific chromatin interactions from multi-chromosome constrained chromatin model. *PLoS computational biology*, 13(7):e1005658, 2017.

[12] David Heckerman, Dan Geiger, and David M Chickering. Learning bayesian networks: The combination of knowledge and statistical data. *Machine learning*, 20(3):197–243, 1995.

[13] NORA Helfand. Polya's urn and the beta-bernoulli process.

[14] Daphne Koller and Nir Friedman. *Probabilistic graphical models: principles and techniques*. MIT press, 2009.

[15] M. Lin, J. Zhang, H. M. Lu, R. Chen, and J. Liang. Constrained proper sampling of conformations of transition state ensemble of protein folding. *J Chem Phys*, 134(7):075103, Feb 2011.

[16] Jun S Liu. *Monte Carlo strategies in scientific computing*. Springer Science & Business Media, 2008.

[24] Erik Blaine Sudderth. *Graphical models for visual object recognition and tracking*. PhD thesis, Massachusetts Institute of Technology, 2006.

[25] Longzhi Tan, Dong Xing, Chi-Han Chang, Heng Li, and X Sunney Xie. Three-dimensional genome structures of single diploid human cells. *Science*, 361(6405):924–928, 2018.

[26] K Vijaymeena, M. K.; Kavitha. A survey on similarity measures in text mining. *Machine Learning and Applications: An International Journal*, 3(1):19–28, 2016.
